# Nucleotide depletion promotes cell fate transitions by inducing DNA replication stress

**DOI:** 10.1101/2022.08.16.503984

**Authors:** Brian T. Do, Peggy P. Hsu, Sidney Y. Vermeulen, Zhishan Wang, Taghreed Hirz, Keene L. Abbott, Najihah Aziz, Joseph M. Replogle, Stefan Bjelosevic, Jonathan Paolino, Samantha Nelson, Samuel Block, Alicia M. Darnell, Raphael Ferreira, Hanyu Zhang, Jelena Milosevic, Daniel R. Schmidt, Christopher Chidley, Isaac S. Harris, Jonathan S. Weissman, Yana Pikman, Kimberly Stegmaier, Sihem Cheloufi, Xiaofeng A. Su, David B. Sykes, Matthew G. Vander Heiden

**Author notes:** These authors contributed equally to this work (Brian T. Do, Peggy P. Hsu).

## Abstract

Control of cellular identity requires coordination of developmental programs with environmental factors such as nutrient availability, suggesting that modulating aspects of metabolism could alter cell state along differentiation trajectories. Here we find that nucleotide depletion and DNA replication stress are common drivers of cell state progression across a variety of normal and transformed hematopoietic systems. DNA replication stress-induced cell state transitions begin during S phase and are independent of ATR/ATM checkpoint signaling, double-stranded DNA break formation, and changes in cell cycle length. In systems where differentiation is blocked by oncogenic transcription factor expression, replication stress leads to increased activity at primed regulatory loci and expression of lineage-appropriate maturation genes while progenitor TF activity is still present. Altering the baseline cell state by manipulating the cohort of transcription factors expressed redirects the effect of replication stress towards induction of a different set of lineage-specific genes. The ability of replication stress to selectively activate primed maturation programs across different cellular contexts suggests a general mechanism by which metabolism can promote lineage-appropriate and potentially therapeutically relevant cell state transitions.

## INTRODUCTION

Mammalian cells must tightly regulate cell identity to ensure proper development and maintenance of tissue homeostasis (Intlekofer and Finley, 2019; Shapira and Christofk, 2020). Diverse external cues can influence cell type-specific developmental programs, ultimately affecting whether cells self-renew or differentiate towards a particular identity defined by their epigenetic and transcriptional state. Dysregulation of cellular identity lies at the root of a wide range of processes including cancer initiation and progression, underscoring the therapeutic potential of novel strategies to deterministically drive changes in cell fate (Le Magnen et al., 2018; Quintanal-Villalonga et al., 2020; Roy and Hebrok, 2015).

It is increasingly recognized that the set of external cues driving cell fate decisions includes not only ligands that activate developmental signaling pathways (Perrimon et al., 2012), but also the availability of environmental nutrients (Baksh and Finley, 2021; Lin et al., 2018). Metabolite profiling and systematic identification of metabolic vulnerabilities in different cell types or stages of differentiation across various systems have demonstrated that nutrient availability can not only gate, but also influence, lineage decisions (Gonzalez-Menendez et al., 2021; Oburoglu et al., 2014; Sharpley et al., 2021; Solmonson et al., 2022). While the mechanistic connections between metabolism and cell fate change remain unclear, some metabolic perturbations have been found to directly alter the levels of metabolites that act as substrates or cofactors for enzymes that control epigenetic marks on chromatin (Baksh and Finley, 2021; Meier, 2013). Intriguingly, the same perturbations can often promote or suppress lineage progression depending on the cell type and environmental context (Carey et al., 2015; TeSlaa et al., 2016), and while it is clear how impacting the activity of enzymes that influence epigenetic state might facilitate transitions between gene expression states, a comprehensive understanding of how changes in metabolism can promote specific cell fate transitions in different cell types is lacking.

Hematopoiesis provides a tractable system to dissect the contribution of metabolism to cell state and differentiation as the hierarchical structure and molecular regulators of lineage commitment and progression have been comprehensively cataloged (Bao et al., 2019). Moreover, many hematopoietic disorders, such as genetically inherited blood diseases and malignancies, involve well-characterized defects in differentiation. In acute myeloid leukemia (AML), a cancer defined by proliferation of undifferentiated myeloid precursors (Döhner et al., 2017), differentiation blockades are maintained by a variety of genetic alterations that reduce the activity of lineage-determining transcription factors (TFs) or constitutively activate progenitor-specific regulatory networks (Papaemmanuil et al., 2016). When paired with mutations that promote increased proliferation, hematopoietic stem and progenitor cells are ultimately arrested in an immature state of self-renewal. Targeted therapies that relieve the differentiation blockade have been transformative in the treatment of select genetically defined subsets of AML, such as acute promyelocytic or IDH-mutant leukemias, yet broad strategies to overcome differentiation arrest are still lacking for most other AML subsets as well as solid tumors (de Thé, 2018; Wang et al., 2021).

Inhibitors of the pyrimidine biosynthetic enzyme dihydroorotate dehydrogenase (DHODH) were previously identified as a novel class of differentiation agents in non-promyelocytic and non-IDH mutated AML (Sykes et al., 2016). As a result, DHODH inhibitors may be effective for the treatment of AML and other cancers (Christian et al., 2019; Zee et al., 2021). Nevertheless, the mechanism by which pyrimidine depletion can induce differentiation remains unclear, and whether perturbing other metabolic enzymes and pathways might also influence lineage-specific gene expression changes in cells remains an open question.

Using metabolism-focused pharmacologic and genome-wide genetic screens, we find that perturbations that lower dNTP levels and interfere with DNA replication, lead to replication stress and drive cell state changes along differentiation trajectories. Our findings are generalizable across a variety of hematopoietic cell systems, including normal and malignant and those of different blood cell lineages and oncogenic backgrounds. We dissect the role of various sequelae of replication stress, including ATR/ATM checkpoint signaling, cell cycle lengthening, and DNA double-stranded breaks, and find that differentiation is largely independent of these processes. At the chromatin level, replication stress primarily activates primed regulatory genomic loci during S phase to induce expression of a more differentiated transcriptional program even in the presence of progenitor transcription factor activity. As a result, altering the baseline chromatin state correspondingly shifts the transcriptional effects of replication stress. Our integrative analysis of hematopoietic cell state transitions upon nucleotide depletion and DNA replication stress add to accumulating evidence in other cellular contexts that DNA replication and, by extension, nucleotide metabolism are tightly intertwined with maintenance of cellular identity.

## RESULTS

### Perturbing nucleotide metabolism is a route to myeloid maturation in ER-Hoxa9 cells

To explore how changes in metabolism can impact cell identity, we first took advantage of a model of hematopoietic differentiation blockade where maturation of murine granulocyte-macrophage progenitor (GMP) cells is blocked by activity of a ER-Hoxa9 (estrogen receptor-homeobox transcription factor A9) fusion protein driven by exposure to estradiol (E2) (**Figure 1A**) (Sykes et al., 2016). Hoxa9, a progenitor transcription factor that is normally downregulated during myeloid differentiation, is overexpressed in over half of AMLs driven by a variety of genetic alterations, underscoring the clinical relevance of this model system (Collins and Hess, 2016). Upon E2 withdrawal or pharmacologic antagonism of the estrogen receptor, ER-Hoxa9 activity is curtailed and the ER-Hoxa9 GMP cells terminally differentiate into neutrophils (Christian et al., 2019; Sykes et al., 2016). This model system is also engineered to express green fluorescent protein (GFP) from the lysozyme M (*Lyz)* locus as a marker of neutrophil maturation, enabling screening for compounds that promote differentiation; a single-dose chemical screen involving a largely unannotated small molecule library previously found that inhibitors of dihydroorotate dehydrogenase (DHODH), an enzyme required for the pyrimidine synthesis, could promote differentiation (Sykes et al., 2016).

**Figure 1.**
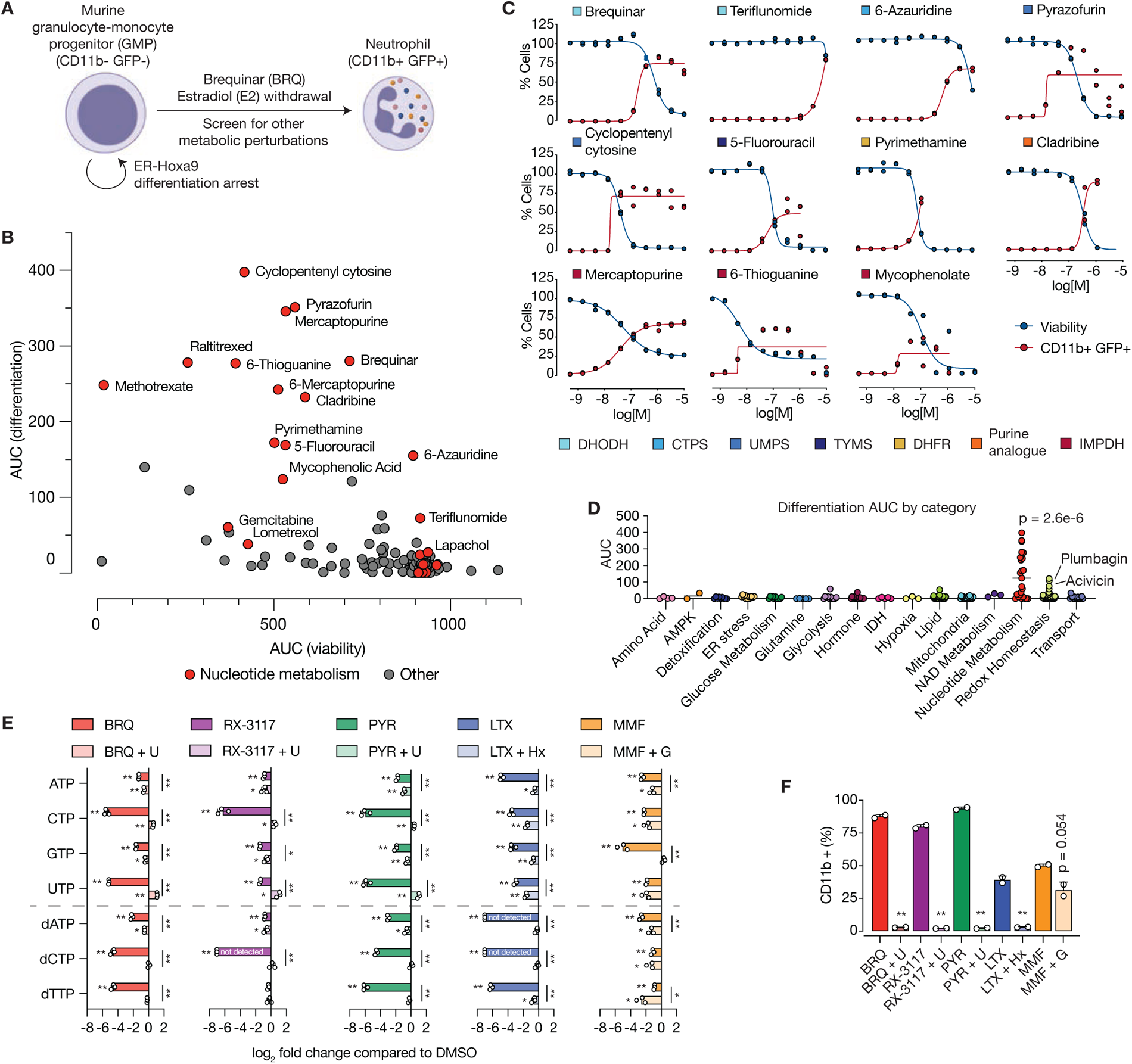
A focused pharmacologic screen reveals molecules targeting nucleotide metabolism can promote leukemia differentiation. A. Schematic of mouse ER-Hoxa9 granulocyte-monocyte progenitor (GMP) system. Treatment with brequinar (BRQ) and withdrawal of estrogen (-E2) to suppress ER-Hoxa9 activity are known inducers of differentiation towards the neutrophil state, and the neutrophil markers CD11b and Lysozyme-GFP can be monitored to assess the extent of lineage progression. A screen was performed for other metabolic perturbations that could induce differentiation. B. Plot showing area under the curve (AUC) for CD11b/GFP expression (differentiation) and viability for ER-Hoxa9 cells in response to all drugs in the metabolic drug library (Harris et al., 2019). Cells were treated for 4 days with ten doses of each drug up to 10 μM (see Methods). C. Percent viability (blue) and differentiation as assessed by CD11b and GFP expression (red) for ER-Hoxa9 GMP cells treated with each of the indicated drugs. Data from two technical screening replicates are plotted. The enzyme target of each drug is color coded as indicated. DHODH, dihydroorotate dehydrogenase. CTPS, cytidine triphosphate synthetase. UMPS, uridine monophosphate synthetase. TYMS, thymidylate synthetase. DHFR, dihydrofolate reductase. IMPDH, inosine monophosphate dehydrogenase. D. Area under the curve (AUC) for CD11b/GFP expression (differentiation) for each drug in the metabolic drug screen, grouped by category of drug target. P-values were calculated using a Mann-Whitney U test for each category compared to the distribution of all AUC values. E. Log2 fold change in levels of NTPs and dNTPs in ER-Hoxa9 GMP cells following 24h of treatment with the indicated drug (all at 1 μM), with or without rescue by addition of the indicated cognate nucleotides (all at 1 mM). Log fold change values are compared to the median of the untreated normalized area under the peak. Some metabolites were not detected due to low abundance. BRQ, brequinar. U, uridine. PYR, pyrazofurin. LTX, lometrexol. MMF, mycophenolate mofetil. Hx, hypoxanthine. G, guanine. U, uridine. (*), p < 0.05; (**), p < 0.01, and p-values were calculated using a Student’s t-test on log fold change values for each treatment compared to DMSO treatment (denoted to the left of the bars), or for each treatment-rescue pair (denoted to the right of the bars). F. Percentage of CD11b-expressing ER-Hoxa9 GMP cells following 96h of treatment with the indicated purine synthesis inhibitors (LTX at 41 nM, MMF at 370 nM) or pyrimidine synthesis inhibitors (BRQ at 1.1 μM, RX-3117 at 1.1 μM, PYR at 123 nM), with or without addition of cognate nucleotides (all at 1 mM). p < 0.05, (**), p < 0.005, or as indicated, and p-values were calculated using a Student’s t-test on samples with and without nucleotides. Abbreviations are as in (E).

To identify other metabolic perturbations that can promote differentiation, we used the ER-Hoxa9 GMPs to perform a targeted flow cytometry screen with a curated library of 240 small molecule inhibitors of various classes of metabolic enzymes, where cells were treated over a 10-point dose range of each compound (**Figure 1B-C, Table S1)** (Harris et al., 2019). Differentiation was assessed with GFP fluorescence and expression of the myeloid surface marker CD11b, and viability was assessed using forward and side scatter. Among compounds that were found to promote both GFP and CD11b expression along with retained cell viability after 96 hours of treatment, the majority targeted nucleotide metabolism (**Figure 1B-D**), including the DHODH inhibitors brequinar (BRQ) and leflunomide as well as additional hits that targeted either pyrimidine synthesis (such as cyclopentenyl cytosine, 5-fluorouracil, and pyrazofurin) or purine synthesis (such as mycophenolic acid, mercaptopurine, and 6-thioguanine). Differentiation was also induced by antifolates (pyrimethamine, methotrexate, and raltitrexed), which inhibit folate-requiring enzymes in purine and/or thymidine synthesis, and nucleoside analogues (cytarabine, cladribine, and 5-fluorouracil) that can both incorporate into nucleic acid polymers and inhibit various nucleotide biosynthetic and salvage enzymes. Importantly, many of these compounds were able to induce differentiation markers at doses that only partially reduced cell viability (**Figure 1C**).

Differentiation-inducing compounds that were not primarily annotated as nucleotide metabolism inhibitors included acivicin, a glutamine analog that could inhibit nucleotide biosynthesis (Kensler et al., 1982), and plumbagin, which has been shown to inhibit pyrimidine biosynthesis (Guan et al., 2021) (**Figure 1D**). In contrast to compounds that target nucleotide metabolism, most other metabolic inhibitors tested had no effect on differentiation marker expression (**Figure 1D**). Indeed, inhibitors of the mitochondrial electron transport chain (rotenone and oligomycin), NAD+ biosynthesis (FK866 and GPP78), ER stress (CCF642, thapsigargin, and tunicamycin), and lipid synthesis (GSK2194069) did not induce differentiation markers in these cells **(Figure S1A)**. For rotenone, we also did not observe induction of differentiation markers at an earlier timepoint where reduced proliferation rate, but not toxicity, was observed **(Figure S1B)**. These data indicate that generalized metabolic stresses or decreased proliferation rate are not sufficient to initiate cell state progression.

To assess the metabolic effects of nucleotide synthesis inhibitors that affect differentiation marker levels, we performed liquid chromatography-mass spectrometry (LC-MS) on metabolites extracted from ER-Hoxa9 cells treated with various inhibitors of *de novo* purine or pyrimidine nucleotide biosynthesis. Treating cells with compounds that affect pyrimidine metabolism (brequinar, RX-3117, and pyrazofurin) and purine metabolism (MMF and lometrexol) led to metabolic changes characterized by depletion of the respective classes of nucleotide precursors, nucleosides, ribonucleotides, and deoxyribonucleotides (**Figure 1E, Figure S1C-D, Table S2)**. Inhibition of either purine or pyrimidine nucleotide synthesis also led to changes in levels of glycolytic and TCA cycle intermediates including phosphoenolpyruvate, alpha-ketoglutarate, and citrate, likely reflecting links between nucleotide anabolism and other aspects of cell metabolism **(Figure S1C-D)**. These changes were broader than those observed following estrogen withdrawal, where the levels of only nine metabolites were significantly altered **(Figure S1E, Table S2)**. Co-treatment with nucleotide synthesis inhibitors and the appropriate nucleobases or nucleosides normalized NTP and dNTP levels. Notably, this rescued differentiation and restored metabolic changes, arguing that these small molecules promote differentiation by perturbing nucleotide metabolism (**Figure 1E-F, Figure S1C-D, Table S2)**.

### Replication stress links nucleotide depletion to differentiation

Nucleotides play a central role in multiple cellular processes, from DNA replication to transcription to protein glycosylation (Mullen and Singh, 2023; Shi et al., 2023). Given the effect of nucleotide synthesis inhibitors on dNTP levels, we first wanted to understand whether their effects on DNA replication might be contributing to the induction of differentiation markers by these agents. To test this, we directly inhibited ribonucleotide reductase (RNR), the enzyme necessary to synthesize dNTPs for DNA replication, using hydroxyurea (HU), gemcitabine, clofarabine, or 3-AP. We found that all of these RNR inhibitors promoted differentiation of ER-Hoxa9 cells, and we confirmed that HU treatment affected dNTP levels without affecting NTP levels, in contrast to BRQ, where both dNTPs and NTPs were depleted (**Figure 2A-B, S2A, Table S2)**. To assess whether an impact on DNA replication might be involved in promoting differentiation downstream of altered dNTP levels, we directly inhibited DNA polymerase using aphidicolin (APH) and found that this also led to differentiation without reducing dNTP or NTP levels, consistent with prior findings in other AML models (Griffin et al., 1982) (**Figure 2A-B).** Metabolite changes induced by HU and APH were less extensive than those induced by BRQ and mostly confined to nucleotides and their precursors **(Figure S2B)**. In addition to inducing maturation-related changes in surface marker expression, BRQ, HU, and APH led to comparable morphological changes consistent with differentiation, such as granule formation, an increased cytoplasmic to nuclear ratio, and ringed or multilobed nuclei (**Figure 2C**). These data suggest a specific role of impaired DNA replication in mediating the effect of nucleotide synthesis inhibitors on cell state.

**Figure 2.**
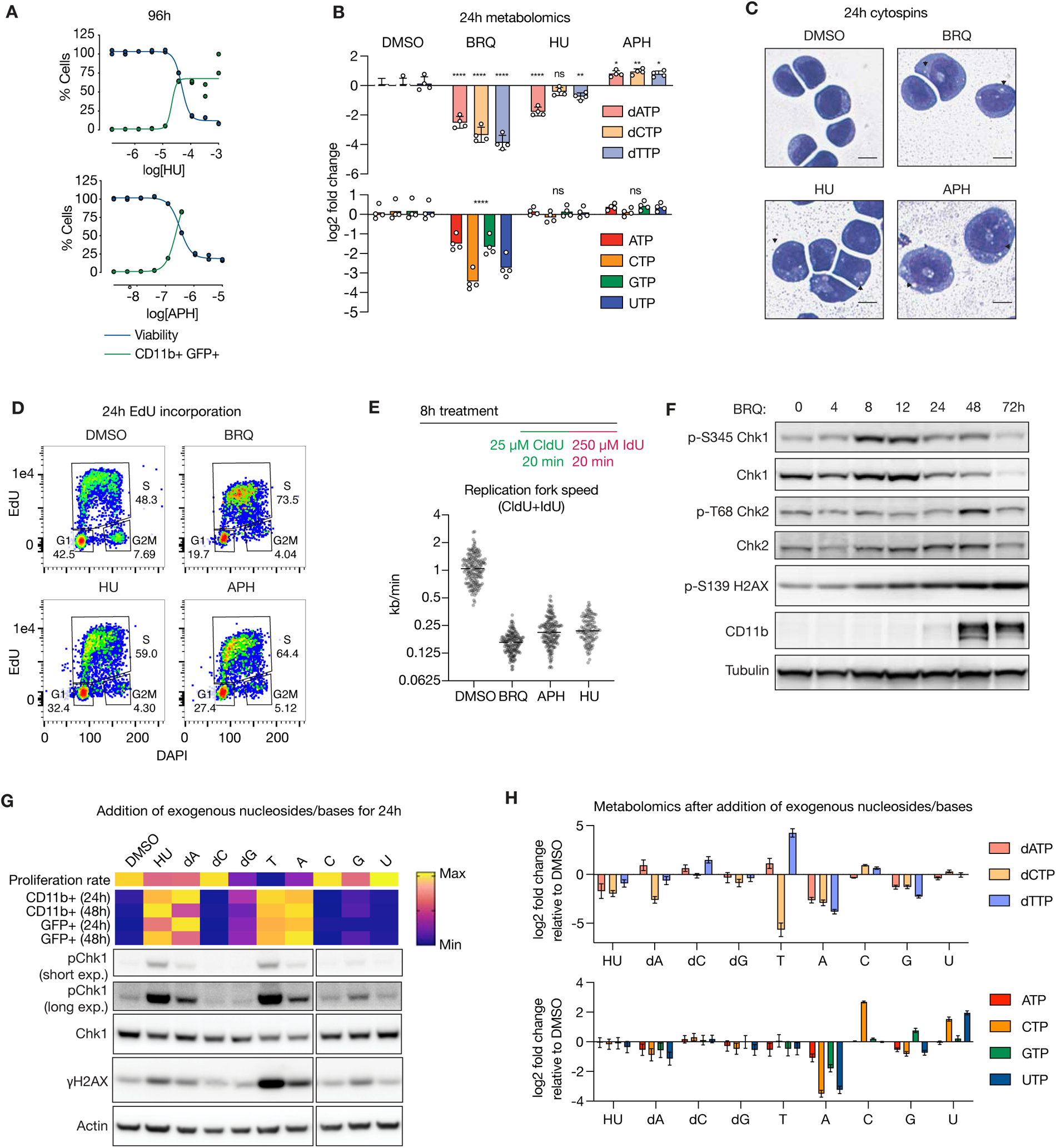
Replication stress links nucleotide depletion and dNTP imbalance to cell state progression. A. Percent viability (blue) and differentiation, as assessed by CD11b and GFP expression (green), for ER-Hoxa9 cells after treatment with the indicated doses of HU or APH for 96 hours (n=2). B. Log fold change of intracellular levels of dATP, dCTP, or TTP (top) and ATP, CTP, GTP, or UTP (bottom) in ER-Hoxa9 following 24h of treatment with DMSO, 1 μM BRQ, 50 μM HU, or 1 μM APH (n=4). (ns) not significant, (*) p < 0.05, (**) p < 0.01, (***) p < 0.001, and (****) p < 0.0001, and adjusted p-values were calculated using a two-way ANOVA with the Šidák correction for multiple comparisons. C. Cytospin images of ER-Hoxa9 cells treated with DMSO, 1 μM BRQ, 50 μM HU, or 1 μM APH for 24 hours. Granules are indicated with black arrowheads. Scale bars represent 10 μm. D. Representative flow cytometry plots (EdU-AF647 vs. DAPI) in ER-Hoxa9 cells treated with DMSO, 1 μM BRQ, 50 μM HU, or 1 μM APH for 24 hours then pulsed with EdU for the last 30 minutes. E. (top) Schematic of DNA fiber experiment. (bottom) Replication fork speed (kb/min) for ER-Hoxa9 cells treated with DMSO, 1 μM BRQ, 50 μM HU, or 1 μM APH for 8 hours. At least 100 fibers were assayed for each condition. F. Immunoblot for ER-Hoxa9 cells treated with 1 µM BRQ for the indicated times. G. Proliferation rates, surface marker expression, and immunoblots for ER-Hoxa9 cells treated for 24 hours (or 48 hours as indicated for CD11b and GFP measurements) with DMSO, 50 µM HU, or the indicated nucleobase/nucleoside (all at 1 mM except for dG, which was supplemented at 100 µM). Proliferation rates and percentage of CD11b+ and GFP+ cells are row-normalized. Immunoblots are cropped from the same blot. H. log2 relative abundance of dNTPs (top) and NTPs (bottom) for ER-Hoxa9 cells treated for 24h as in (G), compared to DMSO control (n=3, mean +/-SD).

While many of our prior differentiation marker measurements were conducted four days after induction of drug treatment, we next examined when the differentiation program was first induced. CD11b surface marker and lysozyme-GFP expression were detectable by flow cytometry within 4-8 hours of drug treatment **(Figure S2C)**. Importantly, while proliferation rate was strongly inhibited, CD11b and GFP expression were not necessarily associated with toxicity, as lower doses of BRQ and APH that induced substantial differentiation at 24 hours did not increase the percentage of dead cells, whereas higher doses could induce a more complete differentiation but at the expense of cell viability **(Figure S2D)**. Moreover, CD11b-positive cells sorted 24 hours after BRQ or APH treatment were able to restart proliferation and lose surface marker expression upon withdrawal of drug, indicating that the cell state maturation program induced by these drugs is reversible and separable from stresses that lead to cell death **(Figure S2E)**.

Decreased dNTP levels or inhibition of DNA polymerase lead to slower replication fork progression and reduced replication fidelity as part of a set of events broadly termed DNA replication stress (Zeman and Cimprich, 2014). We found that BRQ, HU, and APH all reduced the rate of 5-ethynyl-2-deoxyuridine (EdU) incorporation in S phase cells, indicating a slowdown in total DNA replication, and increased the proportion of cells in S phase (**Figure 2D, S2F-G)**. Using DNA fiber analysis, we measured the speed of single replication forks. At doses that drove differentiation, BRQ, HU, and APH all slowed forks at least 4-fold (**Figure 2E**). Accordingly, multiple indicators of DNA replication stress were evident upon drug treatment, including phosphorylation of histone H2AX at serine 139 (γH2AX) and the ATR target Chk1 at serine 345 (**Figure 2F, S2H)**. These signals were maximally induced between 8 and 12 hours after drug treatment, coinciding with when surface marker expression was first detectable, and were not induced by direct ER-Hoxa9 antagonism with fulvestrant. Consistent with these findings, an increase in chromatin-bound RPA was also observed at both 8 and 24 hours, and increased single-stranded DNA breaks were detected by the alkaline comet assay **(Figure S2I-K)** (Saxena and Zou, 2022).

One consequence of unresolved or unrestrained DNA replication stress is DNA double-stranded breaks (DSBs), whose sensing and repair are tightly controlled; an excess of DSBs can lead to replication catastrophe and/or apoptosis. However, BRQ did not lead to increased phosphorylation of Chk2, a downstream effect of DSBs, until 48 hours after treatment (**Figure 2F**). Phosphorylation of RPA at serine 4/8, a target of DNA-PK and ATM, was only mildly increased by BRQ, HU, and APH at 8 and 24 hours, in contrast to induction of high S4/8 phosphorylation upon treatment with a ten-fold higher dose of HU **(Figure S2L-M)**. These findings suggest that DSBs or replication catastrophe are not prerequisites for differentiation markers to be induced.

Previous work has demonstrated that supplementation of cells with exogenous nucleosides can also induce DNA replication stress by causing an imbalance of dNTPs, likely as a consequence of the exquisite feedback loops that maintain balanced dNTP synthesis (Diehl et al., 2022). Thus, to test whether exogenous nucleosides or nucleobases could promote expression of a maturation program, we treated ER-Hoxa9 cells with individual deoxyribonucleosides, ribonucleosides, or nucleobases and assessed CD11b and GFP expression, viability, proliferation, replication stress, and metabolite levels. Dose titrations were first performed with a maximum concentration of 1 mM, and deoxyadenosine (dA), thymidine (T), and adenine (A) induced the largest changes in differentiation markers **(Figure S2N)**. Evidence of differentiation was not observed for cytidine (C) or uridine (U) supplementation even at concentrations up to 10 mM; deoxyguanosine (dG) and guanosine (G) promoted subtle amounts of differentiation, but testing higher concentrations was limited by metabolite insolubility or toxicity. At doses of dA, T, and A where CD11b and GFP expression were induced without significant toxicity, significant Chk1 and H2AX phosphorylation was observed at 24 hours (**Figure 2G**). Using LC-MS, we found that these three treatments strongly depleted intracellular dNTP levels with variable effects on NTPs, consistent with the presence of DNA replication stress (**Figure 2H, Table S2)**. On the other hand, C, dC, and U addition did not cause replication stress or dNTP depletion, consistent with the lack of differentiation marker induction, and G and dG addition led to low to intermediate levels of replication stress and dNTP depletion congruent with their mild differentiation effects. In general, these data suggest that progression towards a more differentiated cell state is observed when supplementation of exogenous nucleosides/bases leads to dNTP depletion and DNA replication stress.

### Replication stress drives cell state changes in AML models and promotes normal erythroid differentiation

The ER-Hoxa9 GMP cells used for our initial studies are untransformed, as their proliferation is dependent on the exogenously provided growth factor SCF, and they require activity of only ER-Hoxa9 to maintain their differentiation blockade. In acute myeloid leukemia, differentiation blockades are more complex, with multiple oncogenic drivers and accessory mutations contributing to the transformed state. Thus, we next tested whether perturbing nucleotide metabolism could also promote activation of differentiation programs in human AML models with different genetic drivers and lineage characteristics. THP-1 (MLL-AF9, TP53^mut^) and U937 (CALM-AF10, TP53^mut^) AML cells can differentiate down a monocytic lineage (Daigneault et al., 2010). Purine and pyrimidine synthesis inhibitors, as well as replication stress inducers, upregulated CD11b as well as the macrophage surface markers CD14 and CD16 in these cells (**Figure 3A, S3A)**. MOLM14 (MLL-AF9, TP53^WT^), OCIAML3 (NPM1c, TP53^WT^), MV411 (MLL-AF4, TP53^WT^), and KASUMI1 (AML1-ETO, TP53^mut^) cells exhibited varying degrees of CD11b and/or CD14 induction upon treatment with BRQ or HU **(Figure S3B)**. In K562 erythroleukemia cells (BCR-ABL, TP53^mut^), which are poised to differentiate down an erythroid lineage (Sutherland et al., 1986), nucleotide depletion with BRQ and HU treatment increased surface expression of the erythroid marker CD235a (glycophorin A) (**Figure 3B**).

**Figure 3.**
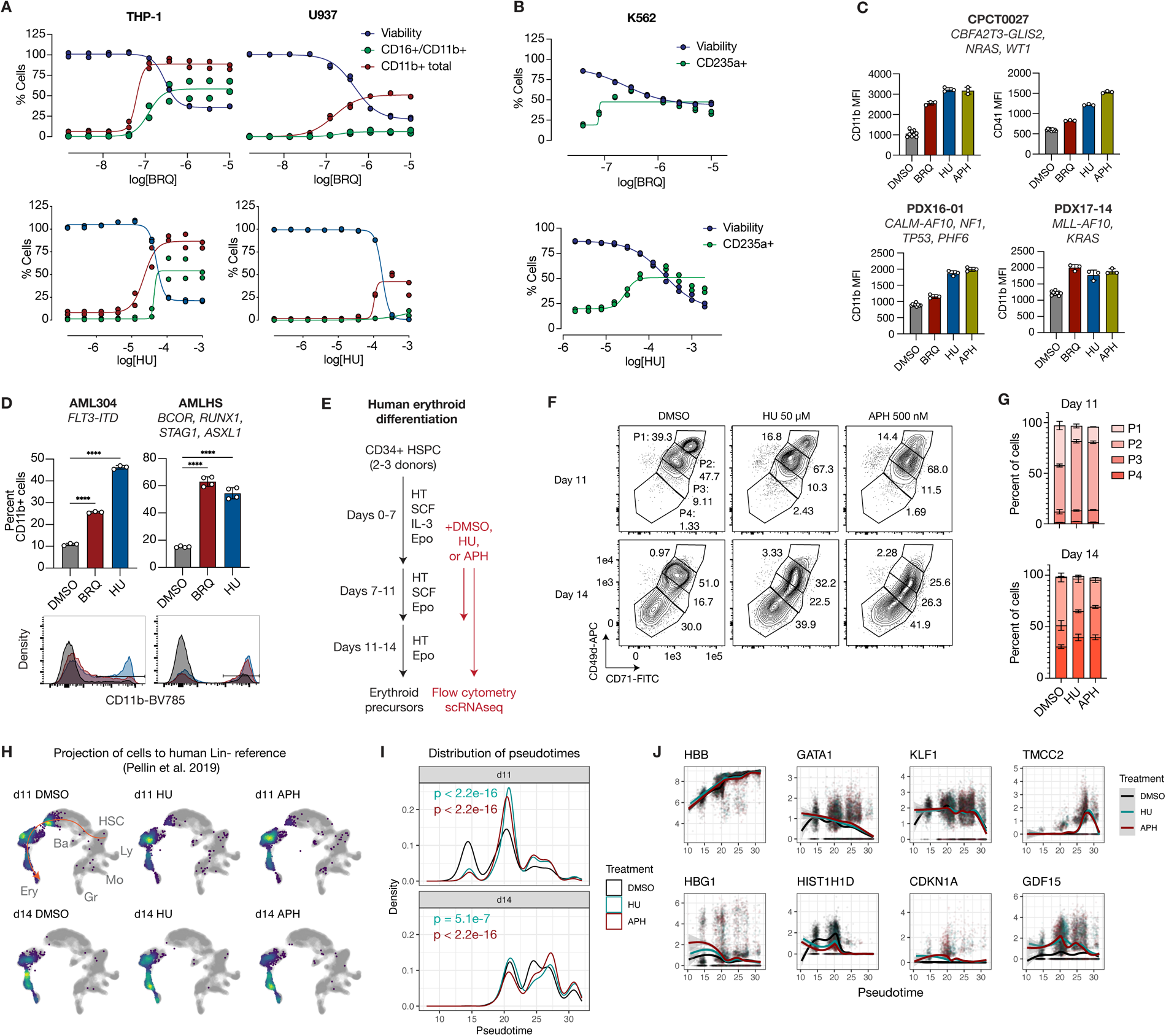
Nucleotide depletion and replication stress induce cell fate progression in AML models and normal erythroid differentiation. A. Percentage of CD11b+, CD11b+/CD16+, and viable THP-1 (left) or U937 (right) cells upon treatment with BRQ (top) or HU (bottom) for 96h (n=2). B. Percentage of CD235a+ and viable K562 cells upon treatment with BRQ (top) or HU (bottom) for 72h (n=2). C. Mean fluorescence intensity of various surface markers for three PDX AML lines cultured *in vitro* with DMSO, BRQ, HU, and APH for 72h. Concentrations are provided in Methods. D. Percentage of CD11b+ cells for two primary AML cell cultures cultured in vitro with DMSO, 1 µM BRQ, or 100 µM HU for 72h. Representative flow cytometry histograms are displayed. (****) p < 0.0001 and p-values were calculated using a one-way ANOVA with Dunnett’s multiple comparisons test. E. Schematic of primary erythroid differentiation experiment from CD34+ HSPCs. F. Representative flow cytometry plots of CD71 and CD49 signal for erythroid progenitors treated with DMSO, 50 µM HU, or 500 nM APH and analyzed at day 11 and 14. G. Percentage of cells in each gate in (F) for each treatment and timepoint (n=3). H. scRNAseq transcriptomes of cells from two donors, projected onto a reference map (in gray) of human lineage-negative bone marrow cells (Pellin et al. 2019). Cells are plotted as a density map with the densest regions in yellow. Selected hematopoietic lineages are annotated. The erythroid trajectory used for pseudotime analysis in (I) is depicted in orange in the top left plot. I. Pseudotime analysis of cells treated with DMSO, HU, or APH at day 11 and 14 using the trajectory depicted in orange in the top left panel of (H). Cells from both donors were pooled for this analysis. P-values are calculated relative to DMSO using the Mann-Whitney U test. J. Loess plots for expression of selected genes (log2 normalized counts; y-axis) versus pseudotime (x-axis) for cells treated with DMSO, HU, and APH. Cells from different donors and timepoints were pooled for this analysis.

We next measured the cellular consequences of perturbing nucleotide synthesis in human AML cell lines. Replication stress markers (pChk1 and γH2AX) were induced in THP-1 cells following BRQ and HU treatment at sublethal doses that induced substantial differentiation **(Figure S3C)**. BRQ and HU both led to depletion of dNTPs **(Figure S3D,G)**. As in ER-Hoxa9 cells, BRQ led to broad metabolic changes including depletion of pyrimidine nucleobases, ribonucleosides, and ribonucleotides and alteration of glycolytic and TCA cycle intermediate abundance, whereas HU-treated cells had more restricted metabolic changes **(Figure S3E, S3H, Table S2)**. In THP-1 cells, we also compared the metabolic effects of BRQ and HU treatment to those seen after treatment with phorbol 12-myristate 13-acetate (PMA), an inhibitor of protein kinase C signaling that orthogonally induces expression of a differentiation program **(Figure S3E-F, Table S2)**. PMA-treated cells exhibited dNTP depletion, likely because PMA treatment causes them to exit the cell cycle. Nevertheless, few metabolites were differentially altered across all three conditions; these included kynurenine, nicotinamide, and ADP-ribose, metabolites that may be associated with macrophage differentiation (Viola et al., 2019). Finally, we added single nucleosides or nucleobases and found that dNTP depletion and differentiation marker induction were observed in instances where replication stress signaling was elevated and proliferation rate and viability was inhibited **(Figure S3I-L)**.

As BRQ treatment *in vivo* has been shown to induce differentiation markers in several leukemia models including THP-1 cells (Christian et al., 2019; Sykes et al., 2016), we explored whether replication stress may also be linked to this process. We subcutaneously injected THP-1 cells into nude mice and treated them intraperitoneally with vehicle or BRQ once measurable tumors formed **(Figure S3M-P)**. After three days, we observed a decrease in tumor growth in response to BRQ. Leukemia cells from these tumors exhibited increased CD11b and CD14 surface marker expression. Concomitantly, these cells also had higher levels of phosphorylated Chk1 and γH2AX as measured by flow cytometry. Thus, BRQ treatment also induces DNA replication stress alongside CD11b and CD14 expression in THP-1 leukemia cells exposed to an *in vivo* environment.

To further extend these observations to patient-derived systems, we tested the effect of nucleotide depletion and DNA replication stress on three Cas9-competent AML PDX models that are able to proliferate in vitro for short periods of time, and that are characterized by different genetic drivers and p53 statuses (CBFA2T3-GLIS2/TP53^WT^, CALM-AF10/TP53^mut^, and MLL-AF10/TP53^WT^) (Lin et al., 2022; Pikman et al., 2021). Upregulation of cell surface markers consistent with cell state maturation were observed following treatment with replication stress inducers in all three cases, including a notable myeloid/megakaryocytic cell state shift (marked by CD11b and CD41) in the CBFA2T3-GLIS2 driven PDX (**Figure 3C**). In all three PDX models, morphological changes consistent with differentiation were also observed, including a decreased nuclear-cytoplasmic ratio, increased cell size, cytoplasmic vacuolation, and altered nuclear morphology **(Figure S3Q)**. We also observed a shift towards a more differentiated myeloid cell state in two patient AML samples that were thawed and directly cultured with BRQ or HU (**Figure 3D**). Thus, pharmacologic perturbation of nucleotide metabolism and DNA replication promotes differentiation in multiple AML models with different genetic drivers, including in primary patient-derived models.

Finally, to assess the effect of replication stress on a model of physiological hematopoietic differentiation, we differentiated human CD34+ hematopoietic stem and progenitor cells (HSPCs) from three donors into erythroid precursors with or without addition of HU or APH starting at day 7 of the differentiation protocol (**Figure 3E**). Using flow cytometry, we observed that HU or APH exposure caused CD71 and CD49d to be lost more rapidly, indicating an accelerated differentiation course (**Figure 3F-G**). To determine if this differentiation was aberrant, we performed single-cell RNAseq (scRNAseq) on vehicle or drug-treated cells from two donors at day 11 and 14 and projected them onto reference maps of human lineage-depleted bone marrow cells to estimate their maturation stage (Pellin et al., 2019) (**Figure 3H**). Pseudotime analysis demonstrated that HU and APH led to accelerated maturation along the erythroid lineage (**Figure 3I**). Erythroid genes such as beta-globin (*HBB*), the transcription factors *GATA1* and *KLF1*, and the terminal erythropoiesis regulator *TMCC2* (Ludwig et al., 2019) had expression that was appropriate for each maturation stage (**Figure 3J**). On the other hand, we observed gene expression changes associated with replication stress, including upregulation of *HBG1* (encoding the gamma-globin subunit of fetal hemoglobin, which is upregulated by hydroxyurea), *CDKN1A* (encoding p21, a target of p53), and *GDF15* (often upregulated during cell stress) as well as downregulation of histone genes (**Figure 3J**). Therefore, DNA replication stress in erythroid progenitors leads to accelerated maturation.

### Genetic perturbation of DNA replication can promote differentiation

To identify genetic modulators of cell state change, we performed a flow cytometry-based genome-wide CRISPR interference (CRISPRi) screen in ER-Hoxa9 cells expressing dCas9-KRAB-mCherry (**Figure 4A, Table S3)**. By comparing guide enrichment in the top and bottom 25% of CD11b or GFP-expressing cells, we identified 185 and 261 hits whose knockdown induced each respective marker with a false discovery rate of <10% (**Figure 4B**). Each set of hits was enriched for a variety of gene sets, with DNA replication and genome integrity-related being the only overlapping gene sets that affected both CD11b and GFP expression (**Figure 4C-D**). We next focused on the 58 overlapping hits, identifying expected genes such as *Hoxa9* and *Meis1*, *Myb*, *Pbx2*, *Kat2a*, *Kat6a*, and *Men1*, which are known regulators of the Hoxa9-driven leukemic program (**Figure 4E**). Eighteen genes were involved in DNA replication and genome integrity, including origin firing (*Cdc7*, *Dbf4*, *Mtbp*, *Ticrr*, *Mcm10*), DNA polymerase subunits (*Pold2*, *Pold3*, and *Pole*), negative regulators of replication stress (*Rad51*, *Rnaseh2c*), and negative regulators of telomerase activity (*Terf1* and *Ylpm1*). Reduced origin firing itself can cause replication stress or sensitize cells to replication stress by impairing the ability of cells to access backup origins, and increased telomerase activity can also induce replication stress (Lu and Pickett, 2022; Saxena and Zou, 2022). While most hits led to reduced proliferation and/or viability as inferred from the degree of sgRNA dropout at day 6 of the screen or by comparison to common leukemia-essential genes, the converse was not true, as most knockdowns that led to reduced viability did not also induce CD11b and GFP (**Figure 4F-G, Figure S4A)**. These data are consistent with results from the chemical screen (**Figure 1**) showing that decreased proliferation/viability is not always coupled to cell state changes.

**Figure 4.**
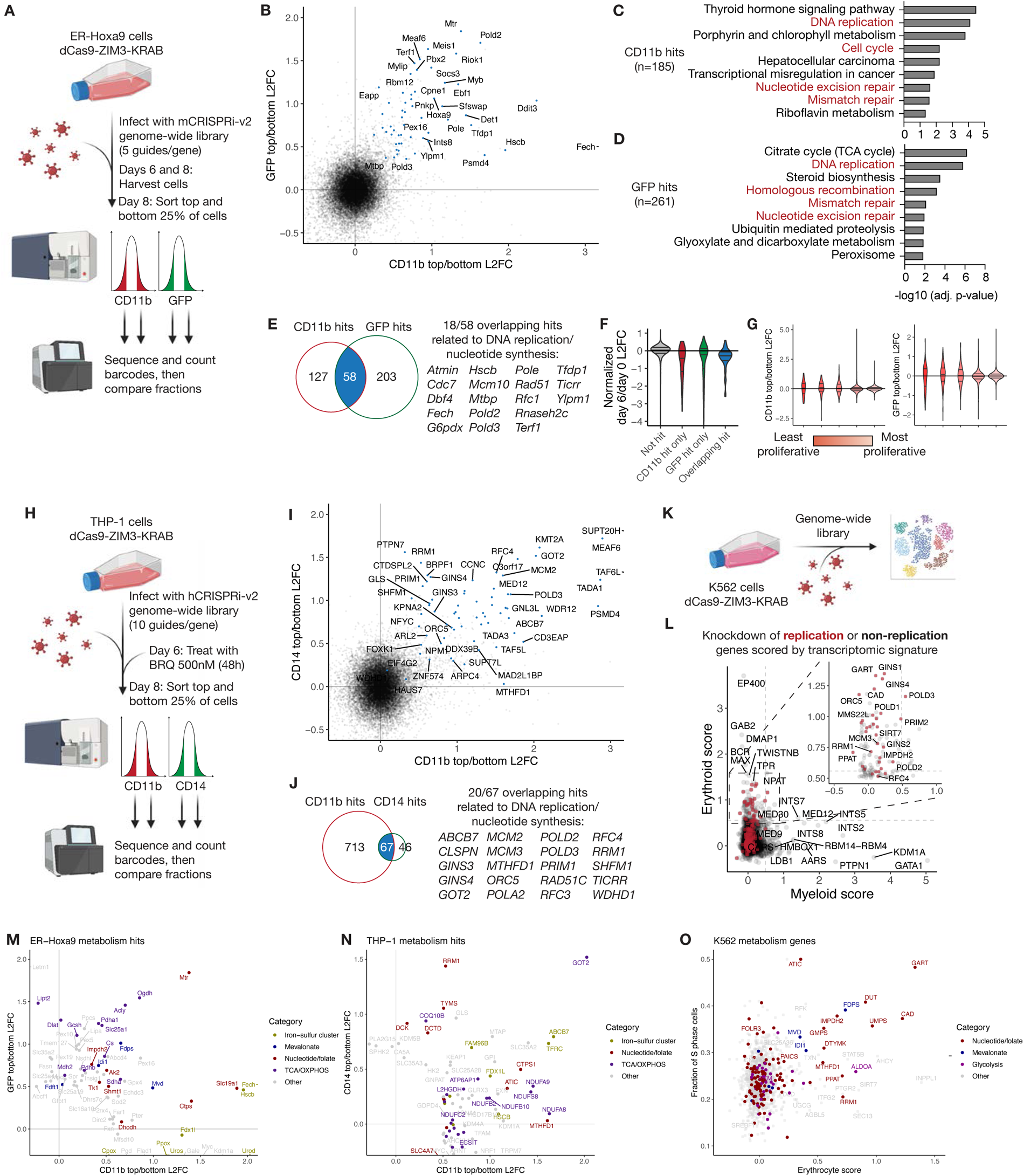
Genetic screens identify altered DNA replication as a common driver of differentiation across three leukemia models. A. Schematic of CRISPRi screen to assess modulators of CD11b and GFP induction in ER-Hoxa9 cells. B. Scatterplot of log2 fold change (L2FC) between the top and bottom 25% of the CD11b and GFP screens. Genes that are hits in both screens (FDR < 0.1 and L2FC > 0) are highlighted in blue and selected genes are labeled. C. Enriched KEGG gene ontology categories for CD11b hits, calculated using Enrichr. The negative log10 adjusted p-value is displayed. Categories related to DNA replication are highlighted in red. D. Same as (C) but for GFP hits. E. Venn diagram of the number of CD11b and GFP hits, with overlap highlighted, and overlapping genes involved in DNA replication or nucleotide synthesis listed. F. Normalized log2 fold change (L2FC) in guide enrichment between day 8 and day 6 of the screen for each class of hits and non-hits in the CD11b and GFP screens. G. CD11b (left) and GFP (right) log2 ratios for each class of genes, sorted into bins of equal width in terms of day 6/day 0 L2FC. H. Schematic of CRISPRi screen to assess modulators of CD11b and CD14 induction in BRQ-treated THP-1 cells cells. I. Same as (B) but plotting results from the CD11b and CD14 screens. J. Same as (E) but displaying overlapping hits from the CD11b and CD14 screens. K. Schematic of Perturb-seq screen performed in K562 cells by Replogle et al. (2022). L. Scatterplot of knockdowns scored by the strength of an induced myeloid or erythroid signature (see STAR Methods). Replication- and nucleotide synthesis-related genes are highlighted in red. Horizontal and vertical dotted lines represent the 99th percentile score. Inset is displayed for clarity. M. CD11b and GFP L2FC scores for metabolism hits in the ER-Hoxa9 screen in (A). All genes are displayed whose L2FC exceeds 0.5 and FDR < 0.2. Selected categories are highlighted. N. Same as (M) but for metabolism hits in THP-1 cells. O. Erythroid score in the K562 Perturb-seq screen for all metabolism genes plotted against the fraction of S phase cells. Genes with an erythrocyte score > 0.5 are labeled, and selected categories are highlighted.

To extend this screening approach to a different cell type with slower proliferation, as well as to explore modulators of differentiation in the setting of nucleotide depletion, we combined BRQ treatment with a genome-wide CRISPRi screen in human THP-1 dCas9-KRAB cells. Cells were treated for 48 hours with 500 nM BRQ, a concentration at which approximately 50% of cells induce CD11b expression (**Figure 4H**). Using flow cytometry to sort the top and bottom 25% of CD11b or CD14-expressing cells and comparing guide enrichment, we identified genes whose knockdown prevented or further sensitized cells to BRQ-mediated induction of CD11b and CD14 expression **(Table S3)**. Genes whose knockdown weakened CD11b or CD14 induction by BRQ included general transcription factors and transcriptional/chromatin machinery **(Figure S4B-C)**, which could reflect a requirement for transcriptional changes to cause marker expression. Knockdown of many more genes increased CD11b compared to CD14; the 67 overlapping genes were again enriched for DNA replication-related categories, including origin recognition and firing (MCM2, MCM3, ORC5, TICRR, GINS3, and GINS4), DNA polymerase components (PRIM1, POLA1, POLD2, and POLD3), and negative regulators of replication stress (RFC3, RFC4, CLSPN, RAD51, WDHD1, and SHFM1) (**Figure 4I-J, Figure S4D)**. Reanalysis of data from a previously reported CRISPR knockout screen in untreated THP-1 cells (Wang et al., 2021) also demonstrated that many of these DNA replication-related genes induced CD11b and CD14 expression when knocked out **(Figure S4E, Table S3)**.

To validate select hits from the CRISPRi screen and confirm that they both perturb DNA replication and promote differentiation, we performed single gene CRISPRi knockdowns of the replication-related genes POLD3, MCM2, and RFC4, the iron-sulfur cluster transporter ABCB7, and the transaminase GOT2 in THP-1 cells. Knockdown of these genes all decreased EdU incorporation in S phase cells, indicating reductions in total DNA replication activity **(Figure S4F)**. The percentage of CD11b+ cells was increased compared to cells expressing non-targeting sgRNAs (sgNTC), and BRQ treatment caused further CD11b induction **(Figure S4G)**. Gene set enrichment analysis (GSEA) revealed upregulation of genes that are induced by the differentiation agent PMA **(Figure S4H)**, including multiple markers of monocyte-to-macrophage differentiation (*MAFB*, *SLC15A3*, *CD86*, *CD163*, and *PLIN2*), as well as downregulation of monocytic genes (*CTSG* and *MS4A3*) **(Figure S4I)**.

We next assessed cell fate progression in K562 cells upon disruption of DNA replication by re-analyzing a Perturb-seq experiment in these cells where single-cell RNA-seq was performed for CRISPRi knockdown of most human genes (Replogle et al., 2022) (**Figure 4K, Table S3)**. K562 cells have characteristics of erythroid precursors but can also acquire myeloid characteristics in response to certain perturbations (Sutherland et al., 1986), affording the opportunity to assess whether disrupting DNA replication is selective for maturation along a particular lineage in these cells. Each CRISPRi knockdown was scored by degree of induction of an erythroid gene expression signature (which included genes such as *HBZ* and *SLC25A37*) or a myeloid gene expression signature (which included genes such as *SPI1* and *CSF3R)*. We found that knockdown of many genes involved in DNA replication and nucleotide synthesis, such as POLD3, GINS4, and GART, led to high erythroid scores, as did knockdown of the oncogenic driver BCR-ABL using guides targeting the *BCR* promoter (**Figure 4L**). The enrichment for genes related to nucleotide synthesis and DNA replication within the top-scoring erythroid transcriptomes stood in stark contrast to genes whose knockdown led to a high myeloid score, most of which were transcription factors such as the erythroid factor GATA-1 or epigenetic machinery such as LSD1 (encoded by *KDM1A*) and the Integrator components INTS2 and INTS5 (**Figure 4L, S4J)**. We also assessed the relationship between cell state maturation and viability. While some knockdowns causing reduced proliferation over the course of the eight-day screen also exhibited some erythroid maturation, depletion of DNA replication genes led to a much greater erythroid score for a given degree of proliferation than depletion of other classes of genes (p=2e-4) **(Figure S4K)**. Taken together, these data support that perturbing DNA replication specifically induces maturation toward an already primed fate (e.g. myeloid in ER-Hoxa9 and THP-1 or erythroid in K562).

### Involvement of metabolism in cell state changes

We next sought to understand the consequences of genetically perturbing metabolism by reanalyzing our previous screens with a focus on 3491 manually curated human genes, and their murine counterparts, spanning a wide range of metabolic pathways (Abbott et al., 2023) **(Table S3)**. Many metabolic genes are essential, and the rapid 12-hour doubling rate of ER-Hoxa9 cells led to significant dropout of sgRNAs targeting many metabolic genes over the course of the eight-day screen (165 genes with >2-fold depletion, versus only 24 genes in the THP-1 screen), reducing our ability to assess phenotypes with confidence. Thus, we considered all metabolic genes from the ER-Hoxa9 CRISPRi screen whose knockdown induced either CD11b or GFP. We identified hits from multiple categories, including citrate metabolism (*Acly*, *Cs*, *Slc25a1*), lipoic acid metabolism and its dependent enzymes (*Gcsh, Dlat, Lipt2, Pdha1, Ogdh*), nucleotides (*Mtr*, *Shmt1*, *Impdh2*, *Tk1*, *Ak2*, *Ctps*, *Dhodh*), heme and iron-sulfur cluster metabolism (e.g. *Fech*, *Urod*, *Cpox*, *Hscb*), and mevalonate synthesis *(Fdps*, *Idi1*, *Mvd*) (**Figure 4M**). While knockdown of TCA cycle hits uniquely affected GFP levels and knockdown of heme pathway hits uniquely affected CD11b levels, knockdown of nucleotide and mevalonate pathway hits affected expression of both markers. Interestingly, depletion of the mevalonate pathway genes *Fdps, Idi1, and Mvd* was previously shown to cause accumulation of a toxic pyrophosphate that may interfere with DNA replication (Horlbeck et al., 2018; Replogle et al., 2020). We followed a similar approach to identifying metabolic hits in THP-1 cells treated with BRQ and identified nucleotide metabolism and iron metabolism as pathways where CRISPRi knockdown increased both CD11b and CD14 (**Figure 4N**). In the K562 Perturb-seq screen, where knockdown of essential genes can be more confidently characterized by analyzing their single-cell transcriptomes, metabolic gene hits whose knockdown led to a high erythroid score were involved in nucleotide, mevalonate, and glycolytic metabolism (**Figure 4O**). Notably, the nucleotide and mevalonate metabolism hits, but not the glycolysis hits, had elevated percentages of cells in S phase, consistent with an effect on DNA replication (**Figure 4O**). Together, these results suggest that while each cell type (as well as programs within each cell type) may be influenced by different metabolic pathways, inhibiting nucleotide synthesis or DNA replication appear to be general drivers of cell state change. We note that many of the other pathways identified in specific cell types, including glycolysis and oxidative phosphorylation, may also feed into nucleotide biosynthesis in a manner that is dependent on cellular context and the extracellular nutrient environment (Shi et al., 2023).

To increase our sensitivity to detect metabolic hits whose knockdown impacts cell state, we built a custom CRISPRi sublibrary focused on hits that most strongly affected CD11b expression in THP-1 cells treated with BRQ, then re-screened THP-1 cells for CD11b and CD14 expression in vehicle (DMSO) or BRQ treatment **(Figure S4L)**. Results from both vehicle and BRQ screens were highly correlated **(Figure S4M)**. Focusing on hits in the vehicle screen, we found that knockdown of various genes involved in nucleotide biosynthesis induced both CD11b and CD14 expression. These included genes involved in the metabolism of pyrimidines (CAD, UMPS, TYMS, CTPS1), purines (ATIC, GART, ADSL, IMPDH2), dNTPs (RRM1, RRM2), and folates (MTHFD1) **(Figure S4N)**. We then infected K562 dCas9-KRAB cells with this sublibrary and isolated cells in the top and bottom quartile of CD235a erythroid marker expression. Of the 40 hits whose knockdown induced both CD235a expression in K562 cells and CD11b/CD14 expression in THP-1 cells, 22 were involved in DNA replication and nucleotide biosynthesis, consistent with results from the Perturb-seq data **(Figure S4O)**. Therefore, these processes appear to be convergent in their ability to influence cell identity across three different hematopoietic model systems.

### Differentiation is induced by perturbed DNA replication independently of replication stress response signaling or DNA damage

Given the effect of perturbing nucleotide synthesis and DNA replication on differentiation, we wondered whether differentiation occurs during S phase as a direct result of defects in DNA replication stress, or during other parts of the cell cycle, perhaps in response to replicative defects that persist past S phase. Using ER-Hoxa9 cells co-expressing the differentiation reporter lysozyme-GFP and the FUCCI geminin-mCherry reporter, which marks cells in S/G2/early M phase (Sakaue-Sawano et al., 2008), we found that mCherry-positive cells were the first to exhibit GFP expression following BRQ or HU treatment (at 12-14 hours) (**Figure 5A**). At later timepoints (20-24 hours), a notable percentage of GFP+ cells was evident in the mCherry-negative G0/G1 fraction, likely as a result of continued cell cycle progression and cell division. On the contrary, ER-Hoxa9 inactivation with fulvestrant allowed cells from both G0/G1 and S/G2/M fractions to differentiate equally quickly following drug exposure (**Figure 5A**). We further localized the onset of replication stress-induced differentiation to S phase, as ∼90% of CD11b+ cells 16 hours after treatment with BRQ or HU were actively incorporating EdU (**Figure 5B**). As an orthogonal method of linking cell-cycle stage to differentiation status, we performed single-cell RNA sequencing at various timepoints. Most cells 24 hours after BRQ treatment were localized in S or G2 phase while also scoring highly for a “granulocyte signature”, whereas cells expressing a granulocytic signature following E2 withdrawal were not localized to any cell cycle phase **(Figure S5A)**. We next asked whether S phase entry is required for nucleotide depletion-induced differentiation. To test this, we prevented entry of ER-Hoxa9 cells into S phase by withdrawing the growth factor SCF (**Figure 5C**). Indeed, limiting S phase entry completely blocked differentiation following BRQ treatment in contrast to ER-Hoxa9 inactivation with fulvestrant (**Figure 5D**). Together, these results demonstrate that nucleotide depletion-induced differentiation requires entry into, and initiates during, S phase.

**Figure 5.**
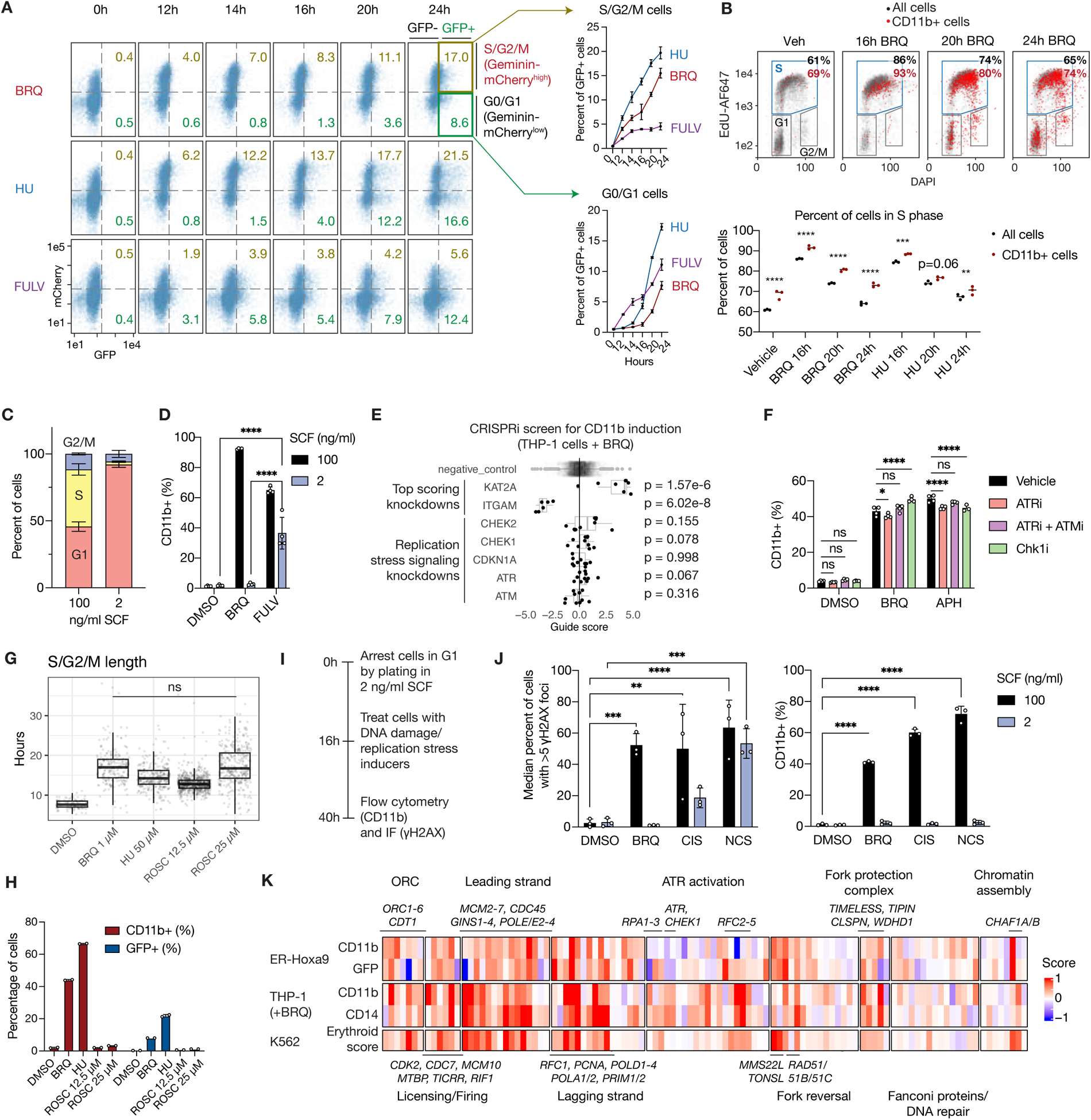
DNA replication stress drives differentiation during S phase and is independent of the replication stress response. A. (Left) Representative flow cytometry plots of ER-Hoxa9 Geminin-mCherry cells treated with 1 μM BRQ, 50 µM HU, or 10 µM FULV for different amounts of time. X-axis represents log10 GFP signal and y-axis represents log10 mCherry signal. (Right) Percentage of GFP+ mCherry^hi^ (S/G2/M) (top) or GFP+ mCherry^lo^ (G0/G1) (bottom) cells (n=4). BRQ, brequinar. HU, hydroxyurea. FULV, fulvestrant. B. (Top) Flow cytometry plots assessing EdU incorporation versus DNA content (DAPI) for ER-Hoxa9 treated with DMSO for 24 hours or BRQ for 16, 20, or 24 hours, with CD11b+ cells colored in red. The percentage of cells in S phase (the flow cytometry gate highlighted in blue) is displayed for all cells (black) or CD11b+ cells (red). (Bottom) Quantification of the percentage of all cells or CD11b+ cells in S phase for cells treated with vehicle or BRQ or HU for 16, 20, or 24 hours (n=3). (**) p < 0.01, (***) p < 0.001, (****) p < 0.0001. P-values were calculated using a two-way ANOVA with the Šidák correction for multiple comparisons. C. Percentage of ER-Hoxa9 cells in each cell cycle phase after 24h of growth in media with 100 ng/ml or 2 ng/ml SCF as determined by flow cytometry. D. Percentage of CD11b-expressing ER-Hoxa9 cells following 48h of treatment with DMSO, 1 μM BRQ, or 10 μM FULV in 100 or 2 ng/ml SCF media. P-values were calculated using a two-way ANOVA with the Šidák correction for multiple comparisons. E. Single-guide CD11b scores for negative control guides or knockdown of *ITGAM* (CD11b; negative control), *KAT2A* (a gene whose knockdown induces significant differentiation), *ATR*, *ATM*, *CDKN1A*, *CHEK1*, *and CHEK2* in THP-1 dCas9-KRAB-mCherry cells from the CRISPRi screen in Figure 3A. P-values were calculated using the Mann-Whitney U test compared to scores for negative control guides. F. Percentage of CD11b-expressing ER-Hoxa9 cells following 24h of treatment with DMSO, 1 μM BRQ, or 1 μM APH combined with either vehicle, ATRi (20 nM AZ20), ATRi + ATMi (80 nM AZD0156), or Chk1i (100 nM rabusertib) (n=3). APH, aphidicolin. P-values were calculated using a two-way ANOVA with the Šidák correction for multiple comparisons. G. S/G2/M phase length (in hours) for ER-Hoxa9 Geminin-mCherry cells treated with DMSO or the indicated drugs. The BRQ and ROSC treatments were not statistically significantly different by the Mann-Whitney U test. ROSC, roscovitine. H. Percentage of CD11b+ or GFP+ ER-Hoxa9 cells after 24h of treatment with DMSO or the indicated drugs (same concentrations as in (G)) (n=2). I. Schematic of experiments testing the effects of compounds in G1-arrested ER-Hoxa9 cells. J. (left) Median percentage of ER-Hoxa9 cells with at least 5 γH2AX foci after 24h of treatment with DMSO, 1 μM BRQ, 2.5 μM CIS, or 2.5 μg/ml NCS during asynchronous cycling or following G1 arrest in 2 ng/ml SCF. CIS, cisplatin. NCS, neocarzinostatin. (right) Percentage of CD11b-expressing ER-Hoxa9 cells after 24h of treatment with DMSO, 1 μM BRQ, 2.5 μM CIS, or 2.5 μg/ml NCS during asynchronous cycling or following G1 arrest in 2 ng/ml SCF. P-values were calculated using a two-way ANOVA with the Šidák correction for multiple comparisons. K. MAGECK gene scores for genes in each step of DNA replication and the replication stress response from the CRISPRi screens in Figure 4A (ER-Hoxa9) and Figure 4F (THP-1), as well as the erythroid score from the Perturb-seq analysis in Figure 4K (K562).

Initiation of a differentiation program during S phase prompted us to investigate the contribution of processes that occur during this phase of the cell cycle. When DNA replication is interrupted, exposed single-stranded DNA accumulates and is rapidly bound by RPA, leading to ATR activation and subsequent Chk1 phosphorylation as part of a signaling response that slows cell cycle progression and suppresses late origin firing (Zeman and Cimprich, 2014). Stressed replication forks can collapse and yield DNA breaks, which activate DNA repair machinery and signaling pathways including ATM, which phosphorylates both Chk1 and Chk2. Given the reported links between ATR/ATM signaling and induction of certain lineage-determining factors (Atashpaz et al., 2020; Sherman et al., 2010), we hypothesized that these pathways might play a role in initiating differentiation. However, results from our CRISPRi screen in THP-1 cells indicated that knockdown of any of the signaling components ATR, ATM, CHEK1, and CHEK2 did not rescue differentiation marker induction upon BRQ treatment (**Figure 5E**). To test the role of this signaling response more directly, we determined the lowest dose of ATR and/or ATM inhibitors that reduced Chk1 S345 phosphorylation to baseline in both ER-Hoxa9 and THP-1 cells **(Figure S5B-C)**. We found that these inhibitors did not suppress, and in some cases potentiated, CD11b expression caused by BRQ or APH, indicating that the replication stress signaling response may not be required for changes in differentiation-related gene expression induced by replication stress (**Figure 5F, S5D)**. Notably, while there was no effect in ER-Hoxa9 cells, ATR inhibition in vehicle-treated THP-1 cells led to low levels of CD11b and CD14 expression, demonstrating that exacerbation of endogenous replication stress by ATR inhibition is sufficient to induce differentiation in some cells (Santos et al., 2014; Toledo et al., 2013). Pharmacologic suppression of Chk1 activity also did not inhibit BRQ or APH-mediated differentiation (**Figure 5F, S5E)**.

Replication stress lengthens the cell cycle, and we tested whether this effect might be sufficient to elicit a differentiation program. To test whether differentiation is linked with the length of time spent in S phase, we used roscovitine, an inhibitor of CDK1/2 that slows origin firing without causing DNA replication stress (Petermann et al., 2010). Using live-cell imaging of previously described ER-Hoxa9 cells expressing Geminin-mCherry, we identified a dose of roscovitine (25 µM) that increased the length of S/G2/early M phase to a median of ∼17 hours, on par with the length of S phase following differentiation-inducing doses of BRQ, HU, and APH and much longer than the normal length of S phase (7-7.5 hours) (**Figure 5G**). Roscovitine slightly increased fork speed and decreased total DNA incorporation without inducing replication stress signaling, which along with a lengthened S phase suggested that it reduces the number of active origins, as reported previously (Petermann et al., 2010) **(Figure S5F-I)**. After 24 hours of roscovitine treatment, minimal CD11b or GFP expression was observed (**Figure 5H**). Thus, S phase lengthening in the absence of replication stress is insufficient to elicit a differentiation program. Next, we investigated the contribution of G1 length to differentiation at mild doses of replication stress where differentiation markers were induced. Median G1 length increased slightly (4.75-6.25 hours compared to 4.25 hours) **(Figure S5J).** However, G1 length on its own was insufficient to explain differentiation because SCF withdrawal, which induces G1 arrest, did not induce differentiation in ER-Hoxa9 cells (**Figure 5D**). A minority of cells experienced extremely long G1 phases (>12 hours), potentially corresponding to cells where under-replicated DNA (measured by quantifying 53BP1 foci) persisted into G1, but these effects were again insufficient to account for the percentage of cells that induced differentiation marker expression **(Figure S5K)**. Moreover, knockdown of p21, a CDK inhibitor that is upregulated by replication stress and has been linked to cell fate changes (Kueh et al., 2013; Santos et al., 2014), did not suppress differentiation **(Figure S5L-N)**. These data together indicate that the amount of time spent in each phase of the cell cycle is not directly coupled to differentiation state.

DNA damage has been linked to cell fate changes in several systems, leading us to explore whether the necessity and sufficiency of DNA damage for differentiation marker induction in our system (Grow et al., 2021; Inomata et al., 2009; Kato et al., 2021; Molinuevo et al., 2020; Santos et al., 2014; Wang et al., 2012). We found that the interstrand DNA crosslinker cisplatin, the DNA topoisomerase inhibitor etoposide, the DNA intercalator daunorubicin, and gamma-irradiation all induced CD11b expression in the cells tested **(Figure S5O-P)**. Exposure to cisplatin or the double-stranded break (DSB) inducer neocarzinostatin (NCS) impaired DNA replication, as evidenced by decreased EdU incorporation during S phase, and induced DNA replication stress, as marked by increased numbers of nuclear RPA foci **(Figure S5Q-R)**. To test the role of DNA damage sensing and repair independently of replication stress, we arrested cells in G1 via SCF withdrawal then exposed them to cisplatin or NCS (**Figure 5I, S5S)** (Liu et al., 2020). Markers of DNA damage, including increased number of γH2AX foci and nuclear intensity, were elevated in both G1-arrested and asynchronously cycling cells (**Figure 5J, S5T-V)**. However, differentiation did not occur in response to these agents in the G1-arrested cells, indicating that sensing and repair of intrastrand crosslinks or DSBs during G1 are not sufficient to drive differentiation in the cells tested (**Figure 5J**). While a differentiating effect of homologous recombination (HR), which mediates DSB repair in S phase or G2, cannot be ruled out, we note that BRQ induced changes towards a differentiated state prior to any evidence of a response to DSBs involving phosphorylation of Chk2 (**Figure 2F, S2L-M)**. Taken together, these data suggest that the presence of DSBs is not required for replication stress-induced differentiation.

In light of these experimental findings, we returned to our CRISPRi screens to understand how loss of the machinery that resolves replication stress influences cell state across the different systems screened. As expected, knockdown of factors involved in origin licensing, origin firing, and leading/lagging strand synthesis, which would cause replication stress, led to cell state changes across all three contexts, with some variability that could partially be attributed to dropout of essential genes (**Figure 5K**). Notably, differentiation markers were also upregulated upon disruption of machinery that buffers against the deleterious effects of replication stress – for instance, the fork protection complex (Timeless, Tipin, Claspin, and WDHD1), which modulates fork speed following redox stresses and enables ATR to activate Chk1 (Somyajit et al., 2017), and proteins that reverse and restart stalled forks, including the RAD51 recombinase and its paralogues, the MMS22L/TONSL complex, which loads RAD51 at ssDNA generated during replication stress, and the CHAF1A/B histone chaperone, whose chromatin assembly activity is important for recruitment of MMS22L/TONSL (Huang et al., 2018). Moreover, loss of ATR or components of ATR activation pathways, including RFC2-5, induced differentiation markers in some cell contexts but not others (Saldivar et al., 2017). Thus, depletion of the machinery that responds to replication stress appears to potentiate cell state changes by amplifying endogenous or exogenous sources of replication stress experienced by the cell. These results are consistent with our experimental findings that activation of cell cycle lengthening and DNA double-stranded breaks are insufficient on their own to drive differentiation and suggest a mechanism of differentiation marker induction that is independent of known cellular responses to replication stress.

### DNA replication stress induces maturation-related gene expression without fully inactivating progenitor programs

To understand how the cell fate change induced by replication stress compares to suppression of the oncogene-enforced differentiation block, we first compared bulk RNA-seq data over time from ER-Hoxa9 cells following either BRQ treatment or E2 withdrawal-induced inactivation of Hoxa9. Both treatments led to expression changes at thousands of genes, with more extensive transcriptome remodeling at later timepoints following E2 withdrawal (**Figure 6A, S6A, Table S4)**. Supplementing BRQ-treated cells with uridine reversed the transcriptional effects of BRQ (1 differentially expressed gene versus 2,299) **(Table S4)**. Within 24-48 hours, cells exposed to either BRQ or E2 withdrawal progressed to a more differentiated state characterized by expression of genes involved in neutrophil activation and regulation of inflammatory responses, such as primary granule genes (*Mpo*, *Elane*, and *Prss57*) and transcription factors (*Spi1*, *Cebpe*) (**Figure 6B, S6B)**. However, unlike cells where Hoxa9 activity was lost, BRQ-treated cells did not downregulate GMP/progenitor transcription factors and markers (*Flt3, Meis1, Cd34*) and alternative B and T cell lineage genes (*Ebf1*, *Thy1*, *Rag1*) (**Figure 6B, S6C)**. In BRQ-treated cells, we also observed consistent upregulation of normally transiently expressed genes such as *Mpo* and *Prtn3* as well as weaker (or no change in) expression of late-differentiation genes (*Lcn2*, *Camp*, *Lyn*, *Fpr1*) even by 96 hours (Lehman and Segal, 2020; Muench et al., 2020) (**Figure 6B**). Comparison with single-cell transcriptomes of normal murine neutrophil precursors demonstrated that unlike E2 withdrawal, which drove steady differentiation, BRQ-treated cells appeared to stall at an intermediate stage after 48 hours (Xie et al., 2020) (**Figure 6C**). Using cleavage under targets and release using nuclease (CUT&RUN), we mapped ER-Hoxa9 binding sites on chromatin and found that they were retained following BRQ treatment, unlike in E2 withdrawal where almost all peaks disappeared **(Figure S6D-E)**. Genes that were downregulated by E2 withdrawal but not BRQ, including the progenitor factors *Meis1* and *Ebf1*, were enriched for ER-Hoxa9 occupancy **(Figure S6F-G)**. Thus, replication stress initiates a partial granulocytic maturation program in ER-Hoxa9 cells despite continued activity of the Hoxa9-driven oncogenic program.

**Figure 6.**
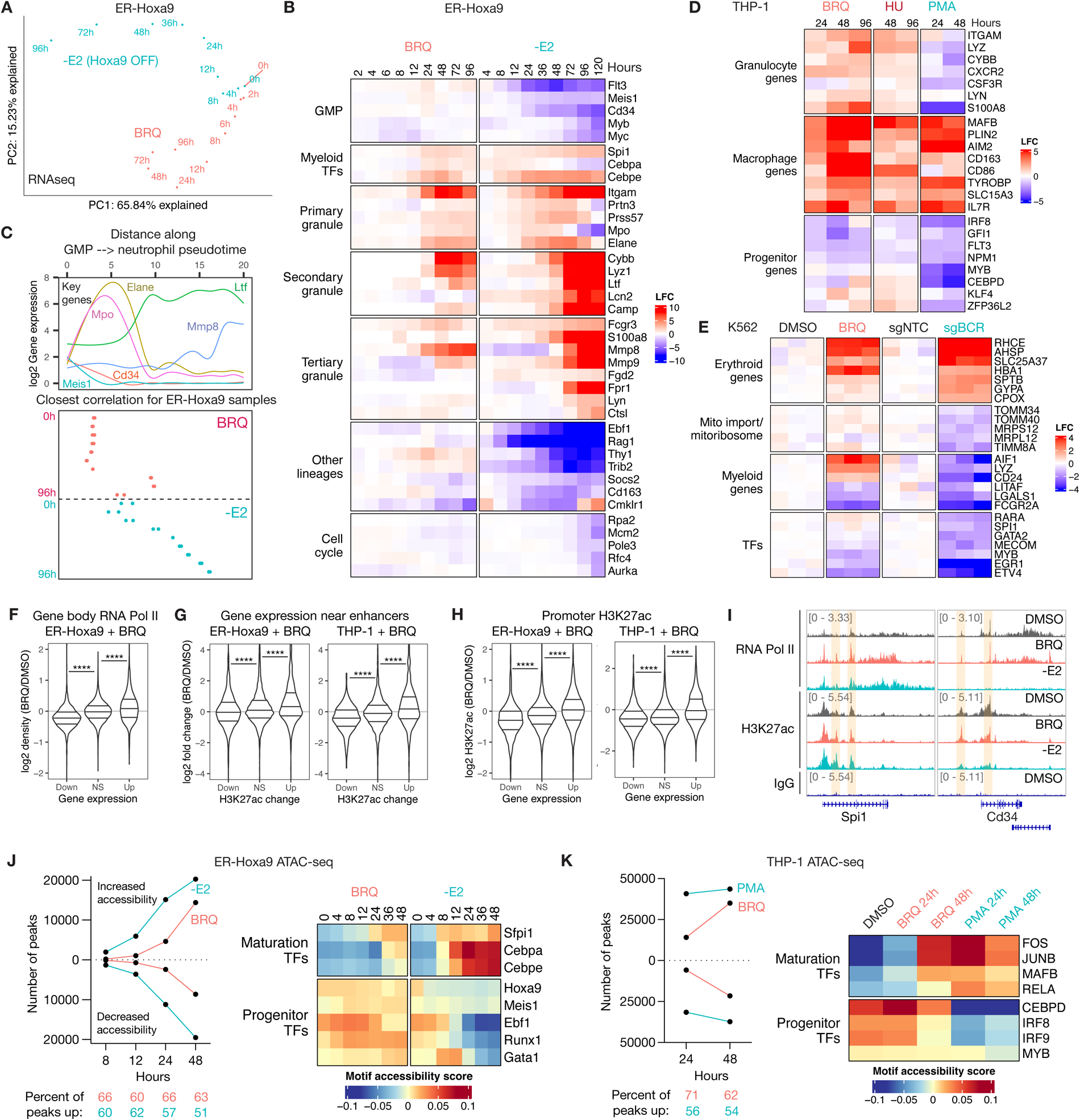
Replication stress changes cell fate at the transcriptional and epigenetic level despite maintenance of progenitor transcription factor activity. A. Principal components analysis (PCA) plot of bulk RNA-sequencing in ER-Hoxa9 cells following 2 μM BRQ treatment or E2 withdrawal for different timepoints. B. Heat maps showing expression of selected genes in ER-Hoxa9 cells within progenitor modules (GMP and other lineages), maturation modules (myeloid TFs, primary/secondary/tertiary granules), and cell cycle following BRQ treatment or E2 withdrawal for different timepoints. C. (Top) Smoothed log2 expression of selected GMP and maturation genes along a pseudotime trajectory of normal mouse GMP to neutrophil differentiation, calculated using scRNAseq data from (Xie et al., 2020). (Bottom) Projection of each timepoint from BRQ and -E2 RNA-seq in ER-Hoxa9 cells to the pseudotime trajectory. The point along the GMP-to-neutrophil trajectory to which each timepoint during BRQ and -E2 treatment is most highly correlated is plotted. D. Heatmap of gene expression in THP-1 cells treated with 500 nM BRQ, 100 μM HU, or 100 nM PMA for the indicated timepoints. Expression of selected granulocyte, macrophage, and progenitor genes is displayed. E. Heatmap of gene expression in K562 dCas9-KRAB cells transduced with a non-targeting guide (sgNTC) or a guide targeting the *BCR* promoter (sgBCR), or treated with DMSO or 250 nM BRQ for 72h. Expression of selected erythroid genes, mitochondrial import or mitoribosome genes, myeloid genes, or transcription factors is shown. F. Log2 fold change in gene body RNA polymerase II occupancy (CUT&RUN) for genes downregulated, upregulated, or not statistically significantly changed (NS) after 24 hours of BRQ treatment in ER-Hoxa9 cells. (****) p < 0.0001, Mann-Whitney U test. G. Log2 fold change in gene expression (RNAseq) for genes within 100kb of distal regulatory elements (“enhancers”) whose H3K27ac signal is downregulated, upregulated, or not statistically significantly changed (NS) after 24 hours of BRQ treatment in ER-Hoxa9 and THP-1 cells. (****) p < 0.0001, Mann-Whitney U test. H. Log2 fold change in promoter H3K27ac signal (CUT&RUN) for genes downregulated, upregulated, or not statistically significantly changed (NS) after 24 hours of BRQ treatment in ER-Hoxa9 and THP-1 cells. (****) p < 0.0001, Mann-Whitney U test. I. IGV browser screenshots for two representative genes (*Spi1* and *Cd34*) in ER-Hoxa9 cells. RNA polymerase II and H3K27ac CUT&RUN signal 24h after DMSO or BRQ treatment or E2 withdrawal are displayed, as well as IgG CUT&RUN signal. Putative regulatory regions with changes in H3K27ac signal are highlighted. J. (left) Number of differentially accessible peaks (increased accessibility plotted as positive values and decreased accessibility plotted as negative values) at various timepoints following 2 μM BRQ treatment or E2 withdrawal in ER-Hoxa9 cells. Shared differentially accessible peaks are also plotted. (right) Selected transcription factor motifs and their accessibility scores from ChromVAR analysis of ATAC-seq. K. Same as (F), but for BRQ and PMA treatment in THP-1 cells.

We next asked whether similar features governed the effect of replication stress in THP-1 and K562 cells, where endogenously enforced oncogenic drivers (MLL-AF9 and BCR-ABL, respectively) maintain a differentiation blockade. In THP-1 cells, we compared BRQ and HU treatment to the differentiation agent PMA, which is known to downregulate the oncogenic driver MLL-AF9 (Pession et al., 2003) **(Table S4)**. All three perturbations increased mRNA levels of genes associated with monocyte-to-macrophage differentiation (*MAFB*, *CD163*, *CD36*) and downregulated the expression of some stem and monocytic progenitor genes (*NPM1*, *IRF8*, and *FLT3*) (**Figure 6D, S6H)**. However, BRQ and HU treatment further activated granulocytic genes (*ITGAM*, *LYZ*, *LYN*) and did not downregulate other leukemia progenitor genes (*MYB*, *KLF4*, *ZFP36L2*), two categories of genes that were suppressed by PMA treatment (Phanstiel et al., 2017; Wang et al., 2021; Zhao et al., 2014). CRISPRi of DNA replication machinery components similarly induced macrophage and granulocyte-related genes while retaining expression of these progenitor factors **(Figure S4I)**. In K562 cells, we compared BRQ treatment to CRISPRi knockdown of BCR-ABL using guides targeting the *BCR* promoter (sgBCR), which induces an erythroid transcriptional signature and surface expression of CD235a (**Figure 4L, S6I-J)**. BRQ treatment did not suppress BCR-ABL expression **(Figure S6J)**, yet like BCR-ABL knockdown strongly induced genes involved in heme biosynthesis and erythroid differentiation (*SLC25A37*, *HBA1*, *RHCE*) and downregulated myeloid genes (*FCGR2A*, *LGALS1*, *LITAF*) and transcription factors (*MYB*, *EGR1*, *ETV4*) that mark a more immature state (**Figure 6E, S6K, Table S4)**. BRQ treatment only weakly induced other erythroid genes (*CPOX, GYPA*, *SPTB*) and led to retained or elevated expression of progenitor TFs (*GATA2*, *SPI1*, *MECOM*), myeloid genes (*AIF1*, *LYZ*, *CD24*), and mitochondrial machinery (*TOMM*/*TIMM* protein import genes and *MRPL*/*MRPS* mitochondrial ribosome genes) (Caielli et al., 2021) (**Figure 6E, S6K)**. Gene set enrichment analysis (GSEA) confirmed that genes normally downregulated during maturation of primary human erythroblasts were less robustly repressed by BRQ treatment than by sgBCR **(Figure S6L)** (Ludwig et al., 2019). Thus, replication stress also enables both THP-1 and K562 cells to express genes characteristic of a more differentiated state despite persistence of an oncogene-enforced differentiation blockades, while still expressing features of the progenitor state.

### Expression changes coincide with changes in chromatin state and occur during S phase

To understand whether changes in transcriptional activity contribute to the gene expression differences observed upon replication stress, we first performed CUT&RUN for RNA polymerase II. In ER-Hoxa9 cells, changes in gene expression were concordant with changes in gene body Pol II occupancy upon both BRQ and E2 withdrawal (**Figure 6F, S7A)**. Notably, a global shift in Pol II occupancy upon BRQ treatment was absent. We next used CUT&RUN to measure levels of the chromatin mark H3K27ac, which decorates active chromatin at both promoters and distal regulatory regions such as enhancers. In both ER-Hoxa9 and THP-1 cells, statistically significant changes in H3K27ac occurred at both promoter and distal regions, and these changes were biased towards gains in H3K27ac **(Figure S7B-C)**. Distal open chromatin regions with increased H3K27ac signal were located nearby genes that gained expression (**Figure 6G, S7D)**. Interestingly, promoters as a class slightly lost H3K27ac during BRQ treatment, but upregulated genes had less reduction of promoter H3K27ac than downregulated genes (**Figure 6H, S7E)**. Notably, genes such as *Spi1* gained RNA polymerase II and H3K27ac signal following both BRQ treatment and E2 withdrawal whereas *Cd34*, whose transcript levels were maintained in BRQ, retained RNA polymerase II and H3K27ac signal (**Figure 6I**). We also investigated whether the transcriptional and epigenetic changes observed upon replication stress initiate while cells are still in S phase. Using THP-1 cells expressing the PIP-FUCCI construct, which allows discrimination of cells from each phase of the cell cycle, we sorted early S phase cells and performed RNAseq and H3K27ac CUT&RUN after 8 hours of HU treatment, when cells were still predominantly in early to mid S phase **(Figure S7F-H)**. At this early timepoint, we observed changes in gene expression and H3K27ac that foreshadowed those seen after 24h of BRQ treatment in asynchronously cycling cells **(Figure S7I-J)**. Together, these data provide further evidence that the differentiation program induced by replication stress begins at the transcriptional and epigenetic level during S phase.

To examine how these changes are related to chromatin accessibility and to nominate transcription factor programs that might drive these changes, we performed a timecourse of ATAC-seq after BRQ treatment or E2 withdrawal in ER-Hoxa9 cells. Chromatin accessibility changes were skewed towards gains in accessibility in BRQ and enriched at introns and intergenic regions (**Figure 6J, S7K)**, consistent with enhancers playing important roles in driving cell fate progression (Blanco et al. 2021). Similar patterns were observed in THP-1 cells treated with BRQ or PMA for 24 or 48 hours (**Figure 6K, S7L)**. We next analyzed transcription factor motif accessibility using ChromVAR (Schep et al., 2017). In ER-Hoxa9 cells, motifs for the myeloid TF PU.1 (Sfpi1/Spi1) gained accessibility within 12 hours following either BRQ treatment or E2 withdrawal, whereas motif accessibility for the neutrophil-specifying Cebp family members, particularly Cebpa and Cebpe, was only weakly induced by BRQ (**Figure 6J, S7M)**. Strikingly, motifs for Hoxa9, its target Meis1, or progenitor/alternative lineage TFs within the Ebf, Runx, and Gata families all retained accessibility, suggesting that these TFs not only remain expressed during BRQ treatment, but also appear to remain active. In THP-1 cells, AP-1 and bZIP family TFs, which play key roles in differentiation of monocytes to macrophages (Phanstiel et al., 2017), gained accessibility after both BRQ and PMA treatment, but motifs for monocyte progenitor-specifying TFs such as CEBPD, MYB, and IRF family members remained open during BRQ treatment (**Figure 6J, S7N)**. In both cell lines, ATAC-seq motif accessibility patterns coincided with motifs enriched within peaks where H3K27ac was significantly changed **(Figure S7O-P)**. Peaks with increased H3K27ac upon BRQ treatment were also enriched for some motifs associated with the progenitor state (MYB in ER-Hoxa9 and PBX/HOXA in THP-1), suggesting that some regulatory loci with these motifs may gain activity without further increases in accessibility. Thus, DNA replication stress appears to drive increased accessibility and enhancer activity at the targets of expressed lineage-specifying TFs even as the motif accessibility of many progenitor TFs is retained.

### Activation of pre-existing accessible regulatory regions initiates a lineage-specific program

Lastly, we sought to understand how replication stress selectively activates a lineage-specific transcriptional program. We observed that genes that were highly upregulated by BRQ treatment in ER-Hoxa9 cells, including effectors such as *Slpi* and *Csf2rb*, were often surrounded by open chromatin regions that gained H3K27ac but that exhibited relatively unchanged chromatin accessibility, whereas H3K27ac and ATAC-seq seemed to change in tandem upon E2 withdrawal (or PMA treatment in THP-1 cells) (**Figure 7A, S7A)**. Among genes that were upregulated upon both treatments (“shared upregulated genes”), fewer increases in accessibility at nearby open chromatin regions (both proximal and distal to the TSS) occurred upon BRQ treatment; we observed similar effects in THP-1 cells **(Figure S8B-C)**. We then analyzed all open chromatin regions that gained H3K27ac following each treatment and found that smaller changes in chromatin accessibility occurred after BRQ treatment than after E2 withdrawal in ER-Hoxa9 cells or PMA treatment in THP-1 cells (**Figure 7B-C, S8D)**. We found that this was because regions with elevated H3K27ac during BRQ treatment were more accessible to begin with (**Figure 7D**). These data suggest that replication stress may preferentially activate regulatory regions with pre-existing open chromatin, whereas oncogene inactivation is necessary to also activate regulatory loci that lie within closed chromatin.

**Figure 7.**
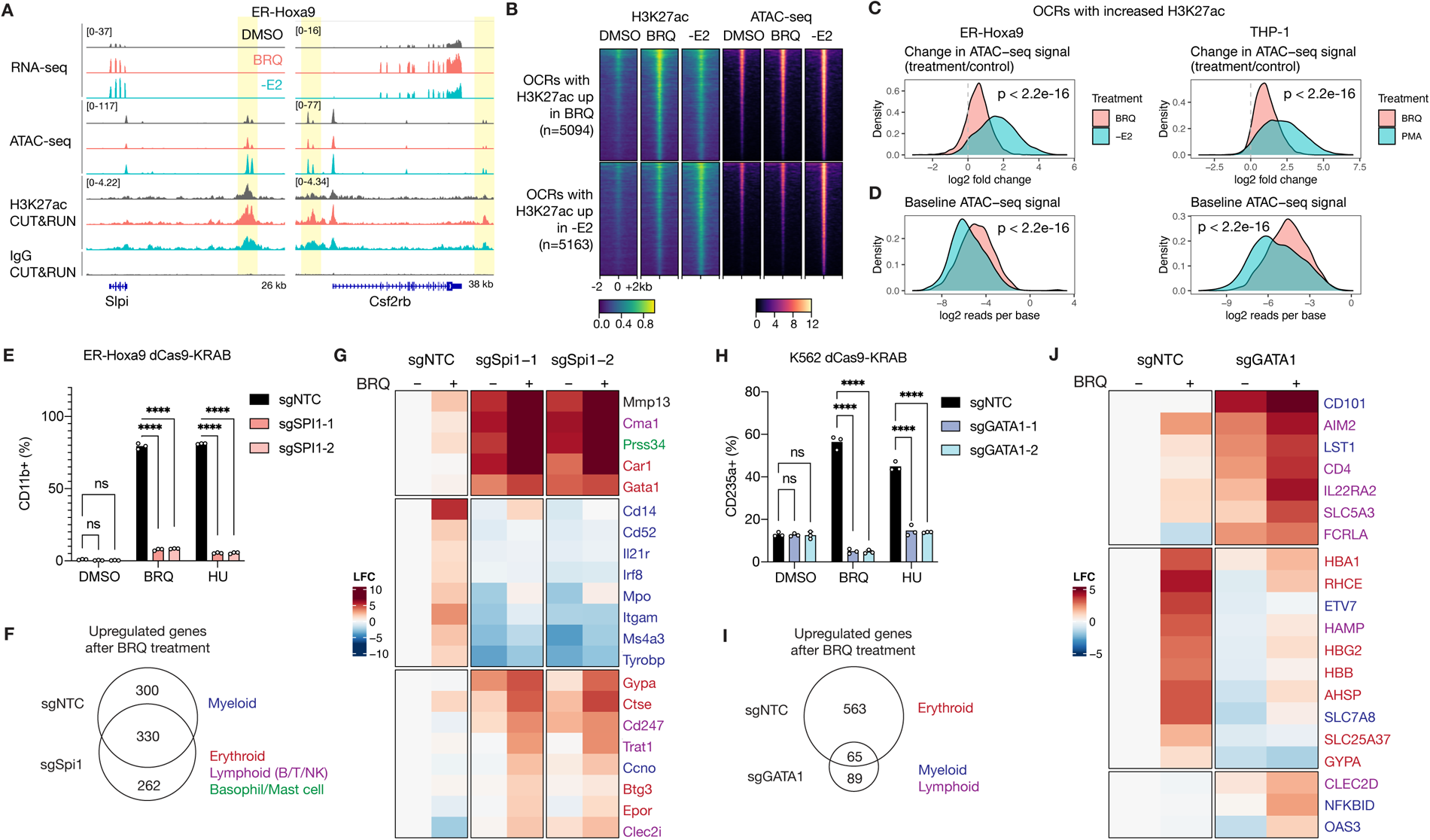
Pre-existing chromatin accessibility guides the epigenetic response to replication stress. A. IGV browser screenshots for two representative genes (*Slpi* and *Csf2rb*) in ER-Hoxa9 cells where chromatin accessibility and transcriptional changes are discordant following BRQ treatment. ATAC-seq, RNA-seq, and H3K27ac CUT&RUN signal at baseline and 24h after BRQ treatment or E2 withdrawal are displayed, as well as IgG CUT&RUN signal. B. Heatmaps of open chromatin regions (OCRs) with significantly increased H3K27ac after BRQ treatment or E2 withdrawal, displaying H3K27ac or ATAC-seq signal before and after treatment in a ±2kb window surrounding the center of the OCR. C. Log2 fold change in ATAC-seq signal (treatment/control) at OCRs with increased H3K27ac after treatment, in ER-Hoxa9 cells treated with BRQ or -E2 (left) and THP-1 cells treated with BRQ or PMA (right). P-values calculated using Mann-Whitney U test. D. Same as (C), but displaying log2 baseline ATAC-seq signal. E. Percentage of CD11b-expressing ER-Hoxa9 dCas9-KRAB cells expressing sgNTC, sgSpi1-1, or sgSpi1-2 (n=3) after 48h of treatment with DMSO, 1 μM BRQ, or 50 μM HU. (****) p < 0.0001, and adjusted p-values were calculated using a two-way ANOVA with the Šidák correction for multiple comparisons. F. Venn diagram of upregulated genes (|LFC| > 1 and adjusted p-value < 0.01) after 48 hours of 1 μM BRQ treatment of sgNTC or sgSpi1-1 ER-Hoxa9 cells, compared to DMSO. G. Heatmap showing log fold changes for selected genes in sgNTC, sgSpi1-1, or sgSpi1-2 ER-Hoxa9 cells treated with DMSO or 1 μM BRQ for 48h, relative to the sgNTC DMSO condition (n=4). H. Percentage of CD235a-expressing K562 dCas9-KRAB cells expressing sgNTC, sgGATA1-1, and sgGATA1-2 (n=3) after 72h of treatment with DMSO, 250 nM BRQ, or 125 μM HU. (****) p < 0.0001, and adjusted p-values were calculated using a two-way ANOVA with the Šidák correction for multiple comparisons. I. Venn diagram of upregulated genes (|LFC| > 1 and adjusted p-value < 0.01) after 72h of 250 nM BRQ treatment of sgNTC or sgGATA1-1 K562 dCas9 cells, compared to DMSO. J. Heatmap showing log fold changes for selected genes in sgNTC or sgGATA1-1 K562 cells treated with DMSO or 250 nM BRQ for 72h, relative to the sgNTC DMSO condition (n=3).

We hypothesized that perturbing the baseline set of open chromatin regions would predictably influence the gene expression changes caused by replication stress. To test this hypothesis, we depleted the transcription factor PU.1, encoded by *Spi1*, which establishes the myeloid lineage gene expression program and is rapidly upregulated, along with its target genes, upon induction of replication stress **(Figure S8E-H)**. ATAC-seq demonstrated that control and PU.1 knockdown cells had markedly different baseline accessibility landscapes: GATA motif accessibility was higher at baseline in PU.1 knockdown cells whereas PU.1, IRF, and ETS motif accessibility were significantly reduced **(Figure S8I)**. Upon replication stress induction in PU.1 knockdown cells, CD11b surface marker expression was impaired (**Figure 7E**), and RNA sequencing showed that 300 genes were no longer upregulated, many of which were related to myeloid differentiation such as *Itgam*, *Tyrobp*, and *Ms4a3* and had reduced baseline accessibility at nearby open chromatin regions (**Figure 7E-F, S8J-K, Table S4)**. However, 262 genes were now activated, many of which were part of the erythroid (*Gypa*, *Gata1*, *Car1*, *Epor*), mast cell/basophil (*Cma1*, *Prss34*), or T cell (*Cd247* and *Trat1*) lineage programs and had increased baseline chromatin accessibility (**Figure 7F, S8J-K)**. We also tested this paradigm in erythroid-biased K562 cells, where our Perturb-seq data had indicated that knockdown of GATA-1 reprograms these cells towards a myeloid fate (**Figure 4L**). Previous work has shown that *GATA1* knockdown in K562 cells creates a chromatin landscape characterized by increased accessibility of PU.1 and related motifs (Pierce et al., 2021). We therefore asked whether replication stress would upregulate myeloid rather than erythroid genes in *GATA1* knockdown cells. Indeed, after BRQ or HU treatment, *GATA1* knockdown cells no longer increased surface expression of the erythroid differentiation marker CD235a (**Figure 7G, S8L, Table S4)**, and they failed to upregulate many erythroid genes (**Figure 7G-H, S8M)**. Instead, 89 genes were uniquely activated in *GATA1* knockdown cells, many of which were associated with the myeloid or lymphoid lineages and include known targets of PU.1 (such as *NFKBID*, *FCRLA*, *CD4*, and *AIM2*) (**Figure 7H, S8M)**. *SPI1* was upregulated in GATA1 knockdown cells compared to controls, but it was not further upregulated upon BRQ treatment, suggesting that the myeloid genes can be induced without a requirement for PU.1 to first be upregulated further. Of the 65 genes upregulated by replication stress by in both control and GATA1 knockdown backgrounds, there was an enrichment for erythroid genes, but the magnitude of upregulation was lower in the *GATA1* knockdown cells (**Figure 7H**). Taken together, these observations are consistent with the notion that DNA replication stress activates gene expression defined by primed lineage identity programs.

## DISCUSSION

Using chemical and genetic screens and integrative genomic analyses in both non-transformed and transformed models of hematopoiesis, we find that perturbations that cause DNA replication stress are common drivers of cell state progression. In contrast to suggestions from prior work, we find that pyrimidine depletion (Sykes et al., 2016; White et al., 2011) does not appear to be unique among nucleotide-related perturbations in its ability to induce cell state progression. Our genetic screens in multiple cell lines converge on nucleotide metabolism and the DNA replication machinery as general factors critical for maintaining cell identity. Further, nucleotide depletion induces lineage markers, maturation gene expression, and H3K27ac induction at differentiation-related loci during S phase. These data argue that nucleotide depletion primarily induces differentiation through replication stress rather than disruption of other cellular processes that also require nucleotide homeostasis, such as RNA synthesis and protein glycosylation (Shi et al., 2023; Tan et al., 2016). Indeed, inhibitors of nucleotide and DNA synthesis have long been observed to induce expression of lineage-specific markers in hematopoietic cells (Ebert, 1976; Griffin et al., 1982; Gusella and Housman, 1976; Huang et al., 2004; Park et al., 2001; Santos et al., 2014). However, it was previously unclear whether this was due to downregulation of progenitor/oncogenic programs or upregulation of more mature lineage programs. We now find that replication stress can activate lineage maturation programs at the epigenetic and transcriptional level without directly suppressing the progenitor program. In non-transformed cells that lack an obligate differentiation blockade, we speculate that activation of lineage programs by replication stress can indirectly lead to downregulation of progenitor states, consistent with the accelerated, but largely normal, differentiation we and others have observed in erythroid cells treated with agents that cause replication stress (Nathan, 1985; Rencricca et al., 1975).

One possibility suggested by our work is that replication stress may selectively activate lineage-specific programs by impairing the ability of hematopoietic progenitors and oncogenic programs to actively repress primed enhancers that specify a maturation program. Although the molecular machinery that mediates this repression may be cell context-dependent, corepressors such as LSD1/GFI1, NPM1c, and FOXC1 are known to inhibit the binding or activity of multiple lineage-specific TFs at target sites (Brunetti et al., 2018; Singh et al., 2022; Stengel et al., 2021; Vinyard et al., 2019; Wang et al., 2021), and negative regulators of coactivators such as the Mediator kinase module directly repress lineage-specific enhancers in multiple leukemic settings to suppress differentiation (Dimitrova et al., 2022; Freitas et al., 2022; Pelish et al., 2015). Notably, similar to the consequences of replication stress, inhibition of LSD1 can promote differentiation without reducing chromatin accessibility at leukemic genes, indicating that the differentiation and progenitor states are not always mutually exclusive in cancer cells (Cusan et al., 2018). We note that other non-epigenetic factors, including RNA-binding proteins, may be harnessed by oncogenes to post-transcriptionally alter levels of important myeloid maturation genes; whether the activity of these factors could also be modulated by replication stress is an important question for future investigation (Wang et al., 2021). Nevertheless, the fact that replication stress can induce cell state changes in pre-leukemic and leukemic models driven by a diverse array of genetic lesions suggests that active maintenance of differentiation blocks relies at least in part on fidelity of DNA replication.

Replication stress can locally alter the fidelity of histone recycling, including for parental histones carrying repressive marks, or promote deposition of alternative histone variants (Escobar et al., 2019; Jasencakova et al., 2010; Kim et al., 2018; Wenger et al., 2023), which could ultimately lead to epigenetic and gene expression changes (Papadopoulou et al., 2015). Intriguingly, loss of the CAF-1 histone chaperone, which primarily deposits newly made histones, facilitates cell fate changes in multiple systems, yet gene expression changes often do not correlate with sites that gain chromatin accessibility, reminiscent of our observations of a discordance between chromatin accessibility and H3K27ac/gene expression changes (Cheloufi et al., 2015; Cheng et al., 2019; Franklin et al., 2022; Ishiuchi et al., 2015; Volk et al., 2018). In addition, stalled replication forks can directly recruit epigenetic machinery, including corepressors (Callen et al., 2012; Nakamura et al., 2021; Ray Chaudhuri et al., 2016; Rondinelli et al., 2017); the repressive histone mark H3K9me3 was recently observed to accumulate at stalled replication forks (Gaggioli et al., 2023). How these processes might locally or globally alter epigenetic state to drive cell state changes downstream of replication stress is another important area for future investigation.

Cell fate changes have often been documented to occur during G1 phase (Buttitta and Edgar, 2007; Pauklin and Vallier, 2013). However, other stages of the cell cycle involve dismantling and reconfiguration of chromatin state, as is seen during mitosis and G1 reentry (Hsiung et al., 2016; Zhang et al., 2021); these large changes have been postulated to provide cells with an opportunity to more easily undergo cell fate transitions. Similarly, there has been an increasing appreciation of S phase as a vulnerable period during which barriers to cell identity, such as inaccessible chromatin, repressive histone marks, or TF binding, might be more easily lowered by replication stress or alterations in replication fork speed (Ramachandran et al., 2017; Raz et al., 2021; Šviković and Sale, 2017). Recent studies across a diverse set of non-transformed systems including iPSC-derived cardiomyocytes, pancreatic endocrine differentiation, and *C. elegans* cell extrusion have also suggested that replication stress and slowed replication fork speed can facilitate or accelerate cell fate progression (Dwivedi et al., 2021; Nakano et al., 2017; Sui et al., 2021). Notably, replication stress and slowed replication forks appear to shift mouse embryonic stem cells towards a less differentiated 2-cell-like cell (2CLC) fate (Atashpaz et al., 2020; Grow et al., 2021) and can facilitate reprogramming (Nakatani et al., 2022), emphasizing that perturbing DNA replication may not always lead cells to a more differentiated state, but rather may cause cells to have increased plasticity to transit towards various primed cell fates.

The recognition that metabolic perturbations that affect DNA replication can also promote cell state changes may have implications for understanding and developing cancer therapies. These findings inform current efforts to revisit targeting of nucleotide biosynthesis and the replication machinery as differentiation therapy (de Thé, 2018), as unlike tailored therapies such as retinoic acid in acute promyelocytic leukemia, replication stress inducers appear to be more agnostic to the specific oncogenic driver or cell state. This suggests that induction of replication stress to promote differentiation may be useful in settings that are less amenable to targeted approaches; however, since replication stress is unable to suppress all oncogene-enforced progenitor programs, it may also allow cells to access states that are not therapeutically beneficial or potentially harmful. A more comprehensive understanding of how nucleotide levels and replication stress modulate transcriptional and epigenetic programs in different contexts is needed to develop strategies that alter cell state for therapeutic benefit, including in settings outside of leukemia and hematopoiesis.

## METHODS

### Cell culture

Mouse Lys-GFP-ER-Hoxa9 cells were derived as described in Sykes et al. 2016. ER-Hoxa9 cells were grown in RPMI (Corning, 15-040-CV) with 10% dialyzed fetal bovine serum (Sigma, F0392), 4mM glutamine (Thermo Fisher; #25300164), and 1% penicillin/streptomycin (Corning, 45000-652), supplemented with 0.5 μM beta-estradiol (Sigma, E2758; made from a 10 mM stock dissolved in 100% ethanol) and stem cell factor (SCF) from conditioned media generated from Chinese hamster ovary (CHO) cells that stably secrete SCF (Sykes et al., 2016; this conditioned media was added at 1% to achieve a final concentration of ∼100 ng/ml). MV411 and MOLM14 cells (a gift from Angela Koehler), OCI-AML3 cells (a gift from Yadira Soto-Feliciano), and THP-1, U937, K562, and KASUMI1 cells (ATCC) were grown in RPMI with 10% dialyzed fetal bovine serum (Gibco), 4mM glutamine, and 1% penicillin/streptomycin. Lenti-X 293T cells (ATCC) were grown in DMEM with 10% heat inactivated fetal bovine serum (Gibco) without antibiotics. All cell lines were regularly tested for mycoplasma.

For PDX samples (CPCT0027, PDX16-01, and PDX17-14), primary patient samples were acquired following written informed consent in accordance with the Declaration of Helsinki, and PDXs were established under protocols approved by Dana-Farber Cancer Institute and Cincinnati Children’s Hospital Medical Center (CCHMC) Institutional Review Boards. Detailed information regarding PDX cytogenetics, sources and molecular characterization are provided in (Lin et al., 2022; Pikman et al., 2021). For short-term in vitro liquid propagation, PDX cells were cultured in Iscove’s modified Dulbecco’s medium (IMDM, Life Technologies, #12440061), supplemented with 20% dialyzed fetal bovine serum (FBS, Life Technologies, #26400044), 1% penicillin streptomycin (P/S, Life Technologies, #15140163) and 10 ng/mL human SCF, TPO, FLT3L, IL-3, and IL-6 (PeproTech, #300-07, #300-18, #300-19, #200-03, and #200-06). For differentiation experiments, cells were cultured in brequinar (Selleck; 1 µM CPCT0027, 2 µM PDX16-01 and 5 µM PDX17-14), hydroxyurea (Selleck; 200 µM CPCT0027, and 100 µM PDX16-01 and PDX17-14) and aphidicolin (Selleck; 1 µM CPCT0027 and PDX16-01, 200 nM PDX17-14).

For primary AML patient samples (AML304 and AMLHS), cells were obtained through DFCI cell banking protocols. Cells were maintained in RPMI-1640 (Cellgro) supplemented with 1% penicillin/streptomycin (PS), 10% FBS (Sigma-Aldrich), and human IL-3 (10 ng/mL), IL-6 (10 ng/mL), TPO (10 ng/mL), SCF (25 ng/mL), and FLT3L (25 ng/mL) (PeproTech). Cell growth and viability were confirmed through daily cell counts. Upon confirmation of proliferation, cells were cultured with hydroxyurea 100 µM (Selleck), brequinar 1 µM (Selleck), or DMSO.

For primary human CD34+ HSPCs, cells were obtained from the Fred Hutchinson Hematopoietic Cell Procurement and Resource Development repository (Seattle, WA). No human subjects were involved in this study. Each vial had greater than 90% CD34 purity and had been positively selected using the Miltenyi CliniMacs system. Donors were deidentified and were negative for standard virology screening. Cells were differentiated into mature erythroid cells using a three-phase culture protocol, with base media consisting of Iscove’s modified Dulbecco’s medium (IMDM) supplemented with 2% human blood plasma (STEMCELL Technologies, #70039), 3% human AB serum (Sigma-Aldrich, H4522), 3 U/ml heparin (Sigma-Aldrich, #H3149-25KU), and 10 µg/ml insulin (Sigma-Aldrich, #I2643). In phase 1 (day 0-7), cells were cultured at a density of 1e5-1e6/ml in base media supplemented with 200 µg/ml holo-transferrin (Santa Cruz Biotechnology, #sc-391098A), 10 ng/ml SCF (Peprotech, #300-07), 1 ng/ml IL-3 (Peprotech, #200-03), and 3U/ml Epo (GenScript, #Z02975-50). In phase 2 (days 7-11), cells were cultured at a density of 1e5-1e6/ml in base media supplemented with 200 µg/ml holo-transferrin, 10 ng/ml SCF, and 3U/ml Epo. In phase 3 (days 11-15), cells were cultured at a density of 1e6-5e6/ml in base media supplemented with 500 µg/ml holo-transferrin and 0.1U/ml Epo.

### Drugs

The following drugs were used: brequinar (Sigma-Aldrich; SML0113), teriflunomide (Sigma-Aldrich; SML0936), 6-azauridine (Sigma-Aldrich, A1882), pyrazofurin (Sigma-Aldrich; SML1502), 5-fluorouracil (Sigma-Aldrich; F6627), pyrimethamine (Cayman Chemical; #16472), cladribine (Cayman Chemical; #12085), lometrexol (Cayman Chemical; #18049), 6-mercaptopurine (Sigma-Aldrich; #852678), 6-thioguanine (Cayman Chemical; #15774), mycophenolate mofetil (Sigma-Aldrich; SML0284), fulvestrant (Selleck Chemical; S1191), RX-3117 (MedChemExpress; HY-15228), hydroxyurea (Sigma-Aldrich; H8627), aphidicolin (Sigma-Aldrich; A0781), 3-AP (ApexBio; C4564), gemcitabine (MedChemExpress; HY-17026), clofarabine (Sigma-Aldrich; C7495), phorbol 12-myristate 13-acetate (Sigma-Aldrich; P8139), etoposide (Sigma-Aldrich; E1383), cisplatin (Cayman Chemical; #13119), daunorubicin (Sigma-Aldrich; #30450), cytarabine (Selleck Chemical; S1648), neocarzinostatin (Sigma-Aldrich; N9162), VE-822 (Cayman Chemical; #24198), AZ20 (Selleck Chemical; S7050), AZD0156 (Selleck Chemical, S8375), rabusertib (Selleck Chemical; S2626), palbociclib (Selleck Chemical; S1116), roscovitine (Selleck Chemical; S1153), imatinib (Tocris; #5906), fulvestrant (Cayman Chemical; #11011269), and azaserine (Cayman Chemical; #14834). All drugs were stored at a stock concentration of 10 mM in DMSO at -20°C except for hydroxyurea, which was stored at a stock concentration of 500 mM in water at -20°C, and cisplatin, which was prepared fresh from powder prior to each use and diluted into DMF. The following nucleosides and nucleobases were used: adenine (Sigma-Aldrich; A8626), cytidine (Sigma-Aldrich; C4654), guanosine (Sigma-Aldrich; G6752), uridine (Sigma-Aldrich; U3003), thymidine (Sigma-Aldrich; #89270), hypoxanthine (Sigma-Aldrich; H9636), deoxyadenosine (Sigma-Aldrich; D8668), deoxycytidine (Sigma-Aldrich, #D3897), and deoxyguanosine (Sigma-Aldrich, D7145). Adenine was prepared at 50 mM in 0.1 M NaOH. Guanosine was prepared at 300 mM in DMSO and deoxyguanosine was prepared at 30 mM in DMSO. All other nucleobases and nucleosides were prepared at 100 mM in water, and 10 mM final concentrations of uridine and cytidine were prepared directly in cell culture media. All nucleoside and nucleobase stocks were stored at -20°C.

### Chemical screen

The Ludwig metabolic library has been described previously (Harris et al. 2019) and was screened in a 10-point 3X dilution series with the highest concentration screened at 10 μM at the ICCB Longwood Screening Facility at Harvard Medical School. The Lys-GFP-ER-Hoxa9 cells were diluted to a concentration of 75,000 cells per ml, and 200 μl was dispensed into 96-well round-bottom tissue-culture treated plates. 200 nl of each compound were transferred from the 384-well compound plates to the 96-well cell plates by robotic pinning. Brequinar at a final concentration of 2 μM was used as a positive control on every 96 well plate, and DMSO was used as a negative control. Two technical screening replicates were performed. After 4 days, cells were incubated with anti-CD11b-APC antibody (final concentration of 1:400; Clone M1/70, Biolegend), washed, and resuspended in FACS buffer with DAPI dye (Thermo). Cells were analyzed in 96-well format using an iQue3 (IntelliCyt).

Viability was determined using forward and side scatter profiles. Viability for each well was normalized to the average of the 72 DMSO-treated negative controls across the four 96 well plates corresponding to one 384-well compound source plate to determine a normalized percentage viability. Differentiation was determined by measuring APC and GFP fluorescence on DAPI-negative live cells. The percent differentiation was the percentage of live cells that fell in the APC positive, GFP positive flow quadrant. Initial data processing was performed using the ForeCyt 8 software (Sartorius). The AUC of viability and differentiation were calculated with in-house R scripts. Data were graphed using Prism GraphPad and least squares curve fit (log(inhibitor) vs response; variable slope four parameters).

### Flow cytometry

Flow cytometry analysis was performed on BD FACSCanto, LSR II, and Celesta machines. For surface staining, the following antibodies were used at a 1:400 dilution: CD11b-APC (Biolegend; clone M1/70; #101211), CD11b-BV785 (Biolegend; clone ICRF44; #301346), CD14-FITC (BD; clone M5E2; #557153), CD16-FITC (BD; clone 3G8; #555406), CD235a-APC (Biolegend; clone HI264; #349114), CD41A-V450 antibody (BD Biosciences, #561425), CD71-FITC (Biolegend; clone A015; #334103), and CD49d-APC (Biolegend; clone 9F10; #304307). Cells were washed once with PBS, then stained for 15 minutes on ice followed by a single wash with PBS and resuspension in PBS with 1 ng/ml DAPI for live/dead cell discrimination.

For intracellular flow cytometry, the following antibodies were used at a 1:100 dilution: p-S345 Chk1 (Cell Signaling Technologies; clone 133D3; #2348) and p-S139 H2AX (Cell Signaling Technologies; clone 20D3; #9718). 100,000 cells were washed once with PBS, fixed for 10 minutes at room temperature with 20 μl 4% paraformaldehyde, then washed again and incubated at 4°C for 30 minutes with primary antibody diluted in 1X Click-iT saponin-based permeabilization and wash reagent (Thermo Fisher, #C10419). Cells were then washed once with PBS with 1% BSA, then incubated at 4°C for 30 minutes with goat anti-rabbit AF488 secondary antibody (Thermo Fisher; #A-11008) diluted at 1:1000 in the same saponin reagent, washed, and analyzed by flow cytometry.

For EdU incorporation experiments, the Click-iT Plus EdU Alexa Fluor 647 Flow Cytometry Assay Kit protocol (Thermo Fisher; #C10634) was used with minor modifications to the manufacturer’s instructions. Briefly, 100,000 cells were pulsed with 20 μM EdU for 30 minutes prior to cell collection. After fixation, permeabilization was performed for 15 minutes with 25 μl of 1X permeabilization and wash reagent. A master mix was made with the following amounts of reagent per sample (added in order): 19.79 μl PBS, 0.83 μl copper protectant, 0.21 μl AF647 picolyl azide, and 4.16 μl reaction buffer. 25 μl of this master mix was then added to each sample of permeabilized cells, and the click reaction was allowed to proceed at room temperature for 30 minutes in the dark, washed once with PBS, and incubated with DAPI (0.1 μg/ml) or propidium iodide (CST #4087) at 4°C for 30 minutes prior to flow cytometry analysis.

For early S phase cell isolation experiments, 40 million THP-1 cells expressing the PIP-FUCCI system (Grant et al., 2018) were concentrated to a final concentration of 2 million/ml. DyeCycle Violet (Invitrogen, #V35003) was added to a final concentration of 5 µM, and the cells were mixed well and returned to the incubator for 1 hour. At the 40-minute timepoint, ¼ of the cells was spiked with 20 µM EdU and returned to the incubator. At the 1-hour timepoint, cells were concentrated to 10 million/ml, the pellet was resuspended in the appropriate amount of supernatant containing DyeCycle Violet, and 500 µl of cells was aliquoted into flow cytometry tubes and placed on ice and sorted. S phase cells were identified as those that were Cdt1-mVenus^low^, and S phase cells in the bottom 1/3 of DyeCycle Violet signal (a proxy for DNA content) were considered to be in early S phase. Cells were sorted in a chamber cooled to 4C and kept on ice prior to HU treatment or immediate RNAseq/CUT&RUN processing in order to minimize further cell cycle progression. To validate our sorting approach, EdU-treated cells were sorted in an identical manner and fixed immediately after sorting, then subjected to processing with the Click-iT Plus EdU Alexa Fluor 647 Flow Cytometry Assay Kit and stained with DAPI. Similarly, a small aliquot of early S phase cells treated with HU for 8 hours was also spiked with EdU for 20 minutes, fixed, and processed for EdU/DAPI flow cytometry analysis.

Sorting was performed on BD FACSAria machines at the Koch Institute flow cytometry core, using the 70 μm nozzle. All cells were sorted into fresh cell culture media and kept on ice unless otherwise indicated. Gating and downstream analysis were performed using FACSDiva (BD) software, FlowJo (v10.8.0), or in-house scripts based on the FlowCytometryTools package in Python.

### Cytospins

For histological analysis, 150 µl of saturated cell culture (2 million cells/ml) was mounted on a frosted microscope slide (Fisher Scientific #1255015) via cytocentrifugation on a Shandon Cytospin 3 Cytocentrifuge (Thermo Fisher Scientific) at 500g for 5 minutes. Cells were then stained with May-Grünwald-Giemsa solution as per manufacturer’s instructions (Sigma Aldrich #MG500-500ML and GS500-500ML), prior to light microscopy and imaging on an Olympus BX41.

### Immunofluorescence

For immunofluorescence of chromatin-bound RPA and pRPA (Figure S2), 1 million cells were pre-extracted on ice with ice-cold 30 µl CSK buffer (25 mM HEPES pH 7.4, 50 mM NaCl, 1 mM EDTA, 3 mM MgCl2, 300 mM sucrose, 0.2% Triton X-100, 1 Roche cOmplete protease inhibitor cocktail tablet per 50 ml of buffer) for 1 minute, then immediately washed with 800 µl PBS with 1% BSA and pelleted at 500g for 3 minutes. The supernatant was carefully removed and cells were fixed for 10 minutes at room temperature with 50 µl of 4% paraformaldehyde freshly prepared from a 16% stock (Electron Microscopy Sciences). Cells were again washed with 800 µl PBS with 1% BSA and pelleted, then resuspended in 60 µl PBS with 1% BSA and 5 µl of cells was added to each well of a multi-well microscope slide (Electron Microscopy Sciences) pre-coated with 0.1% poly-L-lysine (Sigma Aldrich). Cells were allowed to adhere for 10 minutes, and slides were centrifuged for 5 minutes in a swinging bucket rotor at 500g for 3 minutes with acceleration and deceleration parameters set to their lowest value. After blocking with 10% normal goat serum (Sigma) for 30 minutes, samples were incubated overnight with the following primary antibodies diluted in PBS with 0.1% Tween and 1% BSA: rabbit RPA2 pS4/8 (Bethyl, A300-245A, 1:200) and rabbit RPA2 (Abcam; ab76420; 1:50). After 3 washes with PBS, samples were incubated for 1 hour with the following secondary antibodies, all at 1:1000 in PBS with 0.1% Tween and 1 ng/ml DAPI: goat anti-rabbit AF647 (Invitrogen; A-21245), then washed 3 times with PBS and mounted using ProLong Gold (Invitrogen). For quantitative image-based cytometry, images were acquired on a TissueFAXS SL microscope with a 40x/0.95NA air objective, and cell segmentation and intensity calculation was performed using CellProfiler. For acquisition of representative images, a Leica SP8 spectral confocal microscope was used with a 100X oil objective, and 30-50 z-stacks were obtained and processed using maximum intensity projection.

For RPA, γH2AX, and 53BP1 foci measurement (Figure S5), 10,000 cells were allowed to adhere in each well of multi-well slides, fixed with freshly made 4% paraformaldehyde for 10 minutes, washed 3 times with PBS, permeabilized with 0.2% triton-X in PBS for 20 minutes, washed 3 times with PBS, and blocked as above for 30 minutes. For experiments where cells had been labeled with 20 µM EdU prior to fixation, 5 µl of Alexa Fluor 488 click reaction cocktail (Click-iT Plus EdU Flow Cytometry Kit; Sigma C10633) was added for 30 minutes in the dark at room temperature, after which cells were washed three times with PBS. Samples were then incubated overnight with the following primary antibodies diluted in PBS with 0.1% Tween and 1% BSA: rabbit anti-mouse RPA2 (Abcam; ab76420; 1:50), rat anti-mouse RPA2 (CST; 2208S; 1:200), and rabbit anti-mouse γH2AX (CST; 9718; 1:1000). After 3 washes with PBS, samples were incubated for 1 hour with the following secondary antibodies, all at 1:1000 in PBS with 0.1% Tween and 1 ng/ml DAPI: goat anti-rabbit AF647 (Invitrogen; A-21245) and goat anti-rat AF647 (Invitrogen; A-21247), then washed 3 times with PBS and mounted using Fluoromount. For RPA and γH2AX foci, images were acquired on a DeltaVision Elite microscope platform (GE Healthcare Bio-Sciences) using the 60X Plan APO 1.42NA objective and a CoolSNAP HQ2 camera with 15 z-slices. Images were deconvoluted using constrained iterative deconvolution and stacks were collapsed using maximum intensity projection. Cells were identified and segmented using a semi-supervised approach, and foci and intensities were automatically tabulated using ImageJ. RPA coefficient of variation was calculated for each cell by dividing the standard deviation of RPA signal across each DAPI-positive region by its mean. For 53BP1 foci, images were acquired and analyzed with the TissueFAXS system as described above, and G1 cells were identified as those with low DAPI and low EdU signal when plotting both parameters.

### DNA fiber assay

DNA fibers were prepared as described previously with modifications (Lunt et al., 2015). Briefly, cells at 150,000 cells/ml were treated with the indicated drug for 7 hours and 20 minutes. At that point, chlorodeoxyuridine (CldU; Sigma-Aldrich) was added at a final concentration of 25 µM and allowed to incubate for exactly 20 minutes, after which iododeoxyuridine (IdU; Sigma-Aldrich) was added at a final concentration of 250 µM for exactly 20 minutes. Cells were then quickly transferred into ice-cold Eppendorf tubes, spun down in a cold centrifuge, and resuspended at 400,000 cells/ml in ice-cold PBS. Labeled cells were mixed 1:5 with unlabeled cells treated similarly, and 2.5 µl was spotted onto a glass slide and allowed to air-dry for 5 minutes, after which 7.5 µl of spreading buffer was added on top of the cell droplet. After 5 minutes, the slide was tilted and the droplet was allowed to move slowly down the slide. Slides were air-dried in the dark for 2-4 hours, after which they were fixed for 3 minutes with ice-cold 3:1 methanol:acetic acid, air-dried, and stored at 4°C overnight. Slides were then denatured with 2.5M HCl for 30 minutes, washed 4 times with PBS, and blocked with 5% BSA in PBS for 30 minutes at 37°C in a humidified chamber. For primary antibody staining, 1:50 rat anti-BrdU antibody (Abcam ab6326; detects CldU) and 1:20 mouse anti-BrdU antibody (BD 347580; detects IdU) were combined in 1% BSA in PBS + 0.1% tween (PBST) and added to slides; a coverslip was placed on top and the slides were incubated for 1 hour at 37°C in a humidified chamber. Three washes in PBST were performed, after which secondary antibody detection was performed with 1:50 of goat anti-rat AF488 (Invitrogen A11006) and 1:50 of goat anti-mouse AF546 (Invitrogen A11030) in PBST, with 30 minutes of incubation at 37°C in a humidified chamber. Three washes in PBST were performed, and slides were mounted using 20 µl of ProLong Gold Antifade Mountant (Thermo Fisher). Images were obtained using either 60X or 100X oil objectives with a Nikon 90i widefield fluorescence microscope. CldU and IdU tracts were measured manually using ImageJ, and at least 100 tracts were measured for each condition.

### Comet assay

Alkaline comet assays were performed exactly as described in (Su et al., 2021) using the comet assay kit (Trevigen), with the exception that cells were treated with the indicated drug for 12 hours, and electrophoresis was performed for 18 minutes. At least 150 cells were analyzed for each condition.

### Mouse experiments

All experiments conducted in this study were approved by the Massachusetts General Hospital Institutional Animal Care and Use Committee (IACUC). For subcutaneous tumor growth, a maximum tumor burden of 1 cm^3^ was permitted per IACUC protocol, and these limits were not exceeded. Female NU/J nude mice between 2 and 3 months old were used in this study. All animals were housed at ambient temperature and humidity (18–23°C, 40–60% humidity) with a 12h light and 12h dark cycle and co-housed with littermates with *ad libitum* access to water, unless otherwise stated. All experimental groups were age-matched, numbered and assigned based on treatment, and experiments were conducted in a blinded manner. Data were collected from distinct animals, where *n* represents biologically independent samples. Statistical methods were not used to predetermine sample size.

On the day of the experiment, 100 million THP-1 cells expressing pU6-sgNTC EF1Alpha-puro-T2A-BFP (to enable downstream analysis of tumor cells by flow cytometry) were washed and resuspended in 1 ml PBS, 1 ml Matrigel was added, and the cells were passed multiple times through an 18-gauge syringe to ensure a single cell suspension. 5 million cells (100 μl) were then injected subcutaneously into the right flank of each mouse. 13 days after cell injection, when palpable tumors had formed, animals were randomized and injected IP with either PEG or 50 mg/kg BRQ. Mice were weighed before the start of treatment to ensure that different cohorts had similar starting body weights, and body weights were also measured over the course of each experiment. Tumor volume was determined using (*W*^2^)(*L*), where *W* represents width and *L* represents length as measured by calipers. Four days after drug treatment, animals were euthanized and tumors were explanted. Tumors were dissociated by shaking for 30 minutes in a 37°C water bath in 5 ml of M199 media (Thermo Fisher 11150067) containing 1 mg/ml each of Collagenase Type 1 (Worthington Biochemical LS004216), Type 2 (Worthington Biochemical LS004204), Type 3 (Worthington Biochemical LS004216), and Type 4 (Worthington Biochemical LS004212), as well as 5 μl DNase I (Thermo Fisher #90083). 15 ml RPMI with 10% FBS was then added to stop the dissociation process, and cells were filtered using a 70 μm strainer and centrifuged at 500 g for 5 minutes. Red blood cells were lysed with 5 ml ACK lysis buffer (Quality Biological #118-156-721) for 7 minutes on ice, then 15 ml of RPMI with 10% FBS was added and cells were centrifuged and prepared for flow cytometry. BFP signal was used to gate for tumor cells.

### Western blotting

The following primary antibodies were used, all at a dilution of 1:1000: p-S345 Chk1 (Cell Signaling Technologies #2348) Chk1 (Cell Signaling Technologies #2360), p-T68 Chk2 (Cell Signaling Technologies #2197), Chk2 (Cell Signaling Technologies #2662), p-S139 H2AX (Cell Signaling Technologies #9718), CD11b (Novus; NB110-89474SS), tubulin (Abcam; ab4074), actin (Cell Signaling Technologies #8457), PU.1 (Cell Signaling Technologies #2258), and vinculin (Cell Signaling Technologies #13901). The following secondary antibodies were used: HRP-linked anti-rabbit IgG (Cell Signaling Technologies #7074, used at 1:5000) and HRP-linked anti-mouse IgG (Cell Signaling Technologies #7076, used at 1:2000). Briefly, cells were washed once with 10 ml ice-cold PBS, pelleted, lysed using LDS sample buffer (Invitrogen; NP0007) with 2% beta-mercaptoethanol (Sigma-Aldrich; M6250) at a concentration of 10 million cells/ml, loaded onto a Qiashredder column (Qiagen; #79656), and sheared by centrifugation at 21300 g for 30 seconds, and the eluate was boiled at 95°C for 5 minutes. Samples were normalized using the Pierce 660 nm protein assay (Thermo Scientific; #22662) with the ionic detergent compatibility kit (Thermo Scientific; #22663) using BSA as a standard. Lysates were resolved by SDS-PAGE and proteins were transferred onto nitrocellulose or PVDF membranes using the iBlot2 Dry Blotting System (Thermo Fisher, IB21001, IB23001).

### Metabolite profiling

After washing in blood bank saline, metabolites from 1 million cells were extracted in 500 µl ice-cold extraction buffer (40:40:20 acetonitrile:methanol:water, 0.1M formic acid) containing 500 nM ^13^C/^15^N-labeled amino acid standards (Cambridge Isotope Laboratories, Inc.). After 3 minutes of extraction, 43 µl of 15% (w/v) ammonium bicarbonate (Sigma) was spiked in. Cells were vortexed for 10 min at 4°C, and then centrifuged at maximum speed for 10 min at 4°C. Samples were dried under nitrogen gas and resuspended in 100 µl HPLC water (Sigma).

Metabolites were measured using a Dionex UltiMate 3000 ultra-high performance liquid chromatography system connected to a Q Exactive benchtop Orbitrap mass spectrometer, equipped with an Ion Max source and a HESI II probe (Thermo Fisher Scientific). External mass calibration was performed using the standard calibration mixture every 7 days. Samples were separated by chromatography by injecting 2 μl of sample on a SeQuant ZIC-pHILIC Polymeric column (2.1 × 150 mm 5 μM, Millipore Sigma). Flow rate was set to 150 μl min, temperatures were set to 25°C for column compartment and 4°C for autosampler sample tray. Mobile Phase A consisted of 20 mM ammonium carbonate, 0.1% ammonium hydroxide. Mobile Phase B was 100% acetonitrile. The mobile phase gradient was as follows: 0–20 min.: linear gradient from 80% to 20% B; 20–20.5 min.: linear gradient from 20% to 80% B; 20.5–28 min.: hold at 80% B. Mobile phase was introduced into the ionization source set to the following parameters: sheath gas = 40, auxiliary gas = 15, sweep gas = 1, spray voltage = 3.1 kV, capillary temperature = 275°C, S-lens RF level = 40, probe temperature = 350°C. Metabolites were monitored in full-scan, polarity-switching, mode. An additional narrow range full-scan (220–700 m/z) in negative mode only was included to enhance nucleotide detection. The resolution was set at 70,000, the AGC target at 1x10^6^ and the maximum injection time at 20 msec.

Relative quantitation of metabolites was performed with XCalibur QuanBrowser 2.2 (Thermo Fisher Scientific) using a 5 ppm mass tolerance and referencing an in-house retention time library of chemical standards. Raw peak areas of metabolites were median normalized by sample after an initial normalization to the abundance of internal ^13^C/^15^N-labelled amino acid standards.

### Cell line generation

For all lentiviral transductions, lentiviral particles were generated by transfecting Lenti-X 293T cells (ATCC) with a 4:2:1:1 ratio of the target:pMDLg:pMD2.G:pRSV-REV plasmids using Transit 293T reagent (Mirus). After 48 hours, supernatant was clarified through a 0.45 μm PES filter and either snap frozen in liquid nitrogen and stored at -80°C or added fresh to target cells. All cells were transduced using 250 μl viral supernatant and 250 μl cells at a concentration of 1e6/ml with polybrene (Millipore, TR-1003-G) added at a final concentration of 8 μg/ml.

To generate ER-Hoxa9 and THP-1 cell lines carrying dCas9, the plasmid pHR-UCOE-SFFV-dCas9-mCherry-ZIM3-KRAB (Addgene plasmid #154473) was transduced into cells as described above. Twenty-four hours after transduction, cells were centrifuged and resuspended in fresh media. After another twenty-four hours, cells within the top half of mCherry expression were single-cell sorted into four 96-well plates containing 75 μl conditioned media. Cell colonies were allowed to grow for 1-2 weeks, and viable colonies were screened for high mCherry expression and low baseline differentiation using flow cytometry.

To generate K562 cell lines carrying dCas9, cells were transduced with lentivirus generated from plasmid UCOE-SFFV-dCas9-BFP-KRAB (Addgene plasmid #85969) as described above. The top half of BFP+ cells was bulk sorted twice and then single cell cloned, and CRISPRi knockdown efficiency was validated using 3 benchmark guides (Palmer et al., 2018).

To generate CRISPRi knockdowns, sgRNA oligos were designed and cloned into the CRISPRi-v2 sgRNA plasmid pU6-sgRNA EF1Alpha-puro-T2A-BFP (Addgene plasmid #60955) using sequences from (Horlbeck et al., 2016). Oligos that were used are listed in Table S5. Lentivirus was generated as above and ER-Hoxa9 and THP-1 dCas9 lines were transduced with lentivirus for 24 hours. Subsequently, cells were recovered for 24 hours then selected using 2 μg/ml (ER-Hoxa9 cells) or 1 μg/ml (THP-1 cells) puromycin. Media was changed and fresh puromycin was added every 1-2 days to maintain selective pressure. After 4 days of selection, BFP positivity was assessed using flow cytometry (usually >97% BFP+). Note that for K562 cells whose dCas9 is fused to BFP, the BFP signal induced by the sgRNA plasmid is much stronger, allowing for unambiguous identification of sgRNA-carrying cells. Knockdown efficiency was tested using quantitative PCR, as described below.

To generate ER-Hoxa9 mCherry-Geminin cells, the mCherry-Geminin fusion gene (Sakaue-Sawano et al., 2008) was cloned into the pLV-Hygro vector (Addgene plasmid #85134), using Gibson cloning. Lentivirus was generated and ER-Hoxa9 cells were transduced as above, then cells were sorted twice for the top half of mCherry-positive cells.

To generate THP-1 PIP-FUCCI cells, cells were transduced with lentivirus generated from the PIP-FUCCI plasmid (Addgene plasmid #138715) as described above. Cells that were both mCherry and mVenus positive were single-cell sorted into 96 well plates, and viable colonies were screened for high mCherry/mVenus expression and low baseline differentiation using flow cytometry. One clone was used for the early S phase isolation experiments described in Figure S7.

### Cell cycle length measurements

12-well cell culture plates were coated with 50 µg/ml poly-D-lysine (Sigma-Aldrich) for 1 hour, washed thrice with PBS, and allowed to air dry in the tissue culture hood for 1 hour. 500 µl ER-Hoxa9 cells expressing mCherry-Geminin were plated in each well at a concentration of 150,000/ml in the presence or absence of drug, and plates were placed into an Incucyte Live-Cell Assay incubator (Sartorius) for 72 hours. Bright-field and red fluorescence images were obtained every 15 minutes. For measurement of S/G2/M phase length, red fluorescence images were processed through an in-house ImageJ pipeline: images were registered using StackReg and background subtracted, then cells were called using Triangle thresholding and mean red fluorescence values within each cell were tabulated for each timepoint. Using an in-house Python pipeline, S phase lengths were extracted from each trace as follows: a threshold for mCherry positivity was calculated as the tenth percentile of mCherry signal, all contiguous blocks of signal between 5 and 40 hours were identified, and the first block of signal that did not overlap the start of image acquisition was called as the S phase length for that cell. G1 lengths were obtained by measuring the amount of time between the first two S/G2/M phases.

### Quantitative PCR

Cells were treated as indicated, and 1 million cells were centrifuged, pelleted, and flash frozen in liquid nitrogen and placed at -80°C. Pellets were processed in groups of 24 using the Zymo RNA Miniprep Kit, RNA was quantified using a Nanodrop, and 500 ng RNA was reverse transcribed into cDNA in a 10 μl reaction using the iScript cDNA synthesis kit (Bio-Rad). cDNA was then diluted 1:10 and 4 μl was used for quantitative RT-PCR using the LuminoCt SYBR Green master mix on a Roche LightCycler 480 machine. *RPS18* and *GAPDH*/*Gapdh* were used as expression controls where indicated. The primers used are listed in Table S5.

### RNA-seq and data analysis

The brequinar ER-Hoxa9 timecourse was performed as described previously and raw data for the E2 withdrawal ER-Hoxa9 timecourse were obtained from GSE84874 (Sykes et al., 2016). For all other RNA-seq experiments, cells were treated as indicated (4 biological replicates per condition), then stained with a final concentration of 1 ng/ml DAPI in growth media, and 1 million live cells were sorted using a BD FACSAria and placed on ice. Immediately following the sort, cells were centrifuged, pelleted, and flash frozen in liquid nitrogen and placed at -80°C. Pellets were processed in groups of 24 using the Zymo RNA Miniprep Kit and quantified using the Invitrogen Quant-it RNA kit. For all experiments not involving K562 cells, RNA was diluted to a concentration of 2 ng/μl and 5 μl of each sample was transferred to a 96 well plate, then poly-A enrichment and cDNA synthesis were performed in parallel as described (Soumillon et al., 2014), followed by library preparation using the Nextera XT kit and paired-end sequencing on an Illumina NextSeq 500. For K562 RNA-seq, RNA was diluted to a concentration of 10 ng/μl and 10 μl of each sample was transferred to a 96 well plate, then poly-A enrichment, cDNA synthesis, and library prep were performed in parallel using the Ultra II Directional RNA-Seq Kit (NEB E7760) and reads were sequenced 2x50bp on an Illumina NovaSeq S4.

Mouse and human data were mapped to mm10 and hg38 genomes, respectively, using STAR v2.7.8a and default parameters. Reads that mapped to transcripts were counted using featureCounts v2.0.1 (with parameters -M -O --fraction --primary). Normalization and differential gene expression analysis was performed using the edgeR package (v4.0.1) in R. Gene ontology enrichment was performed using Enrichr. Gene set enrichment analysis was performed using the fgsea package (v1.28.0) in R.

### CUT&RUN

ER-Hoxa9 and THP-1 cells were treated as indicated, and either directly subjected to CUT&RUN or sorted for early S phase cells (Figure S7) and placed on ice prior to starting the assay. CUT&RUN was then performed using the Epicypher CUTANA CUT&RUN kit, using 500,000 cells for all experiments. The following antibodies were used (1 μl per 50 μl reaction): ER (CST #8644), H3K27ac (EMD Millipore #07-360), RNA polymerase II (Santa Cruz Technologies, sc-47701) and IgG (Epicypher #13-0041). Library construction was performed using the NEBNext Ultra II Library Preparation Kit following the manufacturer’s protocol, and each library was indexed using NEBNext Multiplex Oligos for Illumina. For ER-Hoxa9, the library construction protocol was modified to generate shorter fragments as described in https://www.protocols.io/view/library-prep-for-cut-amp-run-with-nebnext-ultra-ii-wvgfe3w. Libraries were quantified using Qubit and Bioanalyzer and pooled for 2x40bp paired-end sequencing using an Illumina NextSeq 500.

### CUT&RUN data analysis

Mouse and human data were mapped to mm10 and hg38 genomes, respectively, using Bowtie2 v2.4.2 (with parameters --dovetail -I 10 -X 700 --very-sensitive-local --local), then normalized using the bamCoverage tool from deepTools (v3.5.0) (with parameters -bs 1 -e 200). Duplication rates were assessed using Picard, and as they were below 15% for all samples, duplicate reads were not removed.

For H3K27ac CUT&RUN, read counts were tabulated for each peak in the master ATAC-seq peak list (below), and differential peaks were identified using edgeR (v4.0.1). Motif enrichment at sites of differential H3K27ac signal was determined using HOMER (v4.11) (Heinz et al., 2010). Peaks corresponding to promoter sites were identified by filtering out peaks that overlapped a +/-2kb region around promoters defined in Ensembl (org.Mm.eg.db or org.Hs.eg.db), using the package ChIPseeker (Yu et al., 2015). All other peaks were considered to be distal regulatory loci, and they were linked to genes using GREAT (McLean et al., 2010) using the “single nearest gene” rule; associations were retained if the center of the peak was within 100kb of the nearest gene’s TSS. For the early S phase CUT&RUN (Figure S7F-J), H3K27ac peaks were defined using MACS2 (v2.2.9.1) with parameters “-q 0.01 --nomodel”, and the union of peaks that appeared in both replicates of each condition was used as the master peak list. For ER-Hoxa9 CUT&RUN, peaks were defined using MACS2 (v2.2.9.1) using default parameters. For RNA polymerase II CUT&RUN, gene bodies were defined as the region from position +250 (downstream of the promoter) to the transcription termination site (TTS). Heatmaps and metaplots were generated using deepTools (v3.5.1).

### ATAC-seq

For THP-1 cells, cells were stained with a final concentration of 1 ng/ml DAPI in growth media, and 200,000 live cells were sorted using a BD FACSAria and placed on ice. 50,000 cells were then processed using the Active Motif ATAC-seq kit (cat. 53150). Libraries were constructed using the dual index barcodes included in the Active Motif ATAC-seq kit, except that after the PCR amplification, libraries were size selected with 1.5X SPRI beads (Beckman Coulter) instead of 1.2X SPRI beads to retain smaller fragments. Libraries were then quantified using Qubit and Bioanalyzer analysis, pooled, and sequenced using 2x40 paired-end sequencing on an Illumina NextSeq 500.

To generate ATAC-seq libraries for ER-Hoxa9 cells, 50,000 cells were used and libraries were constructed as previously described (Cheloufi et al., 2015; Corces et al., 2017). Briefly, cells were washed in PBS twice, counted and nuclei were isolated from 100,000 cells using 100 μl hypotonic buffer (10 mM Tris pH 7.4, 10 mM NaCl, 3 mM MgCl2, 0.1% NP40) to generate two independent transposition reactions. Nuclei were split in half and treated with 2.5 μl Tn5 Transposase (Illumina) for 30 minutes at 37°C. DNA from transposed nuclei was then isolated and PCR-amplified using barcoded Nextera primers (Illumina). Library QC was carried out using high sensitivity DNA bioanalyzer assay and Qubit measurement and sequenced using paired-end sequencing (PE50) on the Illumina HiSeq 2500 platform.

### ATAC-seq data analysis

Mouse and human data were trimmed using Cutadapt (v3.4) (with parameters - m 20 -q 20 -a CTGTCTCTTATA -A CTGTCTCTTATA), then aligned to the mm10 and hg38 genomes with Bowtie2 v2.4.2 (with parameters --dovetail -I 10 -X 1000 --very-sensitive). Reads in the mitochondrial genome and unannotated chromosomes as well as the UCSC blacklist were removed, then duplicates were removed with samtools markdup (v1.1.2), and the insert size distribution was calculated using Picard CollectInsertSizeMetrics (v2.25.2) and validated to contain nucleosomal peaks. BED file was then constructed with each mate pair as a separate read using bedtools bamtobed (v2.3.0), and peaks were called using MACS2 callpeak (v2.2.7.1) (with parameters --nomodel --shift -100 --extsize 200 --keep-dup all --call-summits). Note that the MACS2 command shifts each read by 100 base pairs towards the 5’ end and extends the read length to 200 base pairs, thus treating the 5’ end of the original read (the Tn5 cut site) as the center of a new “fragment” that is then used for peak calling and other downstream analysis. To normalize tracks for visualization, reads overlapping shared peaks were counted for each BAM file using featureCounts (with parameters -M -O) and normalization factors were calculated from these counts using the calcNormFactors function in the edgeR package (v3.32.1) in R. Visualization was performed using the bedGraphToBigWig utility.

To identify changes in motif accessibility, ChromVAR (Schep et al., 2017) was used. For downstream analysis, peaks from all samples from the same experiment were merged as follows. Peak summits (called using MACS2) were extended by 25bp in each direction, then only peaks that were present in both replicates for each condition (overlapping by at least 50%) were retained. Peak lists were then merged across samples, merging peaks that were within 50bp of each other. ATAC-seq and CUT&RUN reads were then counted against this master peak list using featureCounts (-p -O -M --fraction). Differential ATAC-seq peaks were calculated using edgeR with a log2 fold change threshold of 1 and an adjusted p-value of 0.05. Peaks were annotated to genomic features using ChIPseeker (Yu et al., 2015).

To assess concordance between ATAC-seq and gene expression, the master peak list was subsetted to only include peaks that did not fall within the +/-2kb region surrounding promoters, as described above. GREAT was then run on this set of peaks in single-gene mode, assigning peaks to the single closest transcriptional start site within 100kb. High-complexity genes were defined as genes with at least three associated peaks (Kiani et al., 2022). Shared upregulated genes were considered to be high-complexity genes with a log2 fold change in expression > 1 and FDR < 0.05, and peaks were considered to be more accessible if their ATAC-seq signal was significantly higher (logFC > 1, FDR < 0.05) compared to baseline. To determine the baseline ATAC-seq signal for each peak, the number of reads overlapping each peak in the untreated condition was normalized for the length of the peak.

### scRNAseq

Two experiments in this manuscript used MULTI-seq (McGinnis et al. 2019) to hash cells for single-cell RNA sequencing. In the first experiment, ER-Hoxa9 cells treated with BRQ or E2 withdrawal were grouped together: for BRQ treatments, 1 million cells were washed twice with PBS and treated with 2 μM BRQ for 6, 12, 24, or 48h, and for E2 withdrawal, 1 million cells were washed twice with PBS and then resuspended in media without E2 for 6, 12, 24, or 48h. In the second experiment, erythroid progenitors derived from CD34+ HSPCs were barcoded and combined; 1 million cells treated with DMSO, HU, or APH starting at day 7 were frozen down at day 11 and 14 in 90% FBS and 10% DMSO and cryopreserved, and on the day of MULTI-seq processing, vials were thawed per standard CD34 protocols. To barcode each sample, MULTI-seq was used; the barcode for each sample is included in the supplementary files. The anchor and co-anchor lipid-modified oligos were a gift from Zev Gartner. Briefly, for each condition, 400,000 cells were rinsed twice in PBS and resuspended in 180 μl PBS. 20 μl of a 10X stock of anchor:barcode (per sample: 0.88 μl anchor, 0.44 μl 100 μM barcode, and 20.7 μl PBS) was added to each sample, pipetted gently to mix, and incubated on ice for 5 minutes. Next, 20 μl of a 10X stock of coanchor (per sample: 0.88 μl co-anchor and 21.2 μl PBS) was added to each sample, pipetted gently to mix, and incubated on ice for 5 minutes. 700 μl ice cold 1% BSA in PBS was then added to each sample, and cells were centrifuged at 4°C for 5 minutes. The supernatant was aspirated and the wash was repeated 2X with ice cold BSA in PBS. After the last wash, cells were resuspended in 300 μl PBS/BSA, filtered through a cell strainer, and counted using a Nexcelom cell counter. Cells were adjusted to a concentration of 1 million/ml using ice cold BSA in PBS, and 50 μl of each sample was pooled into one tube and placed on ice. Cells were then loaded into the 10X Genomics microfluidics chip for processing and scRNAseq. For the ER-Hoxa9 experiment, 28,500 cells were loaded in order to obtain enough data for ∼15,000 usable cells, and for the erythroid progenitor experiment, 22,000 cells were loaded in order to obtain enough data for ∼12,000 usable cells (calculation performed at https://satijalab.org/costpercell/).

Library construction proceeded as indicated in the 10X Genomics scRNAseq v3 protocol except as follows (adapted from McGinnis et al. 2019). During cDNA amplification, 1 μl of a 2.5 μM MULTI-seq primer (5’-CTTGGCACCCGAGAATTCC-3’) was added to the master mix. After the 0.6X SPRI clean-up, the supernatant was saved for barcode extraction rather than discarded. To the 60 μl supernatant, 260 μl SPRI beads and 180 μl 100% isopropanol were added for a final ratio of 1.8X beads, pipetted up and down 10x, and incubated at room temperature for 5 minutes. The tube was then placed on a magnetic rack for 5 minutes, supernatant discarded, beads washed 2x with 500 μl 80% ethanol, and eluted with 50 μl buffer EB. This SPRI purification step was repeated once more to further purify the barcode fraction. The barcode fraction was quantified using Qubit, and then library preparation PCR was performed using the Kapa HiFi HotStart ReadyMix with 3.5 ng barcode DNA, 2.5 μl 10 μM universal i5 primer, and 2.5 μl 10 μM TruSeq RPI1 primer in a 50 μl reaction for 8-12 cycles. The PCR product was purified using a final SPRI bead concentration of 1.6X, eluted, and quantified on a Bioanalyzer; a peak between 175 and 200bp was expected. The barcode library was mixed 1:10 with the transcript library and sequenced on an Illumina NextSeq 500.

### scRNAseq data analysis

For ER-Hoxa9 data analysis, FASTQ files were mapped to the mouse cDNA and intronic transcriptomes with the 10x v3 whitelist (all files available at https://github.com/pachterlab/MBGBLHGP_2019/releases) using Kallisto v0.4.6. Samples were then demultiplexed using the deMULTIplex package (v1.0.2) in R (https://github.com/chris-mcginnis-ucsf/MULTI-seq) and doublets were discarded (McGinnis et al., 2019), then processed using Seurat (v4.0). Briefly, raw data were filtered for genes expressed in at least 3 cells and cells that expressed at least 200 genes and a mitochondrial RNA percentage of <12%. To calculate a granulocyte score, z-scores were summed for the following genes: *Spi1*, *Clec4d*, *Tyrobp*, *Csf2rb*, *Mpo*, *Fcer1g*, *Fcgr3*, *Slpi*, and *Sell*. To plot cells around a cell cycle, PCA was performed using only S and G2/M genes from Seurat, and a UMAP was then plotted using this reduced representation. Pseudotimes were calculated using the Slingshot package in R (v2.10.0) to create a trajectory of normal murine neutrophil development (Xie et al., 2020), and bulk RNA-seq data from each timepoint of BRQ or -E2 treatment was then projected to this trajectory to obtain an estimate of the extent of maturation.

For data analysis of human CD34-derived erythroid progenitors, FASTQ files were mapped to the human cDNA transcriptome with the 10x v3 whitelist (files available at https://github.com/pachterlab/kallisto-transcriptome-indices/releases) using Kallisto v0.4.6. Samples were demultiplexed and doublets were discarded as above, then processed using Seurat. Genes expressed in fewer than 3 cells or cells with fewer than 200 distinct transcripts or with a mtRNA percentage of >10% were discarded. For the normal bone marrow reference, raw data from four cell subsets (CD34+ CD164+, CD34-CD164^hi^, CD34^lo^ CD16^hi^, and CD34-CD164^lo^) were downloaded from GSE117498, and genes expressed in fewer than 3 cells or cells with fewer than 200 distinct transcripts or an mtRNA percentage of >15% were discarded. The reference data was integrated with our scRNAseq data using the FindTransferAnchors and MapQuery functions in Seurat, and projections of our scRNAseq data to the reference map were used for subsequent analyses. Pseudotimes for cells along the erythroid trajectory were calculated using the Slingshot package in R (v2.10.0) and expression of specific genes was plotted using the geom_smooth function.

### CRISPRi screen

CRISPRi screens were performed using ER-Hoxa9, THP-1, or K562 dCas9-KRAB cells generated as above. For the genome-wide screens, libraries with 5 sgRNAs per gene (mCRISPRi-v2; the “top5” library from Addgene #1000000092) or 10 sgRNAs per gene (hCRISPRi-v2; both the “top5” and “supp5” libraries from Addgene #1000000090) were prepared as described in (Palmer et al., 2018) with several modifications. Briefly, each plasmid library was electroporated into MegaX cells (Invitrogen #C640003), maintaining at least 2000X representation, and plasmid was maxiprepped (Qiagen); the libraries were sequenced as described below to confirm even representation of guides. LentiX cells were transfected with plasmid as described above, and 48 hours later supernatant was collected and frozen in aliquots at -80°C. ER-Hoxa9 and THP-1 cells were always maintained between 250,000 and 1 million cells/ml, and two biological replicates were performed starting at the lentiviral infection step. 200 million cells (THP-1) or 150 million cells (ER-Hoxa9) were infected for 24 hours with lentiviral supernatant with 8 μg/ml polybrene, achieving an MOI of between 0.2 and 0.35. Lentiviral media was then replaced with fresh media, and 24 hours later, 2.5 μg/ml puromycin (ER-Hoxa9) or 1 μg/ml puromycin (THP-1) was added to select sgRNA-expressing cells. Media was exchanged and fresh puromycin was added 24 and 48 hours afterwards. Four days after puromycin treatment, viability was >90% and BFP positivity was >90% for both replicates. At this point, puromycin was removed and 200 million (THP-1) or 50 million cells (ER-Hoxa9) were treated with DMSO or BRQ (500 nM for THP-1 cells). 48 hours later, cells were concentrated to 20 million cells/ml in fresh media and stained with antibodies and propidium iodide, and at least 15 million (THP-1) or 10 million (ER-Hoxa9) BFP-positive live cells from both the top and bottom 25% of flow marker expression (CD11b or CD14 for THP-1; CD11b or GFP for ER-Hoxa9) were sorted using a BD FACSAria, pelleted, and frozen at -80°C for later processing. Aliquots of 15-20 million unsorted cells before and after treatment were also pelleted and frozen as controls for guide enrichment or dropout.

For the sublibrary screen, the top and bottom 300 hits of the THP-1 BRQ CD11b screen were selected, and all ten sgRNAs from the hCRISPRi-v2 library were incorporated into the sublibrary, along with ten sgRNAs for 894 randomly chosen genes and 87 hand-curated genes as well as 490 control sgRNAs, and oligos were synthesized as a pool (Twist Biosciences) (Table S4). Oligos were cloned into pU6-sgRNA EF1Alpha-puro-T2A-BFP (Addgene plasmid #60955) using methods described in Gilbert et al. 2014; briefly, oligos were amplified and digested with BstXI and BlpI, the 33bp insert containing the sgRNA sequence was size selected on a 10% PAGE gel and purified, and ligation into the linearized backbone was performed at a 1:1 molar ratio of insert to vector. Ligated plasmids were electroporated into MegaX cells at approximately 5000X library coverage, maxiprepped, and transfected into LentiX cells. 25 million THP-1 and K562 dCas9-KRAB cells were infected for 24 hours in duplicate with lentivirus and 8 µg/ml polybrene (MOI of between 0.25 and 0.35 for both cell lines), lentiviral media was replaced with fresh media, and 24 hours later puromycin selection was started at a concentration of 1 µg/ml. Cells were maintained at a concentration of between 250,000 and 1 million cells/ml and with a minimum of 500X library coverage throughout the selection, and puromycin was refreshed as above. After four days of selection, >90% of cells were BFP positive. 8 million K562 or THP-1 cells were treated with DMSO and 16 million THP-1 cells were treated with 500 nM BRQ for 48 hours (all screens performed in duplicate). 48 hours later, cells were concentrated to 20 million/ml and stained with antibodies and propidium iodide, and the top and bottom 25% of CD235a-expressing K562 cells or the top and bottom 25% of CD11b or CD14-expressing THP-1 cells were sorted and frozen at -80°C for later processing. At least 2 million BFP+ live cells were collected per gate to achieve a minimum of 100X library coverage. Aliquots of 5 million unsorted cells before and after each treatment were also pelleted and frozen as controls for guide enrichment or dropout.

To obtain amplicons for sequencing, genomic DNA from each cell pellet was extracted using the Machery Nagel Nucleospin Blood L kit (for genome-wide screens) or the Qiagen DNA Blood Mini kit (for sublibrary screens) following the manufacturer’s instructions except for an overnight incubation at 70°C and elution of DNA into 200 μl 10mM Tris (pH 8.5). Guides were then PCR amplified using custom i7 and custom i5 primers indexed for each sample, with reactions split such that each 50 μl PCR reaction contained no more than 2.5 μg DNA in a 20 μl volume. For the THP-1 genome wide CRISPRi screens, each library was amplified using one of the p5 primers oBD370-375 and one of the p7 primers oBD367-368 (Table S5). For the ER-Hoxa9 genome-wide CRISPRi screens and all sublibrary screens, each library was amplified using one of the p5 primers oSV104-111 and one of the p7 primers oSV112-119 (Table S5). PCR was performed using the NEBNext Ultra Q5 2X Master Mix with the following cycling parameters: initial denaturation at 98°C for 2 minutes, then 24 cycles of denaturation at 98°C for 15 seconds and annealing/extension at 65°C for 75 seconds, then a final extension of 65°C for 5 minutes. All reactions for each pellet were pooled, and successful amplification was verified by running 10 μl of the PCR on an agarose gel to obtain an expected band between 250 and 260bp. 200 μl of the PCR reaction (roughly corresponding to 1 μg of amplicon) was then purified using a double-sided selection with AMpure XP beads (Beckman Coulter). First, to remove genomic DNA, 0.5X volume of beads (100 μl) was added to the PCR reaction, mixed well, and incubated at room temperature for 10 minutes, and beads were placed on a magnet and supernatant was saved. Second, to remove primer dimers, 135 μl beads were added to 290 μl of supernatant (final bead volume of 1.2X), mixed well, and incubated at room temperature for 10 minutes, and beads were placed on a magnet and supernatant was discarded. DNA was then eluted from the beads using 20 μl of 0.1X TE (pH 8.5), quantified using Qubit and Bioanalyzer, and pooled for 1x75 sequencing on an Illumina NextSeq 500. For the THP-1 genome-wide CRISPRi screen, we used an equimolar mix of custom read 1 primers, numbered oBD379-384 (Table S5). For the ER-Hoxa9 genome wide screens and all sublibrary screens, we used primer oBD582 (Table S5).

For data analysis, FASTQ files were demultiplexed and barcode sequences were matched to the master list of CRISPRi barcodes; only perfect matches were retained. Raw counts were subsequently processed using MAGECK (v0.5.9) to obtain sgRNA-level and gene-level scores and their corresponding p-values. Hits were defined using the thresholds described in the text and figure legends. Gene ontology analysis was performed using the enrichR package (v3.2) in R. To determine the effect of each knockdown on proliferation rate, we used MAGECK to compare the abundance of sgRNAs in a cell pellet obtained at day 6 to the abundance of sgRNAs in the plasmid library used to generate the lentiviral pool.

### Perturb-seq analysis

We normalized and filtered cells and combined all cells for a given knockdown into a “pseudobulk” transcriptome (Replogle et al., 2022). Replication stress-related genes were identified by taking the intersection of genes from GO ontology #0006261 (DNA-templated DNA replication), GO #0045005 (DNA-templated DNA replication maintenance of fidelity), GO #1903932 (regulation of DNA primase activity), Reactome #R-HSA-905320 (nucleotide biosynthesis), and Wikipathway #4022 (pyrimidine biosynthesis), and genes whose symbols started with “POLR”, “PELO”, or “PNPT1” were removed because they were unrelated to DNA replication. Myeloid scores were calculated by summing z-scores for the following genes: *CSF3R*, *LST1*, *CD33*, *LYZ*, *AIF1*, *CD55*, and *SPI1*. Erythrocyte scores were similarly calculated using the following genes: *HBG1*, *GYPA*, *ERMAP*, *ALAS2*, *HBA1*, *KLF1*, and *SLC25A37*. For each knockdown, the fraction of S phase cells was calculated by assigning cells to the most likely phase based on expression of experimentally-derived genes specific for each phase, and the relative proliferation was obtained from the abundance of each guide at day 8 relative to the plasmid library; these methods are described in more detail in (Replogle et al., 2020).

## Data and code availability

All sequencing data generated in this study are deposited on GEO under accession number GSEXXXXXX. All code is available on GitHub at https://github.com/bdo311/hsu_do_2022.

## ACKNOWLEDGEMENTS

We thank members of the Vander Heiden and Sykes laboratories for scientific discussions and experimental support. We are grateful to Jette Lengefeld, Chris McGinnis, Russell J. H. Ryan, Teemu Miettinen, Justine Rutter, Stuart Levine, M. Andres Blanco, Amy N. Sexauer, Yadira M. Soto-Feliciano, Eliezer Calo, Adam J. Rubin, Yatrik Shah, Christopher W. Ng, Kathleen W. Higgins, Divya Iyer, and Nan Liu for helpful suggestions, critical feedback, and experimental advice and assistance. We also thank the Koch Institute Flow Cytometry Core, BioMicro Center, Microscopy Core, and Histology Core as well as the Whitehead Metabolomics Core and the ICCB-Longwood Screening Facility, for their support and assistance, as well as M. Taipale, M. Trakala, and T. Meyer for sharing plasmids. Certain figures were created with Biorender.com.

## AUTHOR CONTRIBUTIONS

B.T.D., P.P.H., and M.G.V.H. conceived of the project. P.P.H. conducted chemical screens with the help of N.A., I.S.H., and R.F. P.P.H., B.T.D., S.Y.V., and Z.W. conducted the majority of experiments in the paper. B.T.D. and S.Y.V. conducted CRISPRi screens with the help of K.L.A., S. Block, and A.M.D. B.T.D. performed the majority of computational analysis with the help of S.Y.V. C.C. provided reagents and assisted with CRISPRi screen design. X.A.S. generated the FUCCI construct, provided expertise with DNA fiber analysis, and performed DNA comet assays. D.R.S. conducted irradiation experiments. T.H., H.Z., and J.M. conducted *in vivo* experiments. B.T.D. performed erythroid differentiation experiments with S.N. S. Bjelosevic performed PDX experiments under the guidance of K.S. J.P. performed primary AML experiments under the guidance of Y.P. S.C. and D.B.S. performed the RNA-seq and ATAC-seq timecourse in ER-Hoxa9 cells. J.M.R. and J.S.W. provided Perturb-seq data, performed computational analysis, and provided expertise. B.T.D. and P.P.H. wrote the paper with input from M.G.V.H. and all co-authors. D.B.S. and M.G.V.H. obtained funding.

## FUNDING

This work was supported in part by a Koch Institute/DFHCC Bridge Grant and the Koch Institute Support (core) Grant P30-CA14051 from the National Cancer Institute. M.G.V.H. acknowledges support from R35CA242379, the Ludwig Center at MIT, the MIT Center for Precision Cancer Medicine, and a Faculty Scholar award from HHMI. P.P.H. was supported in part by 2T32CA071345-21A1. B.T.D. was supported by F30HL156404 from NHLBI and T32GM007753 from NIGMS. R.F. was supported by the Knut and Alice Wallenberg Foundation (KAW 2019.0581). J.M.R. was supported by F31NS115380 from NINDS. K.L.A. was supported by DGE-1122374 from NSF, F31CA271787 from NCI, and T32GM007287 from NIH. A.M.D. was supported by a Jane Coffin Childs Postdoctoral Fellowship. C.C. was supported by U54-CA225088 from NIH/NCI. X.A.S. was sponsored by Center for Prostate Disease Research, Henry Jackson Foundation and National Cancer Institute. J.P. was supported by 5T32HL007574-38 from NHLBI, the Boston Children’s Hospital Translational Research Program (TRP), and the Wong Family Award in Translational Oncology. S.B was supported by a Leukemia & Lymphoma Society (LLS) Fellowship. I.S.H. was supported by the NIH (R01CA269813) and the American Cancer Society (RSG-23-971782-01-TBE).

## CONFLICTS OF INTEREST

D.B.S. is a co-founder and holds equity in Clear Creek Bio. M.G.V.H. discloses that he is on the scientific advisory board of Agios Pharmaceuticals, iTeos Therapeutics, Drioa Ventures, Sage Therapeutics, and Auron Therapeutics. P.P.H. has consulted for Auron Therapeutics. J.S.W. serves as an advisor to and/or has equity in KSQ Therapeutics, Maze Therapeutics, and 5AM Ventures, all unrelated to the present work.

J.M.R. consults for Maze Therapeutics, Waypoint Bio, and Third Rock Ventures. I.S.H. reports financial support from Kojin Therapeutics and consulting fees for Ono Pharma USA. All other authors declare that they have no conflict of interest.

## Notes

### Competing Interest Statement

The authors have declared no competing interest.

### Summary of Updates

All figures have been revised, along with supplemental figures and the author list.

