## Supplemental Figures for "Nucleotide depletion promotes cell fate transitions by inducing DNA replication stress"

**Figure S1. Proliferation and metabolic effects of selected compounds from chemical screen.** Related to Figure 1.

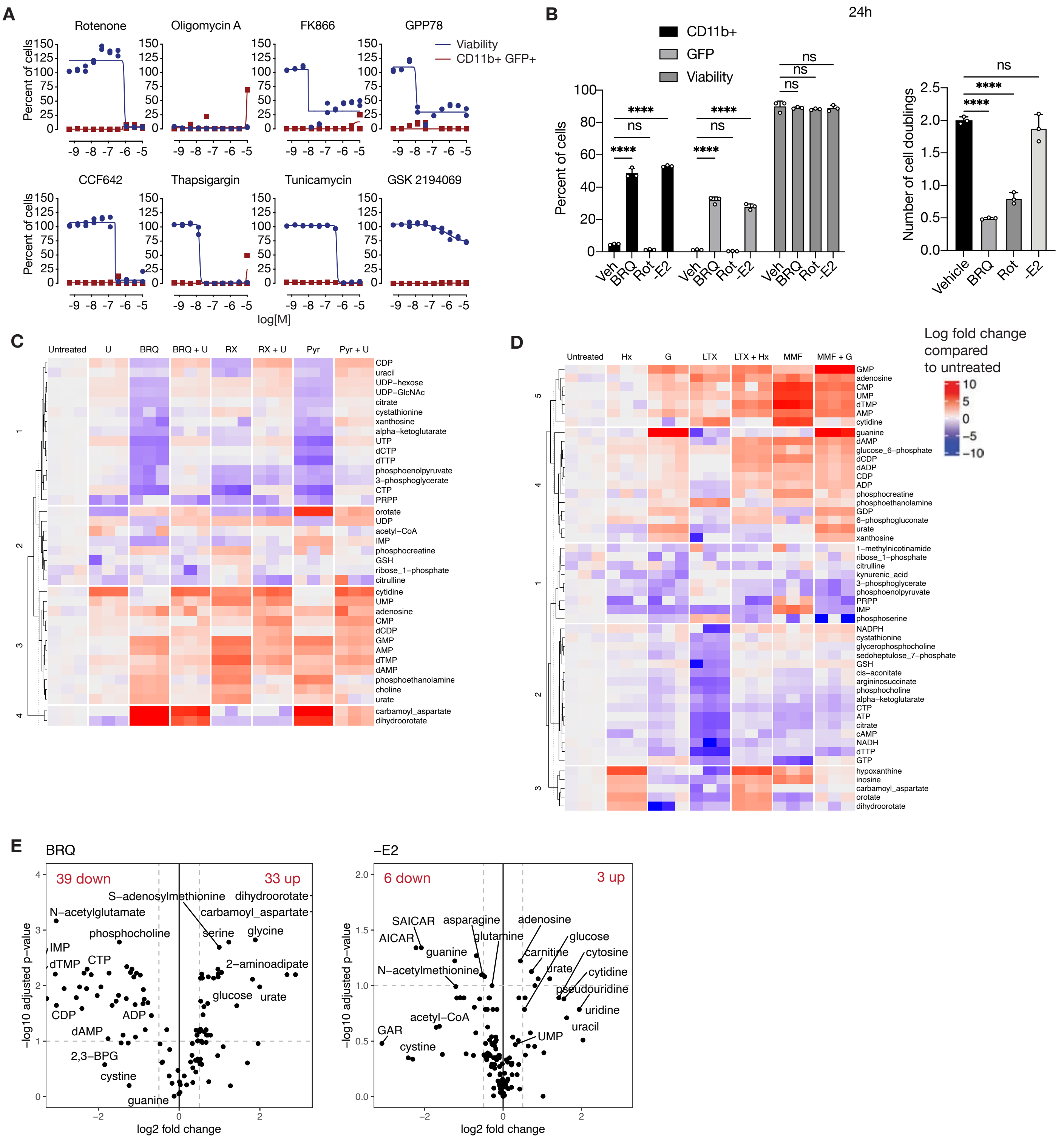

**Figure S2. Cell fate, nucleotide metabolomics, and replication stress signaling in ER-Hoxa9 cells treated with replication stress inducers.** Related to Figure 2.

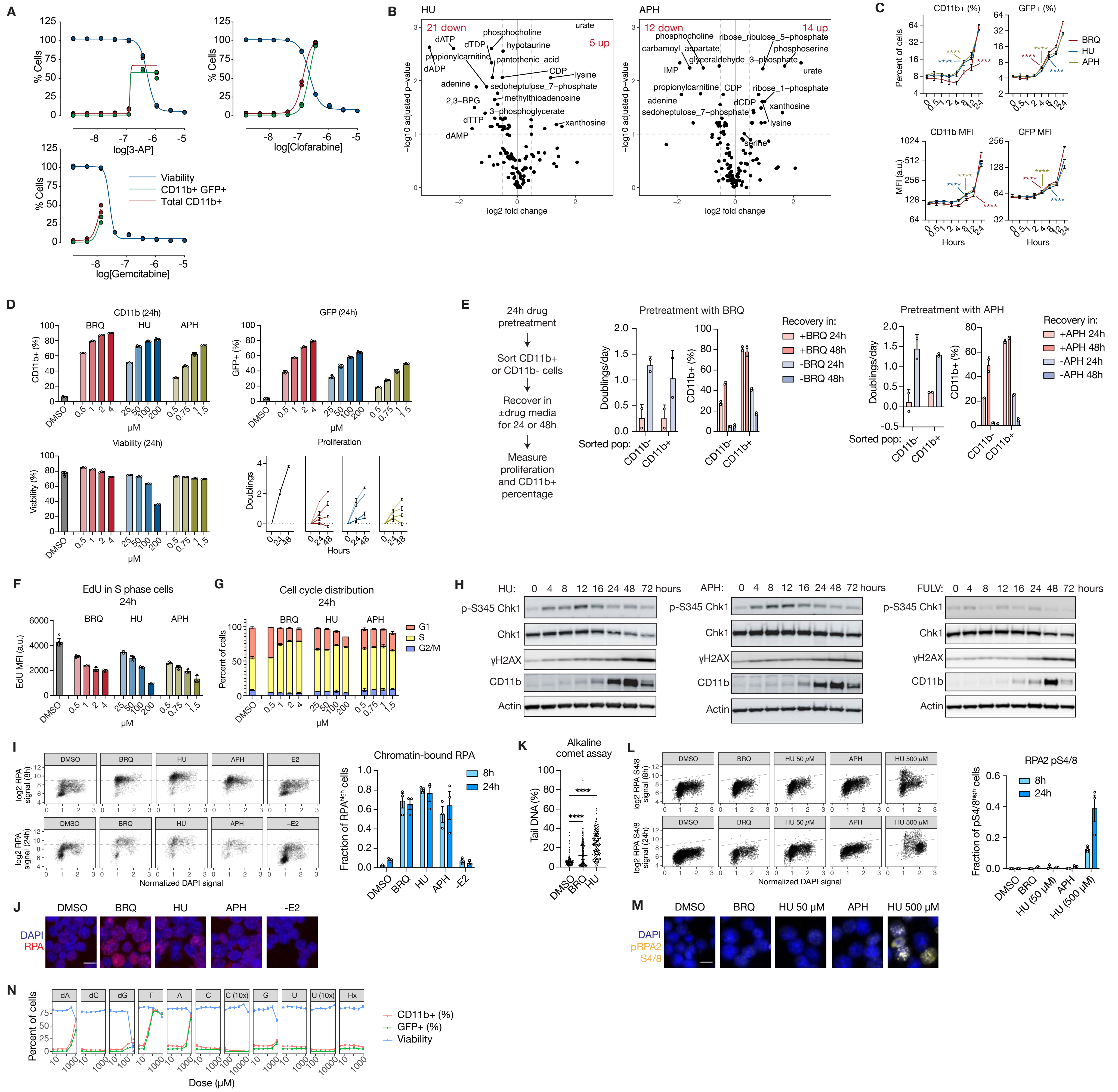

**Figure S3: Effects of nucleotide depletion and replication stress in human leukemia models.** Related to Figure 3.

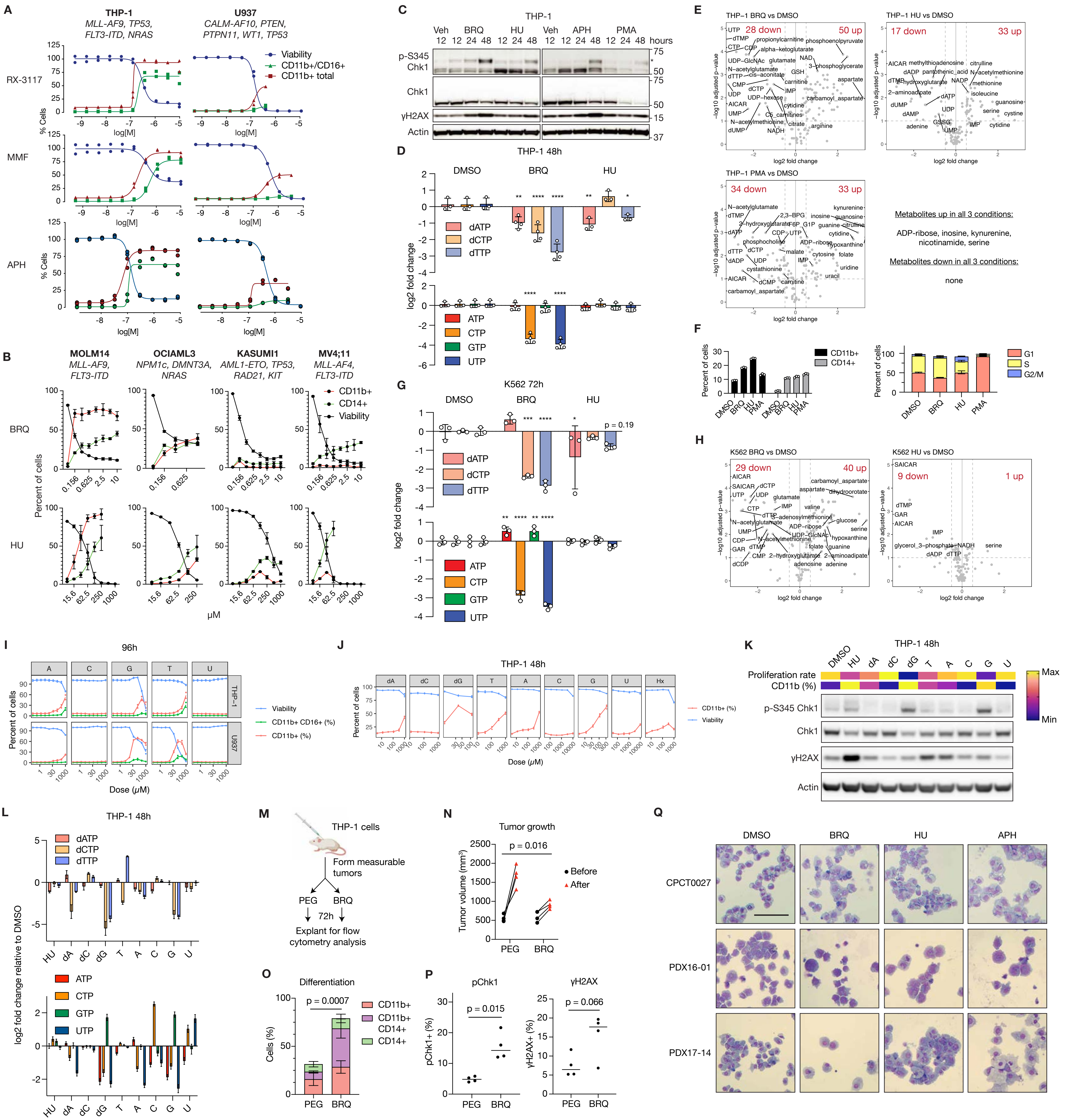

**Figure S4. Analysis of CRISPRi screens.** Related to Figure 4.

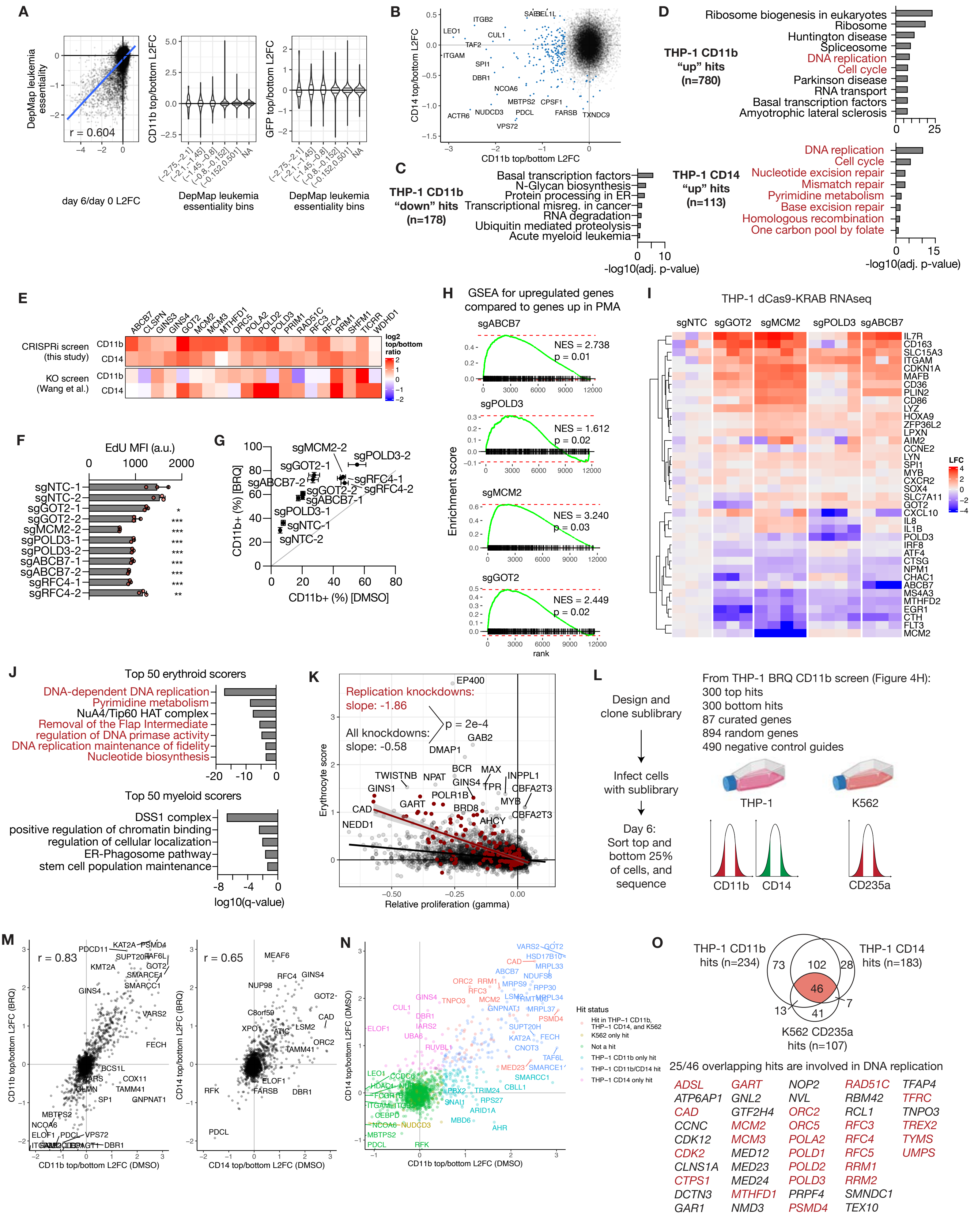

**Figure S5. Relationships between replication stress-induced cell state changes and cell cycle phase, length, replication stress signaling, and DNA damage response.** Related to Figure 5.

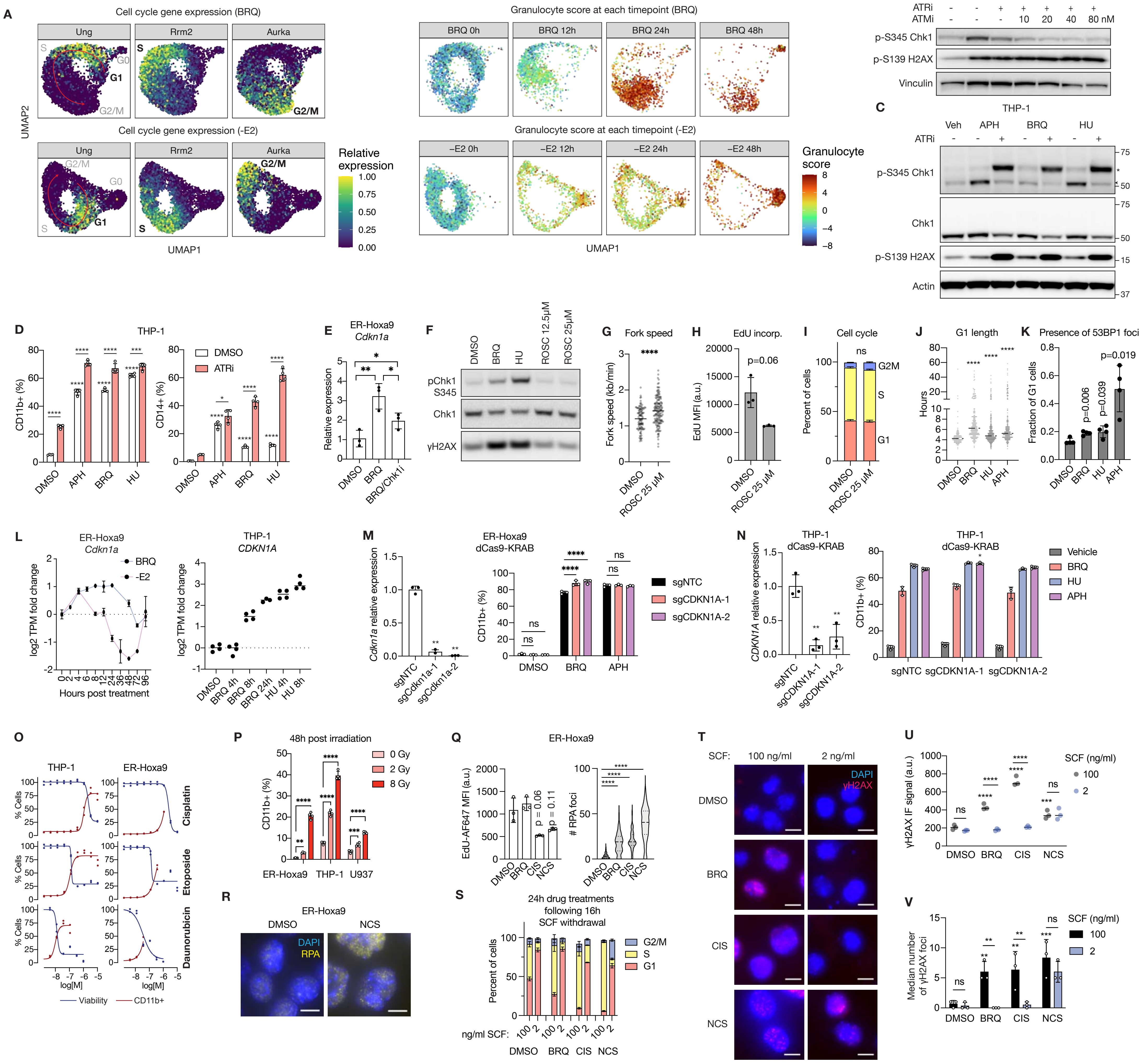

**Figure S6. Analysis of gene expression changes during replication stress.** Related to Figure 6.

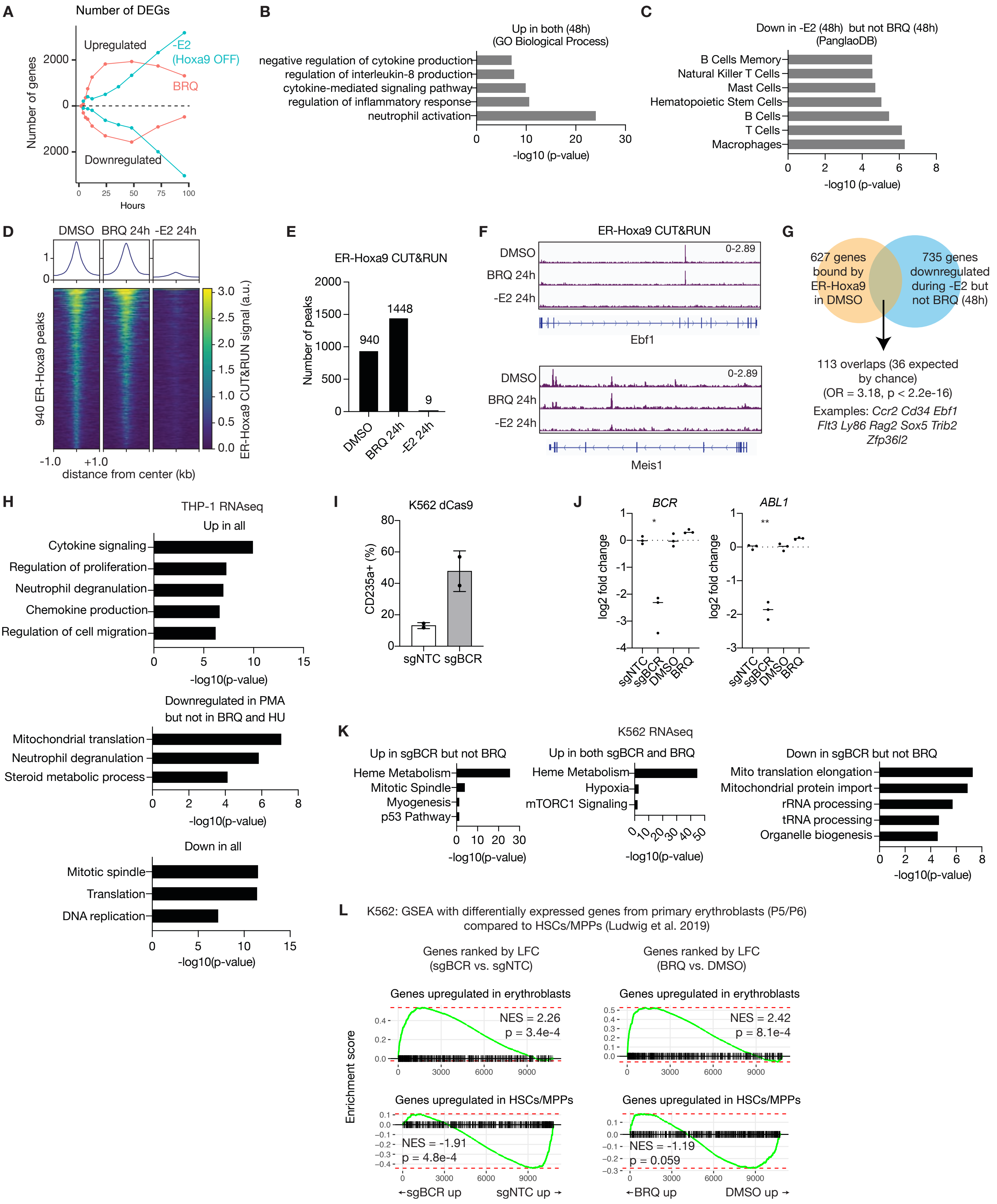

**Figure S7. Analysis of RNA polymerase, H3K27ac, and ATAC-seq changes during replication stress.** Related to Figure 6.

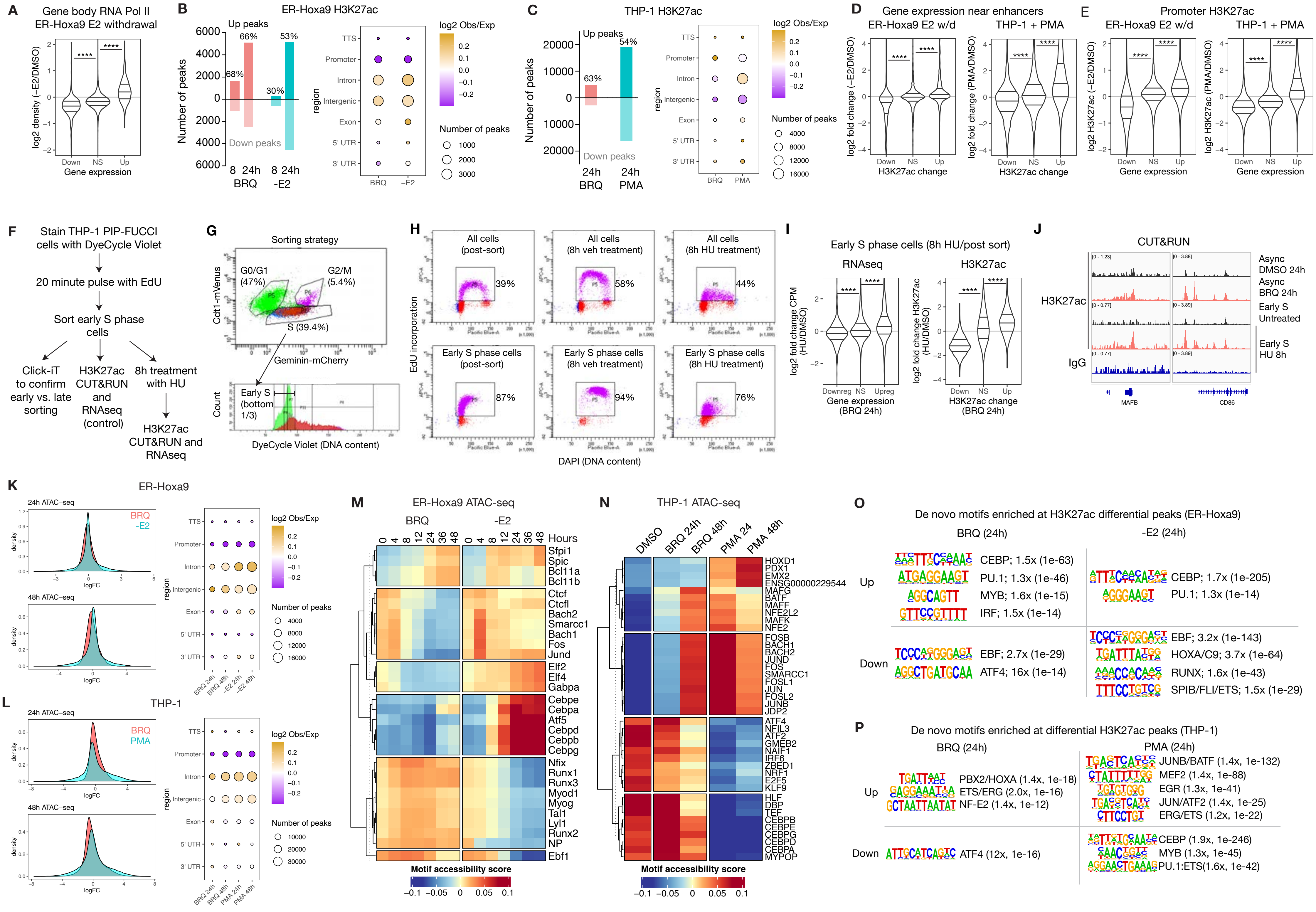

**Figure S8. Preferential activation of primed, lineage-specific gene expression programs during replication stress.** Related to Figure 7.

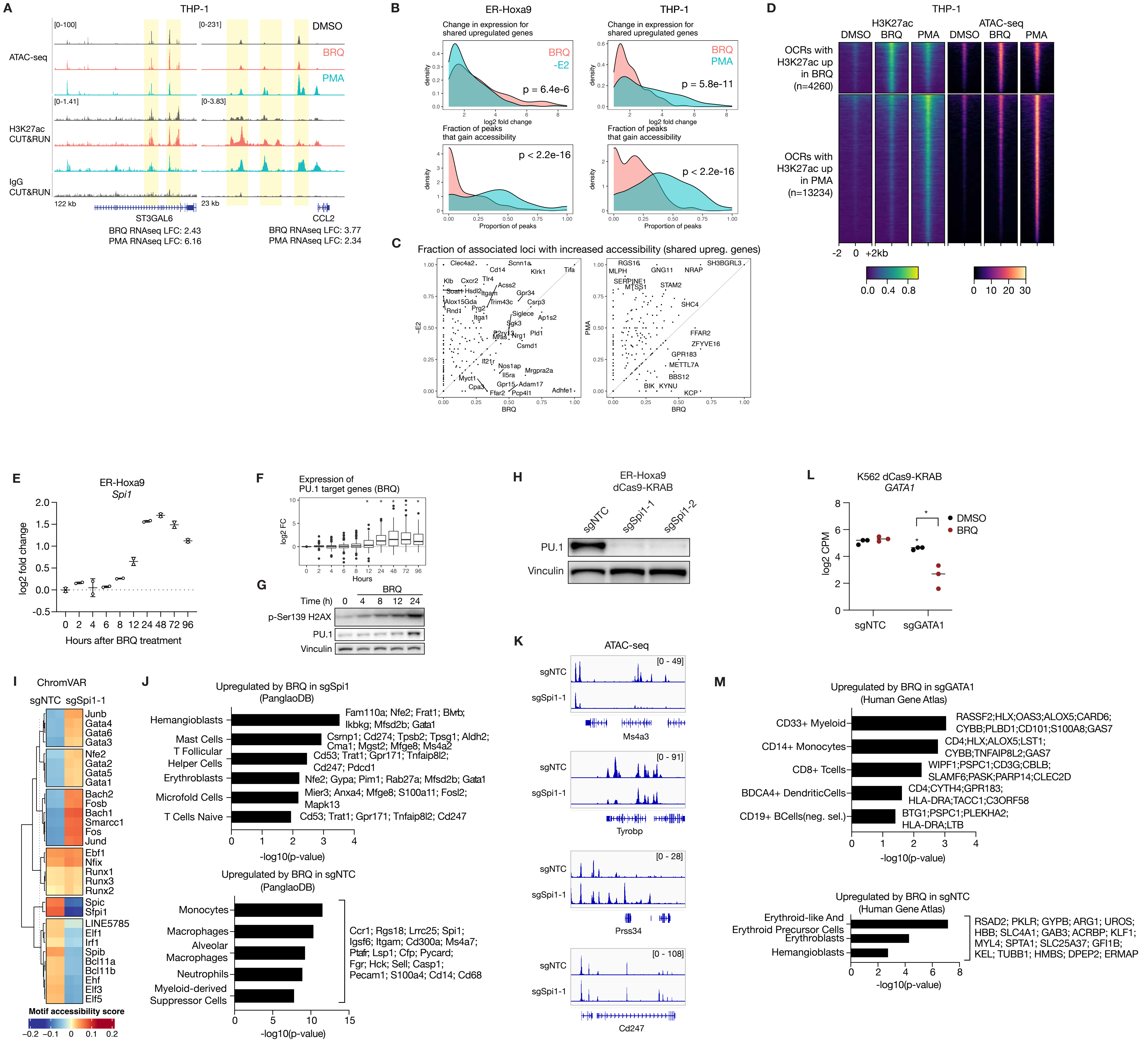
