## Supplemental Information for "Nucleotide depletion promotes cell fate transitions by inducing DNA replication stress"

### SUPPLEMENTAL TABLES

Table S1. Metabolic screen data.

Table S2. Metabolite profiling data.

Table S3. CRISPRi screen data and comparison to Wang et al. (2021) and Replogle et al. (2022) screens.

Table S4. RNAseq analysis for ER-Hoxa9, THP-1, and K562 cells and gene ontology analysis.

Table S5. Primers used in study.

### SUPPLEMENTAL FIGURE LEGENDS

**Figure S1. Proliferation and metabolic effects of selected compounds from the chemical screen.** Related to Figure 1.

A. Percent viability (in blue) and differentiation as measured by CD11b and GFP expression (in red) for ER-Hoxa9 GMP cells treated with the indicated metabolic inhibitors for 96h. Data from two technical screening replicates are plotted.
B. Percentage of viable and CD11b+ or GFP+ cells (left) or number of population doublings (right) for ER-Hoxa9 GMP cells treated for 24h with DMSO, 1 μM BRQ, 200 nM rotenone (Rot), or E2 withdrawal (-E2). (****), $p < 0.0001$ (one-way ANOVA with Dunnett's multiple comparisons test)
C. Heatmap of relative metabolite levels following inhibition of pyrimidine nucleotide synthesis in the absence or presence of uridine addition as indicated. Data are displayed as log fold change values compared to the median of the untreated normalized area under the peak. Only metabolites considered variable (see STAR Methods) are shown. U, uridine. BRQ, brequinar. RX, RX-3117. Pyr, pyrazofurin.
D. Heatmap of relative metabolite levels following inhibition of purine nucleotide synthesis in the absence or presence of guanine or hypoxanthine addition as indicated. Data are displayed as log fold change values compared to the median of the untreated normalized area under the peak. Only metabolites considered variable (variance across all treatments > 1.5) are shown. Hx, hypoxanthine. G, guanine, LTX, lometrexol. MMF, mycophenolate mofetil.
E. Log2 fold change of metabolites after 24 hours of BRQ (left) or -E2 (right) treatment compared to DMSO (n=3). P-values from two-sample Student's t-test with Benjamini-Hochberg multiple hypothesis correction. The number of statistically significant changes in metabolites (fold change >2 or <0.5 and adjusted p-value < 0.01) is displayed.

**Figure S2. Cell fate, nucleotide metabolomics, and replication stress signaling in ER-Hoxa9 cells treated with replication stress inducers.** Related to Figure 2.

A. Percentage of viable and CD11b+ or CD11b+/GFP+ ER-Hoxa9 cells after treatment with the indicated amount of the RNR inhibitors 3-AP, gemcitabine, or clofarabine for 96 hours (n=2).
B. Log2 fold change of metabolites after 24 hours of treatment with 50 μM HU (left) or 1 μM APH (right) treatment compared to DMSO (n=3). P-values from two-sample Student's t-test. The number of statistically significant changes in metabolites (fold change >2 or <0.5 and adjusted p-value < 0.01) is displayed.
C. Percentage of positive CD11b+/GFP+ cells (top) or CD11b/GFP median fluorescence intensity (MFI) (bottom) for cells treated with 1 μM BRQ, 50 μM HU, or 1 μM APH for the indicated times (n=3). (****) p < 0.0001. P-values are from two-way ANOVA with the Šidák correction for multiple comparisons and are only denoted at the earliest timepoint for which p < 0.0001.
D. (first 3 panels) Percentage of CD11b+, GFP+, or viable ER-Hoxa9 cells after treatment with the indicated doses of drug for 24 hours. (bottom right panel) Number of populations doublings at 24h and 48h for these drug doses.
E. (Left) Schematic of pretreatment, sorting, and recovery experiment. (middle) Doublings per day and percentage of CD11b-positive cells for each sorted subpopulation recovered in media with or without 1 μM BRQ for 24h or 48h. (right) Same as middle, but with 1 μM APH. (n=2).
F. S-phase EdU median fluorescence intensity for ER-Hoxa9 cells treated with the indicated drug doses for 24h (n=3).
G. Percentage of G1, S, or G2/M ER-Hoxa9 cells after treatment with the indicated drug doses for 24h (n=3). Percentages do not all add up to 100% because of the presence of sub-2n cells that were not gated as G1.
H. Immunoblots of ER-Hoxa9 cells treated with 50 μM HU, 1 μM APH, or 10 μM FULV (fulvestrant) for the indicated timepoints.
I. (left) Quantitative image-based cytometry (QIBC) of DAPI signal (x-axis) versus chromatin-bound RPA (y-axis) for ER-Hoxa9 cells treated with DMSO, 1 μM BRQ, 50 μM HU, 1 μM APH, or -E2 for 8 or 24 hours. Data from one of three biological replicates is displayed. (right) Percentage of cells with CB-RPA signal higher than the threshold (n=3).
J. Representative immunofluorescence images of cells from (I), stained for DAPI (blue) or chromatin-bound RPA (red). Scale bar represents 20 μm.
K. Percentage of tail DNA from the alkaline comet assay for cells treated with DMSO, 1 μM BRQ, or 50 μM HU for 12 hours. At least 150 cells from each condition were analyzed. (****) p < 0.0001, one-way ANOVA with Dunnett's multiple comparisons test.
L. QIBC plots (left) and quantifiication (right) as in (I) for DAPI and phospho-S4/8 RPA2 signal in ER-Hoxa9 cells treated with DMSO, 1 μM BRQ, 50 μM HU, 1 μM APH, or 500 μM HU.

M. Representative immunofluorescence images of cells from (L), stained for DAPI (blue) or chromatin-bound pS4/8 RPA2 (yellow). Scale bar represents 20 μm.

N. Percentage of viable, CD11b+, and GFP+ ER-Hoxa9 cells after treatment with the indicated dose of nucleobase/nucleoside for 24h (n=3). Each titration represents a three-fold serial dilution starting from the highest dose indicated. Mean and SD are plotted.

**Figure S3. Effects of nucleotide depletion and replication stress in human leukemia models.** Related to Figure 3.

A. Percentage of CD11b+, CD11b+/CD16+, and viable THP-1 (left) or U937 (right) cells upon treatment with RX-3117, MMF, or APH for 96h.

B. Percentage of CD11b+, CD14+, and viable MOLM14, OCIAML3, KASUMI1, or MV411 cells upon treatment with the indicated doses of BRQ (top) or HU (bottom) for 48h.

C. Immunoblots of THP-1 cells treated with vehicle or 1 μM BRQ, 100 μM HU, or 100 nM PMA for the indicated timepoints. The asterisk represents a hyperphosphorylated Chk1 species that occurs alongside DNA damage and is likely due to autophosphorylation as well as phosphorylation of serine 345 by ATM and other kinases.

D. Log fold change of intracellular levels of dATP, dCTP, or TTP (top) and ATP, CTP, GTP, or UTP (bottom) in THP-1 cells following 48h of treatment with DMSO, 1 μM BRQ, or 100 μM HU (n=3). (ns) not significant, (*) $p < 0.05$, (**) $p < 0.01$, (***) $p < 0.001$, and (****) $p < 0.0001$, and adjusted p-values were calculated using a two-way ANOVA with the Šidák correction for multiple comparisons.

E. Log2 fold change of metabolites after 24 hours of treatment with 500 nM BRQ, 100 μM HU, or 100 nM PMA compared to DMSO (n=3). P-values from two-sample Student's t-test. The number of statistically significant changes in metabolites (fold change >2 or <0.5 and adjusted p-value < 0.01) is displayed. Metabolites statistically significantly increased in all treatments are listed.

F. Percentage of CD11b+ and CD14+ THP-1 cells (left) and percentage of cells in each cell cycle phase (right) after treatment with DMSO, BRQ, HU, or PMA for 24 hours.

G. Log2 fold change of intracellular levels of dATP, dCTP, or TTP (top) and ATP, CTP, GTP, or UTP (bottom) in K562 cells following 24h of treatment with DMSO, 250 nM BRQ, or 125 μM HU (n=3). (*) $p < 0.05$, (**) $p < 0.01$, (***) $p < 0.001$, and (****) $p < 0.0001$, and adjusted p-values were calculated using a two-way ANOVA with the Šidák correction for multiple comparisons.

H. Log2 fold change of metabolites after 24 hours of treatment with 250 nM BRQ or 125 μM HU compared to DMSO (n=3). P-values from two-sample Student's t-test. The number of statistically significant changes in metabolites (fold change >2 or <0.5 and adjusted p-value < 0.01) is displayed. Metabolites statistically significantly increased in all treatments are listed.

I. Percentage of viable, CD11b+, and CD11b+ CD16+ THP-1 or U937 cells after treatment with the indicated dose of nucleobase/nucleoside for 96h (n=2). Each titration represents a three-fold serial dilution starting from the highest dose indicated. Mean and SD are plotted.

J. Percentage of viable and CD11b+ THP-1 cells after treatment with the indicated dose of nucleobase/nucleoside for 48h (n=3). Each titration represents a three-fold serial dilution starting from the highest dose indicated. Mean and SD are plotted.

K. Proliferation rates, surface marker expression, and immunoblots for THP-1 cells treated for 48 hours with DMSO, 100 μM HU, or the indicated nucleobase/nucleoside (all at 1 mM except for dG, which was supplemented at 100 μM, and G, which was supplemented at 300 μM). Proliferation rates and percentage of CD11b+ cells are row-normalized.

L. Log2 relative abundance of dNTPs (left) and NTPs (right) for THP-1 cells treated for 48h as in (K), compared to DMSO control (n=3, mean +/- SD displayed).

M. Schematic showing experiment to assess response to brequinar in AML cells in mice. Five million THP-1 cells expressing BFP were injected subcutaneously into the flanks of nude mice, and once tumors formed a measurable size (~11 days) mice were treated with vehicle (PEG) or 50 mg/kg BRQ IP for 3 days, followed by tumor harvest and analysis of BFP+ cells by flow cytometry.

N. Assessment of tumor size before and after vehicle (PEG) or BRQ treatment (n=4). P values from Student's t-test.

O. Percentage of CD11b+ CD14-, CD11b+ CD14+, or CD11b- CD14+ THP-1 cells in tumors as assessed by flow cytometry (n=4). P values from Student's t-test.

P. Percentage of pChk1-positive (left) or γH2AX-positive (right) THP-1 cells in tumors as assessed by flow cytometry (n=4). P values from Student's t-test.

Q. Cytospins and May-Grunwald-Giemsa stains of AML PDX cells treated with the indicated drugs for 72h.

**Figure S4. Analysis of CRISPRi screens.** Related to Figure 4.

A. Analyses of the relationship between proliferation rate and CD11b and GFP enrichment. (Left) Scatterplot of the day 6/day 0 log2 fold change (L2FC) versus the median Dependency Map (DepMap) essentiality score within leukemia cell lines for each gene. The L2FC is calculated by taking the median log2 fold change (L2FC) of sgRNA counts for each gene between day 6 of the screen versus corresponding sgRNA abundances in the mCRISPRi-v2 plasmid library (day 0). (Center and Right) Violin plots of the CD11b and GFP top/bottom log2 ratios for each gene, categorized into equally spaced bins of DepMap essentiality score.

B. Scatterplot of log2 ratio (L2FC) between the top and bottom 25% of the CD11b and CD14 screens in the THP-1 BRQ CRISPRi screen. Genes that are negative hits in either screen (FDR < 0.1 and L2FC < -0.5) are highlighted in blue and selected genes are labeled.
C. KEGG gene ontology analysis of 178 hits that are enriched in the lowest quartile of CD11b expression. Categories with an adjusted p-value of <0.05 are displayed.
D. KEGG gene ontology analysis of hits that are enriched in the highest quartile of CD11b (top) or CD14 (bottom) expression. Categories related to DNA replication are highlighted in red. For the CD11b screen, the most significant 10 categories by adjusted p-value are displayed, and for the CD14 screen, all categories with an adjusted p-value of <0.05 are displayed.
E. Gene scores for selected genes (from Figure 4J) are displayed for the CRISPRi screen in this study and the CRISPR KO study from Wang et al. (2021).
F. Median fluorescence intensity (MFI) of EdU incorporation during S phase in single-gene knockdown THP-1 cell lines treated with DMSO or 500 nM BRQ for 48 hours (n=3). (*) p < 0.05, (**) p < 0.005, (***) p < 0.0005, Student's two-sample t-test.Guide scores for selected DNA replication genes that were validated to increase CD11b expression in THP-1 cells following BRQ treatment.
G. Scatterplot showing percentage of CD11b+ cells in single-gene knockdown THP-1 cells treated with DMSO or 500 nM BRQ for 48 hours (n=3). Dashed line represents y=x.
H. Gene set enrichment analysis for RNA-seq data from THP-1 dCas9-KRAB cells transduced with sgMCM2, sgPOLD3, sgGOT2, and sgABCB7 compared to control sgNTC, using upregulated genes from PMA-treated THP-1 cells compared to untreated cells as the gene set of interest. NES, normalized enrichment score.
I. Heatmap showing RNA-seq data for selected macrophage, monocyte, and cell cycle-related genes from THP-1 dCas9-KRAB cells transduced with sgMCM2, sgPOLD3, sgGOT2, or sgABCB7 (n=3-4 per guide). Data displayed as log2 fold change from sgNTC-transduced cells.
J. Metascape enrichment for genes whose knockdowns lead to top 50 erythroid (top) or myeloid (bottom) scores.
K. Scatterplot of relative proliferation versus erythroid score in K562 cells from the Perturb-seq experiment. Replication- and nucleotide synthesis-related genes are highlighted in red. A linear regression (y-intercept=0) with a 95% confidence interval is plotted for replication-related and all genes. P-value was calculated by comparing the slope of the DNA replication gene set with the slopes calculated from 10,000 equally sized sets of randomly selected genes.
L. Schematic of sublibrary design and screen in THP-1 and K562 cells.
M. Scatterplots of CD11b (left) and CD14 (right) gene scores in DMSO versus BRQ treatment in the THP-1 sublibrary screen. Pearson correlations are displayed.

N. Scatterplot of CD11b versus CD14 score in the THP-1 sublibrary screen. Various categories of hits are colored as indicated.
O. (Top) Venn diagram of overlapping hits (L2FC > 0.5, FDR < 0.2) from the sublibrary screen in THP-1 and K562 cells whose knockdown increases CD11b, CD14, or CD235a expression, respectively. (Bottom) Hits related to DNA replication are highlighted.

**Figure S5. Relationships between replication stress-induced cell state changes and cell cycle phase, length, replication stress signaling, and DNA damage response.** Related to Figure 5.

A. (left) Relative scRNAseq expression of *Ung* (a G1 marker), *Rrm2* (S phase), and *Aurka* (G2/M phase), scaled from 0 to 1, for ER-Hoxa9 cells treated with 1 μM BRQ (top) or estrogen withdrawal (-E2, bottom) for 0, 12, 24, and 48 hours. Cells are plotted in UMAP space calculated from a principal components analysis conducted using only cell cycle-related genes. (rigth) Granulocyte score for BRQ-treated (top) or E2 withdrawn (bottom) cells treated at 0, 12, 24, and 48h, plotted in the same coordinates as on the left.
B. Immunoblots of ER-Hoxa9 cells following 16h of treatment with 1 μM BRQ, 20 nM of the ATR inhibitor VE-821 (ATRi), and various doses of the ATM inhibitor AZD0156 (ATMi) as indicated.
C. Immunoblots of THP-1 cells following 48h of treatment with DMSO (Veh), 1 μM APH, 500 nM BRQ, or 100 μM HU, with or without 500 nM AZ20 (ATRi). One representative blot is shown of two experiments. The arrowhead represents Chk1 that is singly phosphorylated at the S345 position by ATR. The asterisk represents a hyperphosphorylated CHEK1 species that occurs alongside DNA damage and is likely due to autophosphorylation as well as phosphorylation of serine 345 by ATM and other kinases.
D. Percentage of cells expressing CD11b (left) and CD14 (right) in THP-1 cells following 48h of treatment with THP-1 cells with DMSO, 1 μM APH, 500 nM BRQ, or 100 μM HU, with or without 500 nM AZ20 (ATRi) (n=3) (*) $p < 0.05$, (***) $p < 0.001$, (****) $p < 0.0001$, and adjusted p-values were calculated using a two-way ANOVA with the Šidák correction for multiple comparisons.
E. Relative expression (by qRT-PCR) of p21 (*Cdkn1a*) in ER-Hoxa9 cells treated with DMSO, 1μM BRQ, or 1 μM BRQ + 100 nM rabusertib (Chk1i) for 24 hours. Expression is normalized to *GAPDH* and displayed relative to DMSO (n=3). (*) $p < 0.05$, (**) $p < 0.01$, and p-values were calculated using unpaired, two-tailed Student's t tests with unequal variances.
F. Immunoblot of ER-Hoxa9 cells treated with DMSO, 1 μM BRQ, 50 μM HU, or the indicated doses of ROSC (roscovitine) for 24h. Similar results were seen at 8h.
G. Fork speed (kb/min) for ER-Hoxa9 cells treated with DMSO or 25 μM ROSC for 8h. (****) $p < 0.0001$, Mann-Whitney U test. At least 100 forks were analyzed for each condition.
H. EdU median fluorescence intensity (MFI) for ER-Hoxa9 cells treated with DMSO or 25 μM ROSC for 8h (n=3). P-value calculated by Student's t test.

I. Percentage of cells in each phase of the cell cycle for ER-Hoxa9 cells treated with DMSO or 25 µM ROSC for 8h (n=3). P-value calculated by two-way ANOVA.

J. G1 phase length (in hours) for ER-Hoxa9 Geminin-mCherry cells treated with DMSO or the indicated drugs, as measured by the length of time daughter cells were mCherry negative following a mitotic event. At least 100 cells were analyzed for each condition. (****) $p < 0.0001$, Mann-Whitney U test.

K. Fraction of G1 cells with at least one 53BP1 focus, determined by quantitative image-based cytometry. G1 cells were determined to be EdU-negative cells with 2N DNA content (DAPI). n=4 biological replicates, with at least 130 cells per replicate. P-values calculated with two-sample Student's t-test with unequal variances.

L. (left) Change in log2 counts per million (CPM; from RNA-seq) for p21 (*Cdkn1a*) in ER-Hoxa9 cells treated with 1 µM BRQ or E2 withdrawal for the indicated amount of time (n=2). (right) Change in log2 transcripts per million (TPM) for p21 (*CDKN1A*) in THP-1 cells treated with 500 nM BRQ or 1 mM HU for the indicated amount of time (n=3-4).

M. (left) Expression of p21 (*Cdkn1a*) in ER-Hoxa9 dCas9-KRAB-mCherry cells with p21 (sgCdkn1a) or mock (sgNTC) knockdown as assessed by qRT-PCR. Expression is normalized to *Gapdh* and displayed relative to sgNTC (n=3). (**) $p < 0.01$ (unpaired, two-tailed Student's t test with unequal variances). (right) Percentage of CD11b-expressing ER-Hoxa9 dCas9-KRAB-mCherry cells with p21 (sgCDKN1A) or mock (sgNTC) knockdown following treatment with vehicle, 1 µM BRQ, or 1 µM APH for 48 hours (n=3). (****) $p < 0.0001$ relative to sgNTC (two-way ANOVA with the Šidák correction for multiple comparisons).

N. (left) Expression of p21 (*CDKN1A*) in THP-1 dCas9-KRAB-mCherry cells with p21 (sgCDKN1A) or mock (sgNTC) knockdown as assessed by qRT-PCR. Expression is normalized to *GAPDH* and displayed relative to sgNTC (n=3). (**) $p < 0.01$ (unpaired, two-tailed Student's t test with unequal variances). (right) Percentage of CD11b-expressing THP-1 dCas9-KRAB-mCherry cells with p21 (sgCDKN1A) or mock (sgNTC) knockdown following treatment with vehicle, 500 nM BRQ, 100 µM HU, or 1 µM APH for 48 hours (n=3). (*) $p < 0.05$ relative to sgNTC (two-way ANOVA with the Šidák correction for multiple comparisons).

O. Percentage of viable and CD11b-expressing ER-Hoxa9 and THP-1 cells after 96 hours of treatment with the indicated DNA damaging agents (n=2).

P. Percentage of CD11b-expressing ER-Hoxa9, THP-1, and U937 cells 48 hours after gamma-irradiation with 0, 2, or 8 Gy (n=4).

Q. (Left) Median fluorescence intensity (MFI) for EdU incorporation in ER-Hoxa9 cells treated with DMSO, 1 µM brequinar (BRQ), 2.5 µM cisplatin (CIS), or 2.5 µg/ml neocarzinostatin (NCS) for 24 hours (n=3). (Right) Number of RPA foci per cell. (*) $p < 0.05$, (****) $p < 0.0001$, Mann-Whitney U test. 50-300 cells were quantified for each condition, and results from one representative experiment of two are displayed.

R. Representative immunofluorescence images of ER-Hoxa9 cells treated with DMSO or 2.5 μg/ml neocarzinostatin (NCS) for 24h and stained with DAPI (blue) and for RPA (yellow). Scale bar represents 10 μm.

S. Percentage of ER-Hoxa9 cells in each cell cycle phase after 24 hours of treatment with DMSO, 1 μM BRQ, 2.5 μM CIS, or 2.5 μg/ml NCS following 16h of culture in media with 100 or 2 ng/ml SCF (n=3).

T. Representative immunofluorescence images of cells described in (Q) stained with DAPI (blue) or for γH2AX (red) as indicated. Scale bar represents 10 μm.

U. Median γH2AX IF signal per cell for the treatments described in (Q) (n=3 biological replicates, 60-300 cells per replicate). For panels U and V, (*) $p < 0.05$, (**) $p < 0.01$, (***) $p < 0.001$, and (****) $p < 0.0001$, and adjusted p-values were calculated using a two-way ANOVA with the Šidák correction for multiple comparisons.

V. Median number of γH2AX foci per cell in the same samples described in (Q).

**Figure S6. Analysis of gene expression changes during replication stress.** Related to Figure 6.

A. Number of genes differentially expressed in ER-Hoxa9 cells (|LFC| > 1.5, adjusted p-value < 0.05) at each timepoint following BRQ treatment (red) or E2 withdrawal/Hoxa9 OFF (blue) for the indicated amount of time (n=2).

B. Gene ontology categories for genes that are upregulated in both -E2 and BRQ at 48h.

C. Gene ontology categories for genes that are downregulated in -E2 but not in BRQ at 48h.

D. Metaplot (top) and heatmap (bottom) of ER-Hoxa9 CUT&RUN signal after DMSO, BRQ 24h and -E2 24h treatment at peaks identified using MACS2 from the DMSO condition. Signal from 1 kilobase downstream to 1 kilobase upstream of the center of each peak is shown.

E. Number of ER-Hoxa9 CUT&RUN peaks identified using MACS2 after DMSO, BRQ 24h, and -E2 24h treatment.

F. Examples of ER-Hoxa9 CUT&RUN signal at the *Ebf1* and *Meis1* loci.

G. Identification of putative targets of ER-Hoxa9 by intersecting bound genes with genes that are downregulated only after 48h of E2 withdrawal but not after 48h of BRQ treatment. P-value of the overlap (compared to the number expected if genes were randomly selected) was calculated using Fisher's exact test.

H. Gene ontology categories for genes that are downregulated or upregulated in THP-1 cells following treatment with 500 nM BRQ, 100 μM HU, or 100 nM PMA for 48 hours.

I. Percentage of CD235a-positive K562 dCas9-KRAB cells transduced with a negative control guide (sgNTC) or a guide targeting the *BCR* promoter (sgBCR) for 5 days.

J. Log2 fold change in counts per million for *BCR* and *ABL1* transcripts K562 dCas9-KRAB cells expressing sgBCR compared to control sgNTC. (*) $p < 0.05$, (**) $p < 0.005$ (Student's t-test).
K. Gene ontology analysis (MSigDB Hallmarks) for genes that are shared or differentially regulated between 250 nM BRQ treatment (72h) and *BCR* knockdown in K562 dCas9-KRAB cells.
L. Gene set enrichment analysis (GSEA) for RNA-seq of sgBCR cells compared to sgNTC (left) or BRQ-treated cells compared to DMSO (right). The gene sets used are the top 500 upregulated or downregulated genes in P5/P6 erythroblasts compared to hematopoietic stem cells (HSCs) and multipotent progenitors (MPPs) from a primary human erythroid differentiation culture system (Ludwig et al. 2019).

**Figure S7. Analysis of RNA polymerase, H3K27ac, and ATAC-seq changes during replication stress.** Related to Figure 6.

A. Log2 fold change in gene body RNA polymerase II occupancy (CUT&RUN) for genes downregulated, upregulated, or not statistically significantly changed (NS) after 24 hours of E2 withdrawal in ER-Hoxa9 cells. (****) $p < 0.0001$, Mann-Whitney U test.
B. (left) Number and percentage of peaks with significantly increased or decreased H3K27ac signal after 8 or 24 hours of BRQ treatment or E2 withdrawal in ER-Hoxa9 cells. (right) Number of peaks with significantly altered H3K27ac signal within each genomic region after 24 hours of BRQ treatment or E2 withdrawal, with the log2 observed/expected ratio (compared to all peaks) denoted with the color map.
C. Same as (B) for THP-1 cells treated with BRQ or PMA for 24 hours.
D. Log2 fold change in gene expression (RNAseq) for genes within 100kb of distal regulatory elements ("enhancers") whose H3K27ac signal is downregulated, upregulated, or not statistically significantly changed (NS) after 24 hours of E2 withdrawal in ER-Hoxa9 or PMA treatment in THP-1 cells. (****) $p < 0.0001$, Mann-Whitney U test.
E. Log2 fold change in promoter H3K27ac signal (CUT&RUN) for genes downregulated, upregulated, or not statistically significantly changed (NS) after 24 hours of E2 withdrawal in ER-Hoxa9 or PMA treatment in THP-1 cells. (****) $p < 0.0001$, Mann-Whitney U test.
F. Schematic of CUT&RUN experiment in THP-1 PIP-FUCCI cells.
G. FACS plots demonstrating sorting strategy. mCherry versus mVenus signal was used to gate G1, S, and G2/M cells. S phase cells were then gated by DNA content (DyeCycle Violet) such that the top third of cells was sorted.
H. FACS plots of EdU incorporation versus DAPI for all cells (top) or early S phase cells (bottom) immediately post-sort, eight hours after vehicle treatment, or eight hours after treatment with 100 μM HU. S phase cells (EdU-positive) are quantified.

I. (left) Log2 change in gene expression (RNAseq) between early S phase cells treated with HU for 8 hours and immediately post-sort, for genes downregulated, upregulated, or not statistically significantly changed after 24 hours of BRQ treatment in THP-1 cells. (right) Log2 change in H3K27ac CUT&RUN signal between early S phase cells treated with HU for 8 hours and immediately post-sort, for peaks whose H3K27ac signal is downregulated, upregulated, or not statistically significantly changed after 24 hours of BRQ treatment in THP-1 cells. (****) $p < 0.0001$, Mann-Whitney U test.

J. IGV genome browser tracks of H3K27ac or IgG CUT&RUN signal at exemplar loci for asynchronous cells treated with DMSO or BRQ for 24 hours, or early S phase cells immediately post-sort or after HU treatment for 8 hours.

K. (left) Density plot of log2 fold change in ATAC-seq signal for 83,280 peaks after 24 and 48 hours of BRQ or -E2 treatment in ER-Hoxa9 cells compared to DMSO. (right) Number of peaks with significantly altered H3K27ac signal within each genomic region after 24 hours of BRQ treatment or E2 withdrawal, with the log2 observed/expected ratio (compared to all peaks) denoted with the color map.

L. Same as (K), but for 110,278 peaks in THP-1 cells treated with BRQ or PMA for 24 or 48 hours.

M. Top 30 motifs by variability and their accessibility scores from ChromVAR, clustered by k-means analysis, at timepoints after BRQ and -E2 treatment in ER-Hoxa9 cells.

N. Top 30 motifs by variability and their accessibility scores from ChromVAR, clustered by k-means analysis, at timepoints after BRQ and PMA treatment in THP-1 cells.

O. HOMER analysis of *de novo* motifs enriched at peaks differentially marked by H3K27ac after 24h of BRQ treatment or E2 withdrawal in ER-Hoxa9 cells. The fold enrichment and adjusted P-value of the motif compared to all distal regulatory regions are denoted.

P. Same as (O), but for THP-1 cells treated with BRQ or PMA for 24 hours.

**Figure S8. Preferential activation of primed, lineage-specific gene expression programs during replication stress.** Related to Figure 7.

A. IGV browser tracks of ATAC-seq and H3K27ac signal for two genes (*ST3GAL6* and *CCL2*) after 24h or DMSO, BRQ, or PMA treatment in THP-1 cells, with RNAseq log2 fold changes indicated.

B. (top) Distribution of expression changes for shared upregulated genes. (bottom) Distribution of the proportion of peaks that gain accessibility for all shared upregulated genes. Data after 24h of BRQ or -E2 treatment in ER-Hoxa9 cells (left) or BRQ or PMA treatment in THP-1 cells (right) are displayed. P-values were calculated using the Mann-Whitney U test.

C. Fraction of associated loci with increased accessibility for shared upregulated genes after 24h of BRQ versus -E2 treatment in ER-Hoxa9 cells (top), or 24h of BRQ versus PMA treatment in THP-1 cells (bottom).

D. Heatmaps of open chromatin regions (OCRs) with significantly increased H3K27ac after BRQ treatment or PMA treatment in THP-1 cells, displaying H3K27ac or ATAC-seq signal before and after treatment in a ±2kb window surrounding the center of the OCR.
E. Log2 fold change in mRNA expression of PU.1 at each timepoint following BRQ treatment in ER-Hoxa9 cells (n=2).
F. Log2 fold change in expression of PU.1 target genes at each timepoint following BRQ treatment in ER-Hoxa9 cells (n=2). PU.1 target genes (n=110) were obtained from (Turkistany et al., 2011) (*) $p < 0.05$, one-sample Student's t-test.
G. Immunoblot of ER-Hoxa9 cells following treatment with 2 μM BRQ for the indicated timepoints.
H. Western blot analysis of PU.1 (Spi1) expression in ER-Hoxa9 dCas9-KRAB cells expressing control (sgNTC) or PU.1 knockdown (sgSpi1) guides.
I. Top 30 motifs by variability and their accessibility scores from ChromVAR, clustered by k-means analysis, in ER-Hoxa9 dCas9-KRAB cells expressing sgNTC or sgSpi1 guides.
J. Gene ontology categories (from PanglaoDB) for cell types whose marker genes are upregulated by BRQ treatment specifically in the ER-Hoxa9 dCas9-KRAB cells expressing sgNTC or sgSpi1 guides.
K. ATAC-seq tracks in sgNTC and sgSpi1 ER-Hoxa9 dCas9-KRABH cells for representative genes not upregulated (*Ms4a3*, *Tyrobp*) or upregulated (*Prss34*, *Cd247*) upon BRQ treatment in sgSpi1 cells.
L. Log2 counts per million for *GATA1* expression in K562 dCas9-KRAB cells expressing control sgNTC or GATA1 knockdown (sgGATA1) guides and treated with DMSO or 250 nM BRQ for 72 hours. (*) $p < 0.05$, and p-values were calculated using a Student's t-test.
M. Gene ontology categories (from Human Gene Atlas) for cell types whose marker genes are upregulated by BRQ treatment specifically in K562 dCas9-KRAB cells expressing sgNTC or sgGATA1 guides.
